# Global biogeography of geckos (Squamata: Gekkota) reveals Old-World origins and region-specific heterogeneity in diversification

**DOI:** 10.64898/2026.10.21.760521

**Authors:** Yuanxing Liao, Phillip L. Skipwith

## Abstract

**Aim:** Investigating the biogeography and diversification dynamics of global radiations to understand how Earth history has influenced lineage proliferation has long been a goal of macroevolutionary biology. Gecko lizards, being an ancient and globally distributed hyperdiverse lineage, are an excellent system to elucidate how certain regions and events have shaped today’s observed diversity. In this study, we attempt to identify: 1) The heterogeneity in diversification rate across geckos; 2) the ancestral region of geckos; 3) if diversification rate is tied to biogeographic region; and 4) potential mass extinction events in geckos.

Location

Global.

Taxon

Geckos (Squamata: Gekkota).

**Methods:** We generated a massive supermatrix of geckos from Sanger data and, after combination with phylogenomic data, estimate the most comprehensive phylogenetic estimate of geckos to date with 1,783 species. We performed fossil informed ancestral area reconstructions to evaluate the impact of fossils. We performed both state-dependent and state-agnostic diversification analyses to access diversification dynamics. Lastly, we tested for clade-wide mass extinction events.

**Results:** Our divergence date estimates suggested a Jurassic origin for pan-Gekkota and a Cretaceous origin for crown Gekkota. We recovered an Old-World origin for crown Gekkota with high uncertainty and an African origin for the families Phyllodactylidae, Sphaerodactylidae, and Gekkonidae with high confidence. Our results suggest differential diversification dynamics between pygopodoid (Australasian) and gekkonoid lineages (cosmopolitan), with the latter experiencing overall higher rates and more heterogeneity. Our results further suggest Africa being a biodiversity “refugia” while Eurasia acted as a “diversification hotspot” for geckos. Lastly, we find no evidence that crown geckos experienced any mass extinctions despite surviving the K-Pg event and major climatic oscillations during the Cenozoic.

**Main conclusions:** Our study provides an important global view for the biogeographic and diversification patterns of a morphologically and ecologically distinct lineage of lizards, setting the stage for studies placing clade-specific histories in a global context.

## Introduction

Macroevolutionary biology has a long history of studies investigating the biogeography of global radiations, particularly in the context of speciation and trait diversification dynamics. Diverse lineages with large distributions lend themselves well to understanding the historical factors that have driven asymmetry in species diversity and the tempo of species accumulation between and within clades. With more than 12,200 species worldwide, lizards and snakes (Sauropsida: Squamata) constitute major components in vertebrate communities, especially when considering ecological importance and species diversity. Squamate morphological diversity vastly eclipses that seen in the next two most diverse amniote lineages, birds (∼10,000 species) and mammals (∼6,000 species). However, the vast majority of terrestrial, limbed squamates (‘lizards’) are diurnal and most clades have restricted geographic distributions. Geckos (Squamata: Gekkota) represent the only widespread lineage of predominately nocturnal lizards, occupying functional niches inaccessible to other squamates. Further, gekkotans are incredibly diverse with more than 2,400 species distributed globally, albeit with strong circumtropical and eastern hemisphere biases in species diversity (Meiri 2019). The seven extant families are divided between two primary clades, the Pygopodoidea (Diplodactylidae, Carphodactylidae, and Pygopodidae) and Gekkonoidea (Gekkonidae, Phyllodactylidae, Sphaerodactylidae, and Eublepharidae). In addition to being a largely nocturnal clade nested within an otherwise diurnal lineage, geckos possess a number of notable traits including adhesive digital scansors (toepads) and olfactory chemoreception (as opposed to vomeronasal chemoreception, which is more widespread in squamates) (Schwenk 1993, Rehorek et al. 2000, Gamble et al. 2012, 2017). Some authors have linked specific gecko traits with the clade’s cosmopolitan distribution suggesting that some, such as toepads and desiccation-resistant eggs, have made some geckos exceptionally proficient at trans-oceanic dispersal (Andrews 2012, Gamble et al. 2012, Choi et al. 2018, Russell and Gamble 2019, Kukla et al. 2025). In line with these hypotheses explaining their apparent success, Meiri (2019) determined that geckos are more likely to be insular than other squamates with more than a third of species being insular endemics. This apparent predilection for overwater dispersal has resulted in numerous insular lineages that likely represent candidate adaptive radiations where speciation is tied to ecological divergence due to open niche space. These include ecomorphologically diverse and speciose lineages in Sundaland (*Cyrtodactylus*, *Gehyra*, and *Gekko/Lepidodactylus*: Gekkonidae), Australasia (Pygopodoidea), Indian Ocean islands (*Phelsuma*, *Uroplatus*, *Cnemaspis*, and *Hemidactylus*: Gekkonidae), and the Greater Antilles (*Sphaerodactylus*: Sphaerodactylidae) (Yoder and Nowak 2006, Greenbaum et al. 2007, Gamble et al. 2008a, Raxworthy et al. 2008, Heinicke et al. 2011, Oliver et al. 2012, Wood et al. 2012b, Sistrom et al. 2013, Oliver et al. 2014, Garcia-Porta et al. 2016, Skipwith et al. 2016, Ashman et al. 2018, Moritz et al. 2018, Tallowin et al. 2018, Daza et al. 2019, Oliver et al. 2019, Skipwith et al. 2019, Grismer et al. 2022, Skipwith and Oliver 2023).

While geckos are distributed across all continents (except Antarctica), there is clear asymmetry between regions. Species diversity is highest in Southeast Asia, Australasia, and sub-Saharan Africa. Diversity is much lower in the Americas, Europe, and northern Asia (Uetz et al. 2020). Further, this distribution appears to have very strong phylogenetic signal with one of the major clades, the pygopodoids, being entirely restricted to Australasia. Almost 90% of gecko diversity lies within the sister taxon of pygopodoids, the gekkonoids, which are most diverse outside of Australasia. With the advent of molecular phylogenetics, a number of authors have proposed that this pattern represents a split between eastern (pygopodoids) and western (gekkonoids) Gondwanan lineages (Han et al. 2004, Feng et al. 2007, Oliver and Sanders 2009). Further, previous studies leveraging molecular data have proposed that vicariance, trans-oceanic dispersal, and over-land dispersal explain the colonization of the New World by gekkonoids during the late Cretaceous and throughout the Cenozoic (Gamble et al. 2008a, b, Gamble et al. 2011). However, no study to date has implemented fossil occurrence data alongside the newest comparative models to elucidate the origins of the global gecko radiation. Based off the distribution of extant lineages, it is generally assumed that crown gekkotans are of southerly Laurasian or even Gondwanan origin with multiple invasions of Laurasia (Han et al. 2004, Meyer et al. 2023). A Laurasian origin for pan-Gekkota (Gekkota + stem lineages) and crown Gekkota was proposed by Meyer et al. (2023) after incorporating stem fossils alongside extant lineages. While insightful, this study implemented very sparse sampling of crown geckos and used a morphological phylogenetic hypothesis with numerous relationships not recovered in any molecular study (Burbrink et al. 2020, Singhal et al. 2021). In this study, we implement birth-death diversification and biogeographically informed state-dependent models to infer the ancestral areas of origin of pan-Gekkota and crown Gekkota on a combined phylogenomic and supermatrix dataset, sampling ∼70% of extant gecko species and select stem taxa.

Ecological opportunity and novel adaptations have long been associated with the success and/or ultimate failure of clade survival on a geologic time scale. However, these variables are linked with speciation and ecological differentiation and many studies fail to investigate factors that catastrophically impact species diversity through extinction. Geckos writ large are an ideal system to investigate the impact of mass extinction in shaping the tempo of diversification. Being an ancient, southerly lineage, geckos have survived tumultuous periods in the planet’s past. Molecular estimates almost entirely agree that crown geckos would have survived the Cretaceous-Paleogene (K-Pg) extinction event (66 mya) that led to the extinction of countless taxa, including all non-avian dinosaurs and a number of squamate lineages (polyglyphanodonts, mosasaurs, paramacellodids, etc.). The K-Pg extinction event, while fundamental to the evolutionary history of life on Earth, was a single instantaneous event (the impact) with long term cataclysmic consequences (global fallout). In contrast, the Eocene-Oligocene (∼34 mya) boundary (EoB; also called the Eocene-Oligocene cooling event or transition) saw a protracted drop in global temperatures that is associated with mass extinctions followed by turnover and wholesale environmental changes such as the expansion of grasslands (Zanazzi et al. 2007, Stadler 2011, Sun et al. 2014, Godfrey et al. 2020). Patterns of mass extinction with subsequent faunal turnover coinciding with the EoB has been proposed for regional squamate assemblages, with new lineages colonizing more xeric environs (Smith 2006, Kirchhof et al. 2021). However, evidence for this pattern in geckos is mixed, with Brennan and Oliver (2017) and Skipwith et al. (2019) finding strong evidence for and no support for mass extinction and turnover in pygopodoids coinciding with the EoB, respectively.

Given that geckos are an ancient, hyperdiverse lizard lineage, numerous studies have focused on estimating the diversification dynamics of individual clades such as the African *Pachydactylus* group (Heinicke et al. 2017), Malagasy *Phelsuma* (Harmon et al. 2008), Australasian pygopodoids (Garcia-Porta and Ord 2013, Skipwith et al. 2016, Brennan and Oliver 2017, Skipwith et al. 2019, Skipwith and Oliver 2023), and Southeast Asian gekkonids (Wood et al. 2012b, Oliver et al. 2014, Tallowin et al. 2018, Oaks et al. 2019). However, none have attempted to generalize biogeographic history and diversification patterns across all of Gekkota. Here, we use the most taxonomically complete gecko phylogeny to date to: 1) determine if heterogeneity in diversification rate is present across geckos, 2) identify the ancestral region of origin of geckos, 3) infer if observed heterogeneity is tied to particular zoogeographic regions, and 4) if major earth history events (i.e.- K-Pg, EoB, etc.) have driven wholesale changes in gecko diversification. These areas of inquest are pertinent as comparative analyses of globally distributed lineages have highlighted that specific regions have acted as centers of origin or refugia. For instance, Moyle et al. (2016) suggested that oscine songbirds originated in Australasia, but it was the later formation of Wallacea that led to global success of the clade. We demonstrate that several gecko clades have experienced bursts of diversification and identify the Old World as the likely center of origin for Gekkota.

## Materials and Methods

### Taxon sampling

We sampled as many gecko species as possible for nucleotide data to produce the most taxonomically complete phylogeny of geckos to date using publicly available Sanger sequence data. Ultimately, sequence data for 19 additional loci (Table S1) were collected from GenBank as these were commonly used loci in gecko systematics and we combined our matrix with the supermatrix of Zheng and Wiens (2016) (supplementary text). We included sequences for 6 outgroup taxa: *Sphenodon punctatus*, *Dibamus bourreti*, *Lacerta agilis*, *Anolis carolinensis*, *Heloderma suspectum*, and *Anguis fragilis*. Our initial dataset contained 71 genes (Table S1) for 1759 species.

### Phylogenetic analyses

To align protein coding loci, we first adjusted the reading frame using Coding_Translation_Tests.py module in superCRUNCH 1.3.2 (Portik and Wiens 2020). We then used the Align.py module in superCRUNCH with *-a macse* flag to run MACSE 2.07 (Ranwez et al. 2018), with both *--pass_fail* and *--accurate* flags turned on. To align non-protein coding genes such as 12S and 16S, we used TaxonomyAlign v1.0 (Portik et al. 2023). We used genera as taxonomic groups to define groupings in the mapping file, and we used E-INS-i as the alignment algorithm. Individual alignments were then trimmed on both ends by eye to reduce missing data and to remove poorly aligned flanking regions. They were then concatenated into a single alignment file using Concatenator (Vences et al. 2022). See Figure S1 for a infographic flowchart detailing phylogenetic analyses.

Protein coding loci were then partitioned by their codon position (1^st^, 2^nd^ and 3^rd^) while 16S and 12S were treated as a single partition. The best partition scheme was found using ModelFinder (Kalyaanamoorthy et al. 2017). This was done in IQ-TREE 2.2.2.6 (Minh et al. 2020) with the greedy algorithm by subsequently merging partitions (Lanfear et al. 2012). Next, we ran maximum likelihood (ML) tree search with 1000 ultrafast bootstrap replicates (UFBoot, *-B 1000*) (Hoang et al. 2018). The tree search analysis was repeated three times. We identified and removed rogue taxa using RogueNaRok v1.0.1 (Aberer et al. 2013) and we reran each tree search analyses again. The tree with the maximum likelihood was then selected and retained for downstream analyses.

Initially, the supermatrix data set had 95.5% missing data. To account for the effects of missing data, we progressively trimmed our supermatrix down to reduce the missing data by keeping the first 20, 5, 4, and 3 genes with the least missing data and applied the above tree search procedure on these four datasets, repeating each three times (details in supplementary text). Except for the 20-gene dataset, all three other trimmed datasets have fewer species than the 71-gene dataset.

### Divergence dating

We used the 20-gene ML topology and alignment to estimate divergence times. The motivation here was to maintain maximum species diversity while moderating the amount of missing data and computational load. There were initially three fossil calibration points chosen following previous studies (Skipwith et al. 2016, Skipwith et al. 2019), but with different prior distributions (Table 1). For better comparison with published phylogenies, we included three secondary calibration points that were collected from timetree.org (Kumar et al. 2022). To try to reconcile the incongruences of time estimation between nuclear and mitochondrial loci (supplementary text), we picked an additional 12 secondary calibration points based on our own nuclear-loci-only dating estimate as well as previously published studies. We conducted the divergence dating analyses using BEAST 2.7.7 (Bouckaert et al. 2019). The concatenated alignment was divided into two partitions, one with only mitochondrial loci and the other with only nuclear loci. We estimated molecular evolution model for each partition in IQ-TREE 2 (Minh et al. 2020). The molecular clock model was estimated separately for each partition using the optimized relaxed clock (ORC) model (Douglas et al. 2021) with mean clock rate set to 1 initially before being estimated empirically. Tree topology was fixed to the ML topology. The burn-in was set to 5,000,000 iterations, and it was sampled every 1000 iterations until it reached the chain length of at least 100,000,000. This analysis was run with five independent chains in parallel. Convergence was checked in the program Tracer v1.7.2 (Rambaut et al. 2018) to make sure sufficient mixing of every parameter. These five chains were then combined to make sure that the effective sample size (ESS) for each relevant parameter should be greater than 200 for sufficient sampling of the posterior. We then randomly sampled 12,000 trees from the last 40% of trees of each run and then combined so there were total of 60,000 tree samples. The maximum clade credibility (MCC) tree was summarized using TreeAnnotator (Heled and Bouckaert 2013).

**Table 1.**
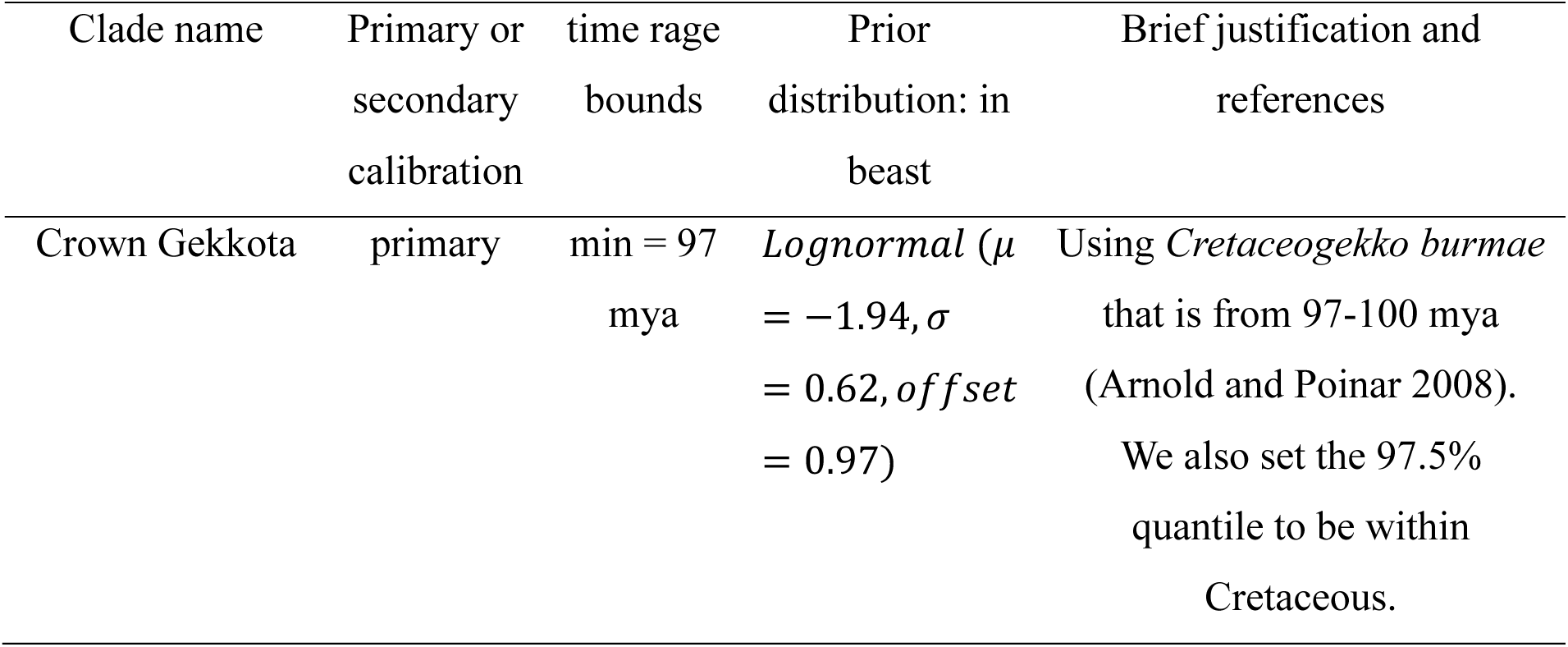
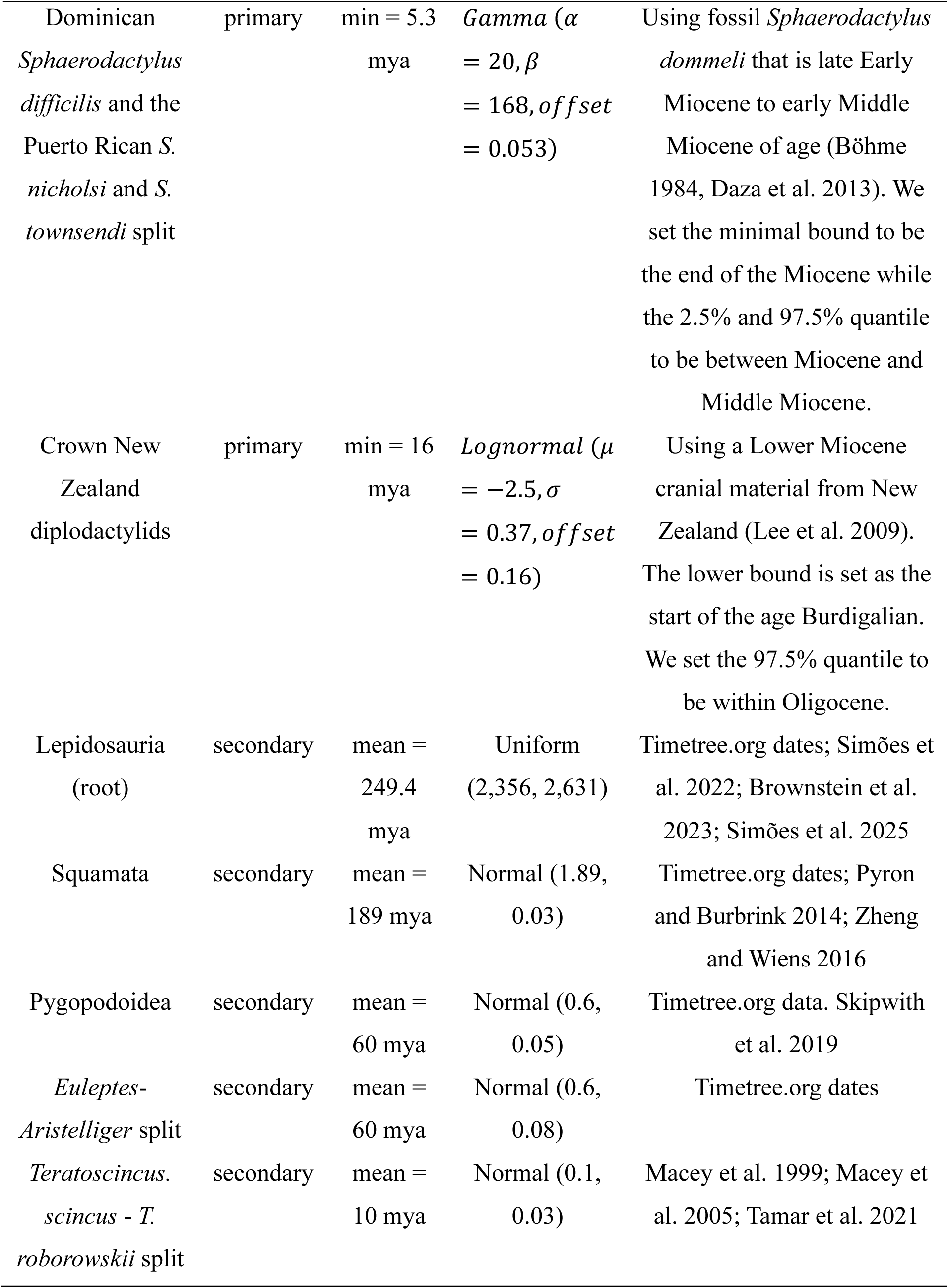
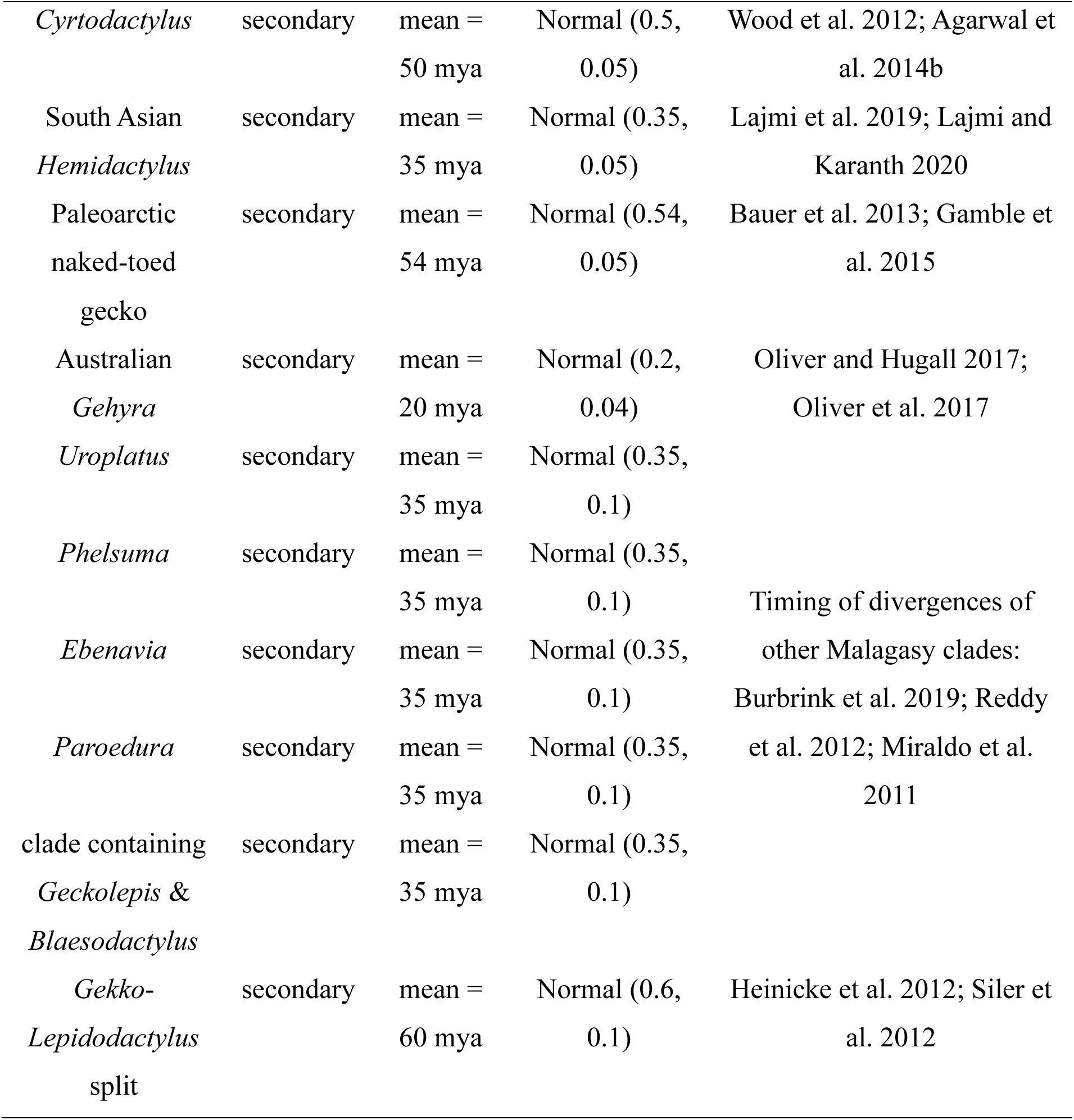
The node of the clade, type of the calibration, and the calibration prior distribution used in the BEAST2 analyses. We picked the last 12 secondary priors to reconcile the incongruence of time estimations between nuclear and mitochondrial loci (Supplementary text). Detailed justification and references can be found in supplementary text.

### Combining Phylogenies and Rationale

The Sanger data available for numerous gecko groups are limited to a handful of loci with few lineages having phylogenomic-scale sequences yielding fully resolved, well-supported trees. Therefore, we used the abovementioned Sanger data for phylogenetic reconstruction of the entirety of Gekkota with the ultimate goal of marrying the Sanger data mined for gekkonoids here with the phylogenomic estimate for pygopodoids from Skipwith et al. (2019). Under this paradigm, all pygopodoids included in tree estimation using our Sanger supermatrices were pruned and then we grafted the Maximum Clade Credibility (MCC) gekkonoid supermatrix toplogy to the UCE MCC pygopodoid tree from Skipwith et al. (2019) using the bind.tree function in package APE 2.8-1 in R 4.4.3 (Paradis et al. 2004, R Core Team 2025), retaining the crown age of our full supermatrix tree. The same process was repeated for 10 random trees from the posteriors of BEAST 2 in this study and MCMCtree from Skipwith et al. (2019). We opted to replace the pygopodoid portion of the tree because the UCE topology was better supported than its Sanger counterpart and had significantly better taxonomic sampling (adding an additional 30 species). To be clear, these two trees were not inferred simultaneously from a single alignment due to unavoidable consequences of missing data amongst gekkonoids. Further, all comparative analyses from this point on in the manuscript used the grafted UCE-Supermatrix tree/s, not the intact Sanger supermatix topology (Figure S1).

### Diversification Dynamics

To assess the general diversification dynamics of geckos globally, we implemented multiple birth-death lineage through time (LTT) models on our grafted UCE-supermatrix MCC tree. We first tested for large rate shifts in diversification using a speciation-extinction analysis Bayesian in Analysis of Macroevolutionary Mixtures (BAMM) (Rabosky 2014). Speciation-extinction analyses were run for 100,000,000 generations and prior sensitivity was assessed by rerunning the analysis with the *expectedNumberOfShifts* prior set to 2 and 3. We opted for *expectedNumberOfShifts* > 1 due to the intermediate size of our tree (1,783 tips). BAMM analyses were summarized and visualized in BAMMtools (Rabosky et al. 2014). To account for incomplete taxon sampling, all BAMM analyses used lineage-specific taxon sampling fractions using genus-level species counts (average genus sampling fraction frequency = 83%). The most probable shift scenario was identified using the maximum *a posteriori* probability (MAP) shift configuration.

While BAMM is relatively robust at identifying large shifts associated with cladogenetic events, it may miss or misidentify small shifts occurring throughout a phylogeny (Maliet et al. 2019). Thus, we estimated diversification rates on branches using the ClaDS2 (constant turnover rate) model in RPANDA (Maliet et al. 2019). ClaDS analyses were run six times, each with 200,000 iterations and a thinning of 50. To identify shifts in speciation rate we used the hypothesis-free ‘search.shift’ function in RRphylo (Castiglione et al. 2018) on the branch rates estimated by ClaDS2 following the implementation used by Pavón-Vázquez et al. (2022). The search.shift algorithm automatically assesses the deviation in rates of individual clades from the whole tree. Because clades likely inherit the diversification patterns of the parental lineage, search.shift assesses each clade individually and only selects those subtending nodes that deviate the most from the background rate for the entire tree. When the probability of a shift was p > 0.975, shifts were inferred to be significantly higher than the background whereas p < 0.025 suggest significantly lower rates when α = 0.05.

In addition to inferring lineage-specific diversification dynamics, we explicitly estimated the timing and magnitude of shifts in possible clade-wide mass extinction events using the reversible-jump MCMC (rjMCMC) CoMET method (May et al. 2016) implemented in TESS (Höhna et al. 2016b). The rjMCMC implementation in CoMET simulates all possible episodically variable birth-death processes from an otherwise infinite array of possible model configurations while simultaneously estimating mass extinction events. We ran CoMET on our grafted tree from above for 200,000 iterations, 10% burnin, and parameters pertaining to the calculation of rate shifts and mass extinction events being estimated empirically from the data. Confidence in the presence of mass extinction events was assessed using Bayes factors, following the significance thresholds suggested by (Kass and Raftery 1995). Here, negative support for the alternative model (H_1_) was considered when BF < 1, weak support when BF = 1 to 3.2, substantial support when BF = 3.2 to 10, strong support when BF = 10 to 100, and unambiguous support when BF > 100. We assessed prior sensitivity by running additional CoMET analyses with the parameter *numExpectedRateChanges* and *numExpectedMassExtinctions* each set to 1, 2, and 3.

### Biogeographic Reconstruction

We divided the distribution of geckos into nine geographic regions based on paleogeographic maps (Scotese 2021). Details about the division of these regions and the justifications can be found in supplementary text. These nine regions are as follows:

A) Africa-Arabia – includes the African continent, Arabian Peninsula, and regions of Iran and Turkey that is south of the Zagros Mountain ranges.
B) Madagascar – includes Madagascar island as well as Réunion and Mauritius islands.
C) Eurasia - from Europe on the west to Japan on the east and Peninsular Malaysia on the south.
D) South Asia - includes India, Pakistan, Bangladesh, Nepal, Bhutan and Sri Lanka.
E) Sundaland - includes Borneo, Sumatra, and other Sunda Shelf islands, eastwards until the Lydekker line. For simplicity, this also includes the Philippines.
F) Australasia - eastwards of the Lydekker’s line, including New Guinea, Australia, New Zealand, New Caledonia and various islands and island nations in South Pacific Ocean, including Micronesia.
G) Caribbean – includes Greater Antilles, Lesser Antilles and the Bahamas
H) North-Central America – from USA, through Mexico, to Panama. This includes islands that are close to Central America geologically, such as the Bay islands of Honduras.
I) South America - from Colombia, Venezuela to Argentina in the south. This includes islands or island nations that are close to South America geographically, including Leeward Antilles island nations such as Aruba and Curaçao, and Trinidad and Tobago.

Biogeographic analyses and ancestral range reconstructions were done using BioGeoBEARS 1.1.3 (Matzke 2013) in R 4.4.3 (R Core Team 2025). We used 10 grafted UCE-Supermatrix trees described above to account for uncertainties in divergence time estimates. Time-stratified analyses were performed on each tree with each time slice having different matrices to modify parameters (dispersal rate for *d* and the dispersal weight for *j* specifically) based on specific modifiers (Table 4). These modifiers included distance matrices where physical distance between regions affects direct dispersal probabilities with longer distances making direct dispersal less likely. These distances were measured on paleogeographic maps (Scotese 2021) using GPlates 2.5.0 software (Müller et al. 2018) (Table S5). We named this the “Distance Hypothesis”. We implemented additional modifier matrices in the form of manual dispersal multiplier matrices. For adjacent but not connected regions, the dispersal multiplier was set to 0.5 as “less likely” to dispersal, and for non-adjacent regions, the multiplier was set to 0.001 as “unlikely” to disperse, since they are more likely to disperse to the adjacent island first (more details in supplementary text). We named this the “Position Hypothesis”. Finally, we fitted dispersal-extinction-cladogenesis (DEC) models (Ree and Smith 2008) with founder-event speciation (*+j*, Matzke 2014), with the distance hypothesis (*+x*, Van Dam and Matzke 2016), with position hypothesis (*+w*), or with any combination of them. We also fitted DEC and DEC+*j* models without time-stratification as null hypotheses (More details in supplementary text). Overall, 8 models were fitted and compared using AICc (Hurvich and Tsai 1993; Table S6). We performed 100 biogeographic stochastic mappings (BSM, Dupin et al. 2017) on each tree using the runBSM function within BioGeoBEARS package. To determine the input parameters of BSM, we used a model averaging approach instead of choosing the most supported model.

We incorporated fossil taxa in our ancestral area reconstruction analyses using a method inspired by Yan et al. (2021). For stem gecko fossil lineages, we sequentially attached *Hoburogekko suchanovi*, *Cretaceogekko burmae*, and *Gobekko creatacicus* outside of the crown clade. For fossil lineages that could be confidently assigned to extant genera, we grafted them to the most inclusive clades to which they could be assigned (More details in the supplementary text). All of the above procedures were done analytically in R using the bind.tip function in phytools package 2.5-2 (Revell 2012). To compare with the biogeographic hypotheses without fossils, identical sets of 8 geographic models were tested (DEC, *+j*, *+x*, *+w*, and any combination) using the same ten grafted phylogenetic trees.

### Biogeography and Diversification

To explore the role of biogeography on diversification rate within Gekkota, we implemented the multistate hidden-state state-dependent speciation and extinction models (MuHiSSE) in RevBayes v1.3.1 (Höhna et al. 2016a, Nakov et al. 2019). This model allows each region and its hidden state to have its own associated speciation rate and extinction rate. The regions codes used in this analysis were the same as the BioGeoBEARS analyses, and we used the grafted UCE-Supermatrix MCC tree for this analysis. Direct transitions between regions are allowed only in certain region pairs and in certain directions to reflect prior geologic knowledge (Table S10, the RevBayes script is available in the supplementary material). In situations where both forward and backward transitions between regions are allowed, the rates of forward and backward transitions are estimated independently. Due to the large number of states and state combinations, as well as the complexity of the model itself, only anagenetic events (e.g. dispersal without speciation and extirpation) were considered. For the same reason, the maximum number of areas that each lineage can occupy is 1. This means the states that included two biogeographic regions were coded as ambiguous states for which the likelihood would be integrated over all the possible states on the tips. The ancestral states were inferred with the joint conditional sampling implemented in the RevBayes function mnJointConditionalAncestralState(). To speed up computation, we ran this analysis with 20 independent chains. We did not set a fixed percentage of samples as burn-in for each run because the generations took to reach the equilibrium state were variable. Instead, we discarded samples as burn-in based on the trace plot in Tracer v1.7.2 (Rambaut et al. 2018). We then combined the log files to make sure every parameter had an ESS greater than 200. To determine if two posterior distributions were significantly different, we obtained a new distribution by subtracting values in each iteration from the two posterior distributions in comparison. If the 95% quantile of this new distribution included 0, then they were not significantly different, and vice versa.

## Results

### Supermatrix Topolgy

The ML topology of the 71-gene and 20-gene datasets are largely similar, so only the 20-gene dataset is described here unless otherwise noted. The effects of missing data was minimal with the majority of groupings consistent across datasets (Figure S2, supplementary material). Alignment statistics and topologies from all other datasets are included in the supplementary material (Table S1). All gecko families were recovered as monophyletic and they split into two major clades: Pygopodoidea, which is restricted to Australasia, and Gekkonoidea, which comprises the rest of the families which have a cosmopolitan distribution. Within Pygopodoidae, family Carphodactylidae is sister to Pygopodidae, and together, they are sister to Diplodactylidae (Figure 1). Within Gekkonoidea, Gekkonidae is sister to Phyllodactylidae, which together are sister to Sphaerodactylidae to the exclusion of Eublepharidae. These family-level relationships are all highly supported with bootstrap values (BS) of 100. See the supplement for details on generic-level relationships.

**Figure 1.**
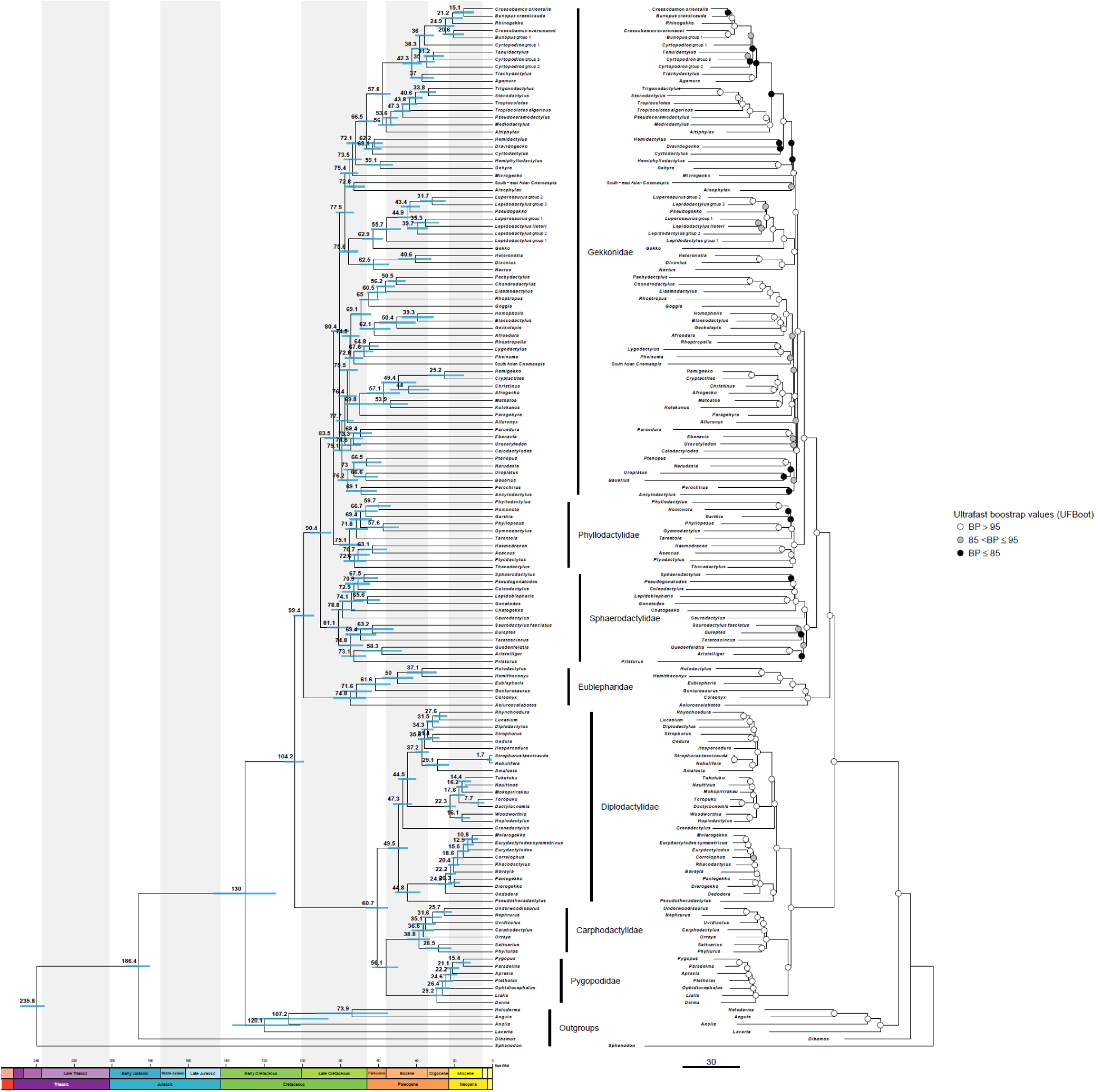
The dated phylogeny of Gekkota including outgroups on the left. The 95% Highest Posterior Density (HPD) estimates of the node dates are plotting as the blue bar with centers on the mean estimates. The topology is restricted to the ML topology returned from the 20-gene dataset, as plotted on the right. The node color is based on the ultra-fast bootstrap values.

### Supermatrix Divergence Dates

The BEAST2 analyses place the crown age of Gekkota to be within the Early Cretaceous with the highest posterior density (HPD) spanning from the age of Cenomanian to Albian (Table 2). All three families of Pygopodoidea have crown ages estimated to fall within Paleogene period. The youngest of these three families is the Pygopodidae which originated in Oligocene with the HPD spanning across much of the epoch. The family Carphodactylidae and Diplodactylidae are slightly older where they originated during the Eocene, again with the HPD spanning nearly the entirety of the epoch. These are in contrast with the Gekkonoidea families where all of them likely have started diversifying during the Cretaceous. These Gekkonoidea families have comparable crown ages where their mean ages are estimated to fall within the age of Campanian.

**Table 2.**
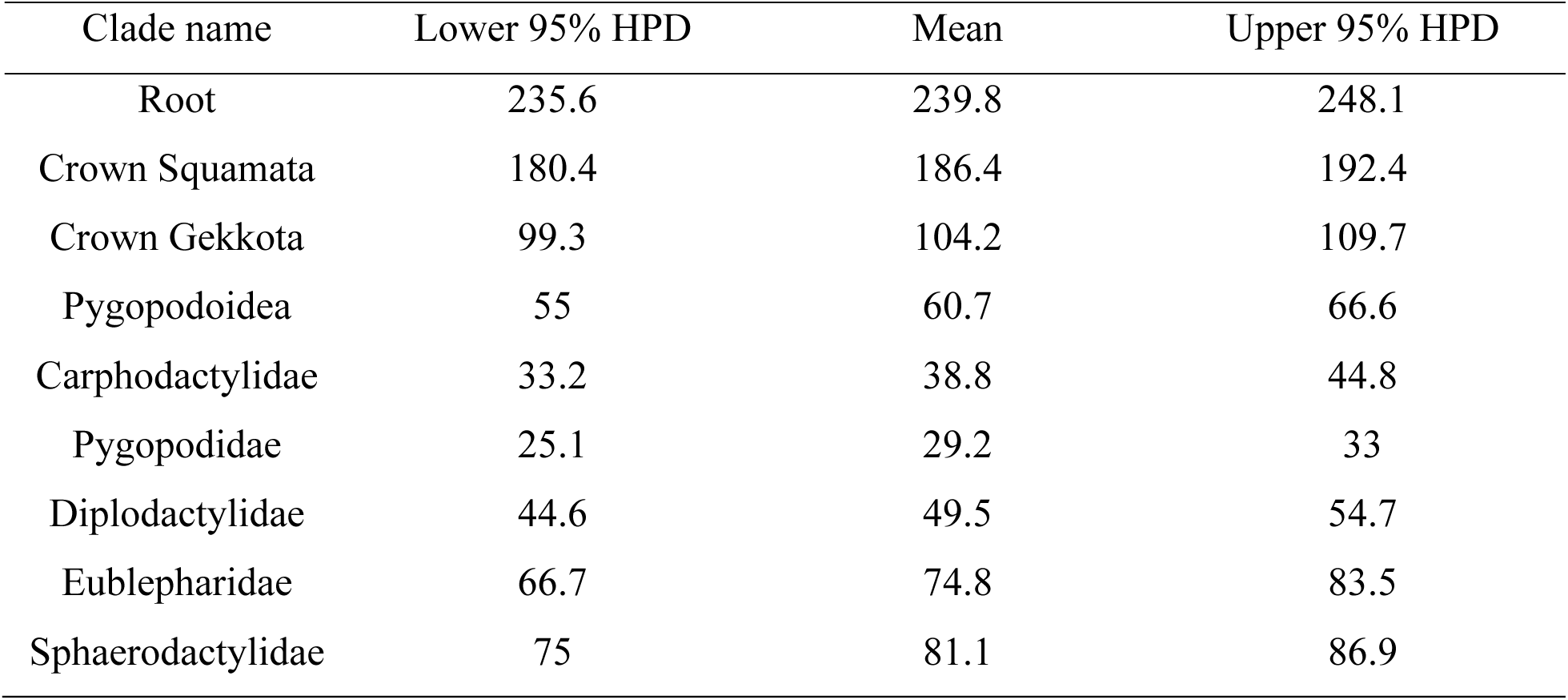
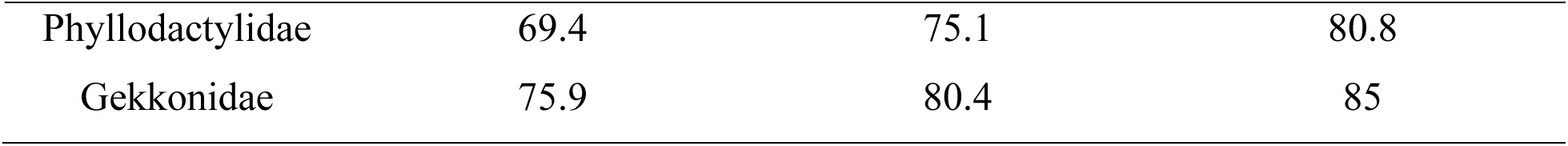
Node age estimated returned from BEAST2 analyses with full dataset with three fossil priors and 15 secondary priors. Topology is restricted to being the ML topology of the 20-gene supermatrix. See supplementary table S4 for comparison of age estimates across different datasets. The unit is in million years ago (my).

### Diversification Dynamics

We ran all diversification analyses on the grafted UCE-supermatrix MCC tree. All birth-death analyses found compelling evidence for diversification rate heterogeneity across geckos. Estimation of the MAP shift configuration of BAMM analysis identifies at least rate 6 shifts in speciation across Gekkota, limited entirely to the Gekkonoidea (*sans* Eublepharidae). All of these shifts were rate increases above the mean background speciation (λ) rate of 0.06 sp/my (mean μ < 0.004). We recovered a doubling in speciation rate (λ = 0.14 sp/my) at the base of the clade containing Sphaerodactylidae, Phyllodactylidae, and Gekkonidae (Figure 4). More were identified near the base of *Sphaerodactylus* (λ = 0.15 sp/my), the crown of east Asian gekkonids (λ = 0.14 sp/my), and near the base of *Cyrtodactylus* (λ = 0.24 sp/my). Two additional shifts were also identified; one deeply nested within the South Asian *Cnemaspis* clade (λ = 0.15 sp/my) and the other at the base of the Sundaland *Cnemaspis* clade (λ = 0.15 sp/my) (Table 3). It should be noted that the abovementioned clade-specific shifts are instantaneous node estimates and that the subtending subtrees uniformly display overall decreases in speciation rate towards the present. The location of shifts identified by BAMM were consistent when perturbating prior choice on the expected number of shifts, with *expectedNumberOfShifts* = 3 resulting in the same 6 shifts. BAMM failed to find heterogeneity in pygopodoids or eublepharids. Macroevolutionary cohort analysis suggested that each of the clades that experienced shifts in speciation rate occupy their own distinct rate regime.

**Table 3:**
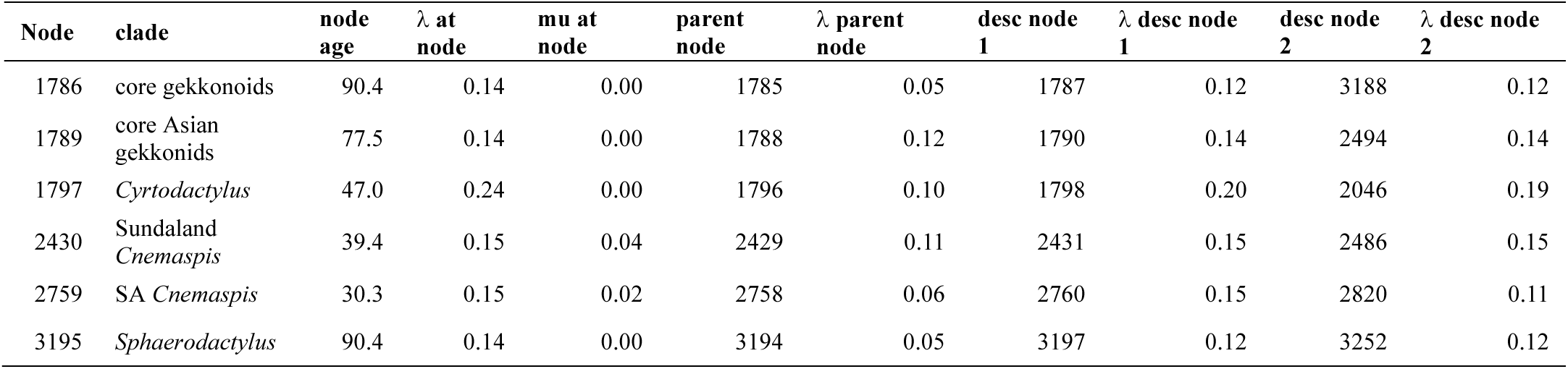
Speciation rates estimate by BAMM for nodes recovered to have experienced rate shifts. Included are clade age (my), regime, λ on the node subtending the node of interest, instantaneous λ at node of interest, μ at the node of interest, and λ on the two descendant nodes of the node in interest.

Where BAMM identified strong support for several large-magnitude shifts, our analysis with CLaDS in combination with RRphylo located a large number of small shifts throughout the grafted tree (N = 47) (Table S12). As with our BAMM assessment, heterogeneity was overwhelmingly concentrated in gekkonoids (N = 43), though 4 shifts were recovered in pygopodoids. The only family that did not experience shifts based on the CLaDS2 model was the Eublepharidae. Surprisingly, most rate shifts were decreases from the background rate (N = 29). Only the Sphaerodactylidae seems to be biased to negative shifts relative to the background rate while all other clades appear to have both rate increases and decreases. All clades estimated to experience shifts in BAMM were found to have small, but significant shifts in CLaDS. Though, the CLaDS model did find a number shifts not indicated by the BAMM model under any shift prior configuration. Our investigation of potential mass extinctions revealed no support for such an event in Gekkota with all mass extinction BFs being well below 3.

### Biogeographic Reconstruction

Without fossils included, the DEC*+j+w+x* model (model included jump dispersal, distance and manual dispersal multiplier matrices) was the best supported model across 10 different trees from the posterior, with AICc weight ranging from 0.62 to 1 (Table 4; Table S8). Adding fossils randomly to the phylogenies did not impact model selection, with the DEC*+j+w+x* model having AICc weights ranging from 0.57 to 0.91. This suggests that both distance and position among regions play a large role in model fit for these data, not the inclusion of fossil taxa, supporting both “Distance” and “Position” hypotheses.

**Table 4.**
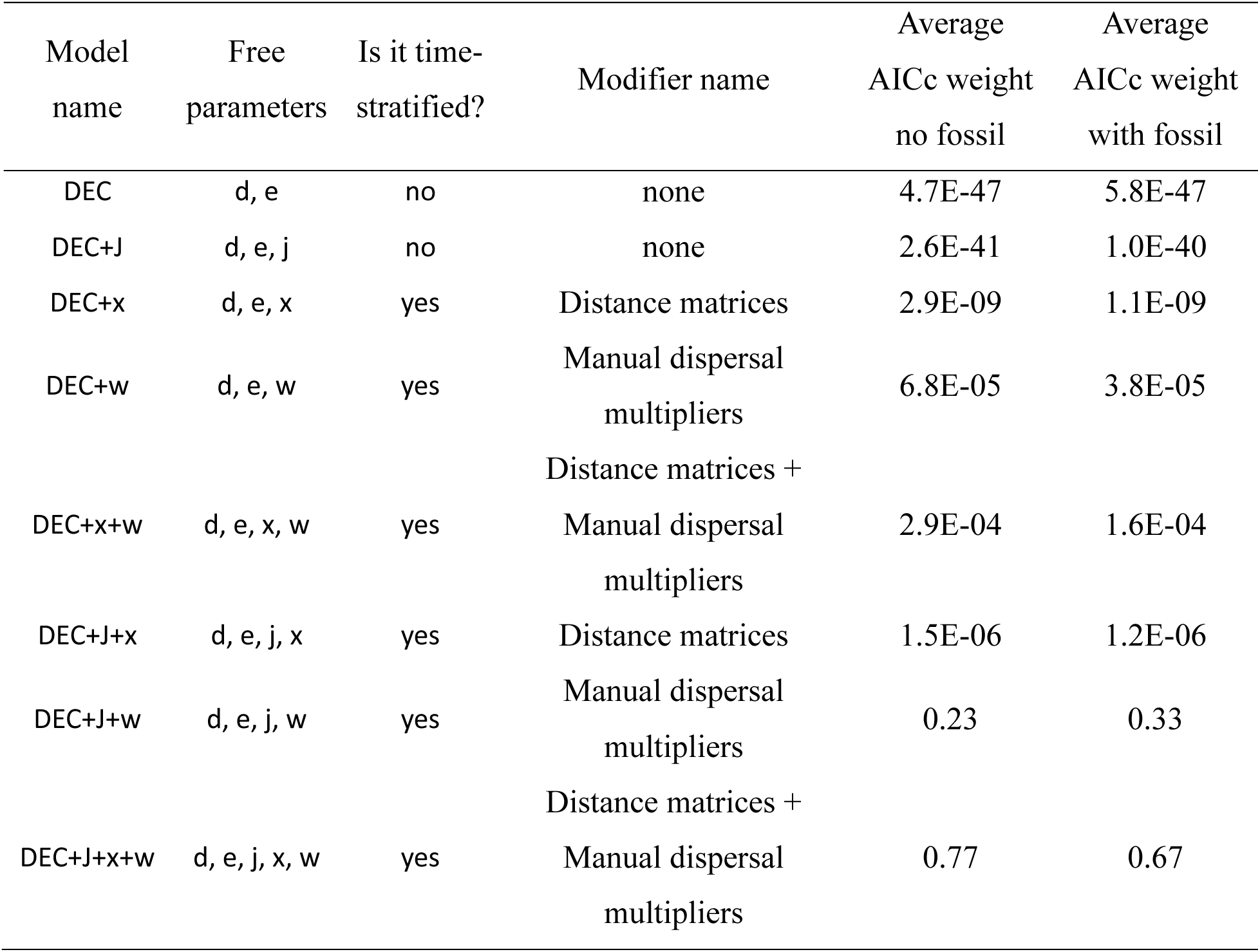
Models fitted in BioGeoBEARS. d: dispersal rate (anagenetic); e: extirpation rate (anagenetic); j: jump dispersal weight (cladogenetic); x: distance matrix modifier power where d_actual_ = *d_base_* ∗ *distance^x^* and *j_actual_* = *j_base_* ∗ *distance^x^*; w: Manual dispersal multiplier power where d_actual_ = *d_base_* ∗ (*manual dispersal multiplier*)^w^ and *j_actual_* = *j_base_* ∗ (*manual dispersal multiplier*)^w^. The averaged AICc weight is averaged across 10 BioGeoBEARS runs with different grafted UCE-Supermatrix time trees (same topology, different node ages). Fossils were randomly attached to each time tree prior to each run, resulting in different fossil topology for each run. For more detailed results, see table S8.

Adding fossils randomly to phylogenies, especially to the root, did affect ancestral state reconstruction for crown Gekkota. Without fossils, the ancestral state was inferred to be in Africa-Arabia-Eurasia-Australasia (ACF) across all 10 phylogenetic trees (Figure S10). The Biogeographic Stochastic Mapping (BSM) corroborated this result. Across 1000 BSM maps (10 trees with 100 BSM maps per tree), 483 maps support Africa-Arabia-Eurasia-Australasia as the ancestral state, followed by 101 maps support Africa-Arabia-Australasia-North-Central America (AFH) as ancestral state for crown Gekkota (Figure 2A). Adding fossils changed the ancestral state for crown Gekkota, with the ancestral state inferred to be Africa-Arabia-Eurasia-Australasia (ACF) in only 4 out of 10 phylogenies, Eurasia-Australasia (CF) in 5 of them and Eurasia (C) in the remainder (Figure 3; Figure S9). BSM maps returned similar results as well. A total of 313 maps supported Africa-Arabia-Eurasia-Australasia as the ancestral state, 336 maps supported Eurasia-Australasia while 182 supported Eurasia as the ancestral state (Figure 2B).

**Figure 2.**
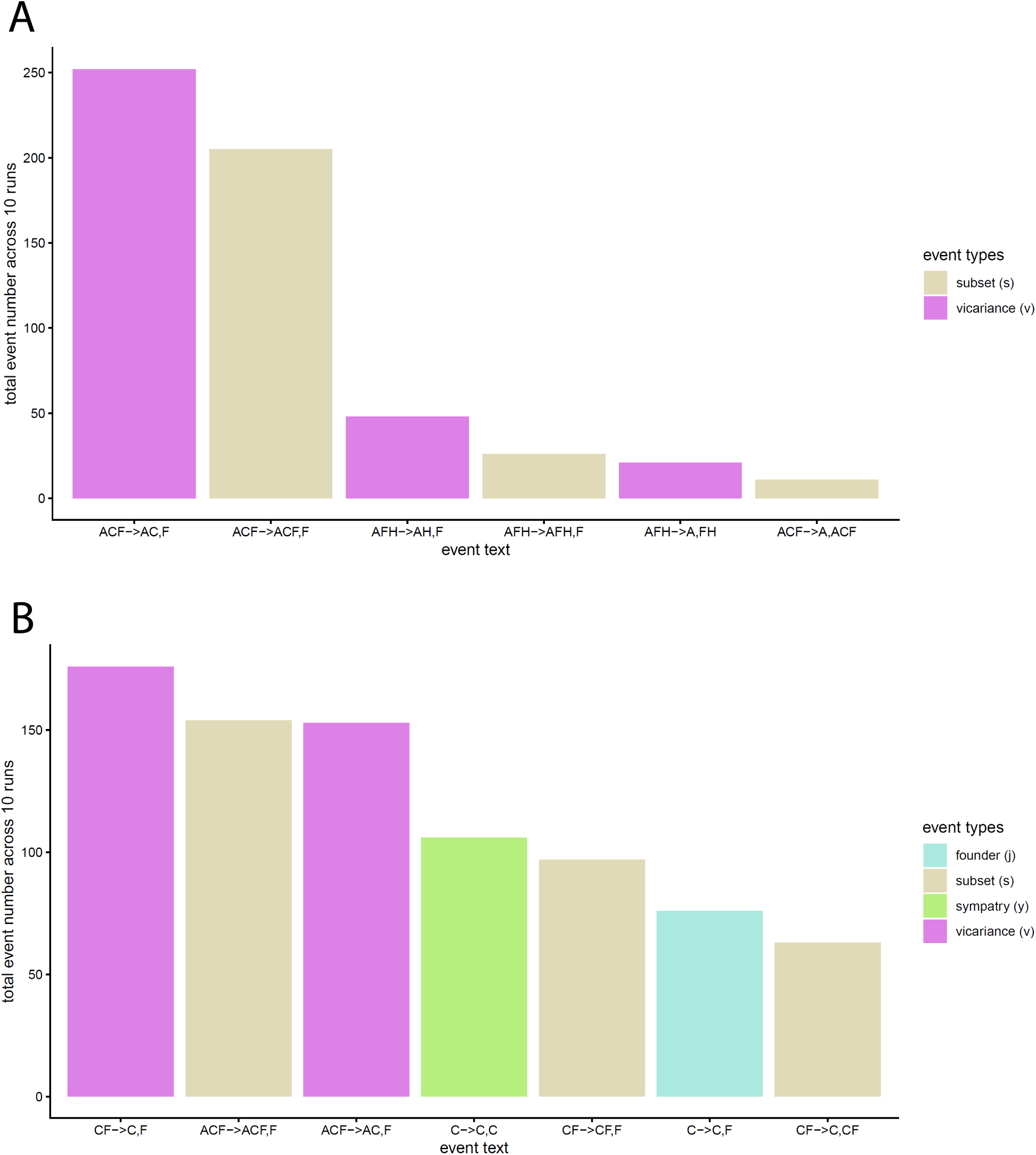
Cladogenetic events frequencies for the crown gekkota node. A: Crown gekkota states and events with no fossil added to the root. B: Crown gekkota states and events with fossil randomly added to the phylogenies, including the root. The event text describes the biogeographic event that happened. For example, CF->C,F are describing an ancestral state of CF with subsequent vicariance events on this node with one daughter lineage inherited state C while another daughter lineage inherited state F. State coding: A-Africa-Arabia; B-Madagascar; C-Eurasia; D-South Asia; E-Sundaland; F-Australasia; G-Carribean; H-North-Central America; I-South America.

**Figure 3.**
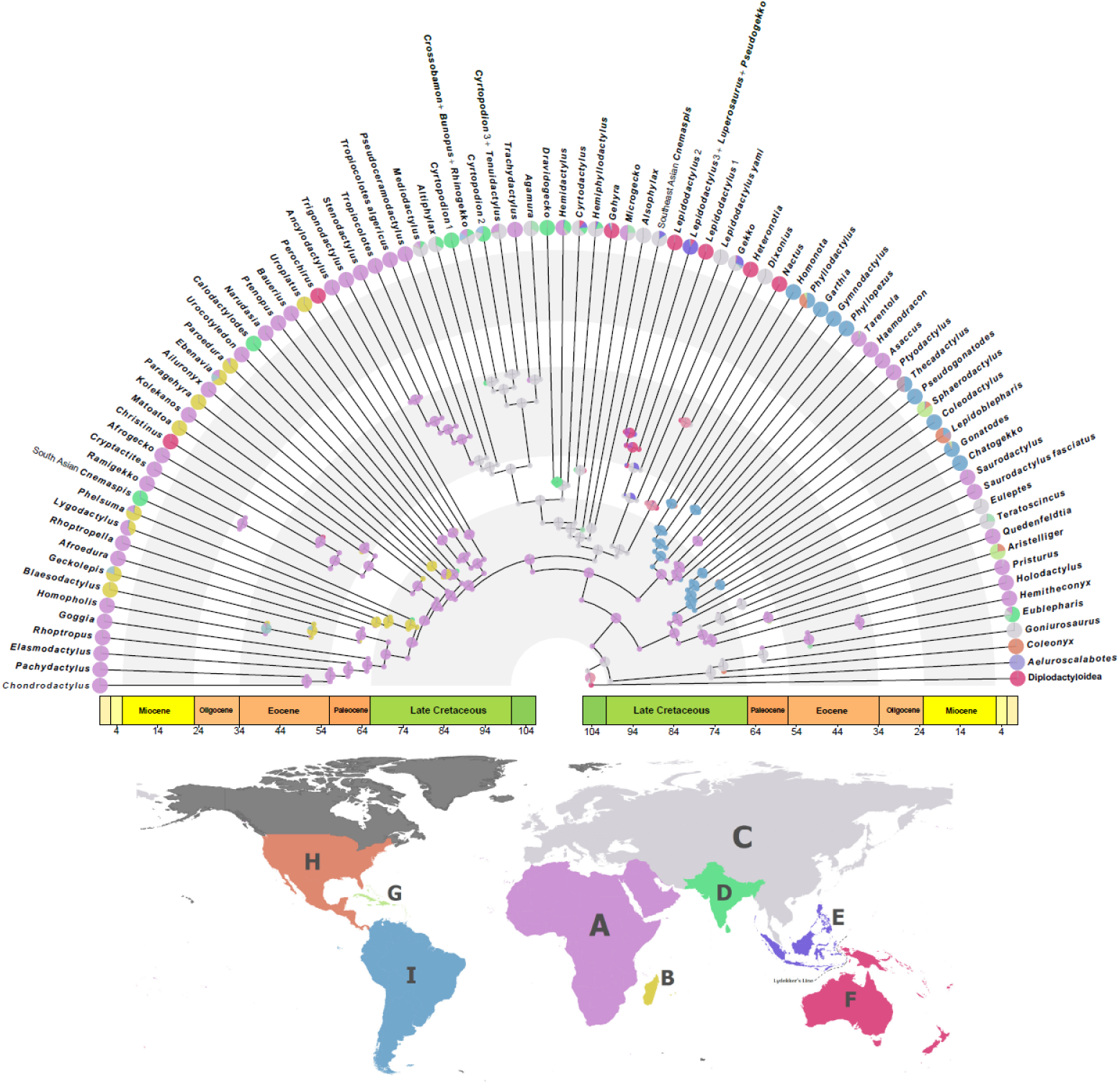
The ancestral reconstruction summarized from the ten BioGeoBEARS runs with fossil randomly attached to the grafted phylogeny, and with DEC*+j+w+x* model fitted. The fossil taxa are removed before plotting. The pie on the node represents proportion of each state realized across the ten runs. The pie on the tips represents the proportion of each state within each clade. The ancestral state pie has inferred states as following: 5/10 as CF, 4/10 as ACF, and 1/10 as C. State coding: A-Africa-Arabia; B-Madagascar; C-Eurasia; D-South Asia; E-Sundaland; F-Australasia; G-Carribean; H-North-Central America; I-South America.

The ancestral state for the Pygopodoidea is estimated to be in Australasia (F) and was supported by 970 BSM maps. The ancestral state of the family Eublepharidae is estimated to be in Eurasia (C) while the ancestral states of the family Sphaerodactylidae, Phyllodactylidae and Gekkonidae were all inferred to be in Africa-Arabia (A). These results are also supported by 747, 893, 648, and 595 BSM maps respectively without fossil, and by 887, 819, 611, and 480 BSM maps respectively with fossil.

### Biogeography and Diversification

Both hidden states and the observed biogeographic area appear to affect net diversification rate. Species with hidden state 0 diversify significantly faster than species in hidden state 1 (Figure 5A; figure S14). Within hidden state 0, species inhabiting Eurasia, South Asia, and Australasia have the highest net diversification rates while species inhabiting Africa-Arabia, Madagascar, and South America have the lowest diversification rates (Figure 5B; Figure S15A). Species in Sundaland, the Caribbean, and North-Central America have intermediate diversification rates, and they are not significantly different than rates observed in other states. A similar pattern was found with hidden state 1 as well. Species inhabiting Eurasia, South Asia, and Australasia still have higher diversification rates compared to species in Africa-Arabia, Madagascar and South America, but only some comparisons are significant (Figure 5C, Figure S15B). Species in South Aisa have significantly higher diversification rates compared to species in Africa-Arabia, Madagascar and South America, while the diversification rates of species in Eurasia are only significantly higher than those of species in South America. Similarly, diversification rates of species in Australasia are only significantly higher than that of species in Africa-Arabia. Consistent with patterns seen within hidden state 0, species in Sundaland, Caribbean, and North-Central America have intermediate diversification rates, and they are not significantly different than other rates.

The ancestral reconstruction of the MuHisse model further corroborated the results in our BioGeoBEARs analyses. The ancestral states of crown Gekkota was inferred to be Africa-Arabia or Eurasia with the posterior probabilities of 0.54 and 0.42, respectively (Figure S25). The model also inferred that the ancestral states would be Australasia with a low probability of 0.02. The ancestral states for the family Eublepharidae were estimated to be in Eurasia with posterior probability of 0.66 and Africa-Arabia with posterior probability of 0.34. The ancestral states of the family Sphaerodatylidae, Phyllodactylidae and Gekkonidae are all inferred to be in Africa-Arabia with posterior probability of 0.98, 0.99 and 0.95 respectively.

## Discussion

### Supermatrix Topolgy and Divergence Dates

At the familial level, our topology is consistent with other concatenated approaches using either Sanger data (Gamble et al. 2012, Gamble et al. 2015, Zheng and Wiens 2016), or phylogenomic data (Skipwith et al. 2019, Title et al. 2024: topology produced by their fully unconstrained approach). However, the placement of some genera may be controversial. For instance, the placement of *Perochirus* has been historically problematic, though some hypotheses place it near the *Gehyra* + *Hemiphyllodactylus* clade with relatively low statistical support (Heinicke et al. 2011, Gamble et al. 2012, Gamble et al. 2015). Morphologically, *Perochirus* shares the same paraphalageal pattern with *Gehyra* and *Hemiphyllodactylus* (Russell and Bauer 1988). Our phylogeny, however, supports a sister relationship of *Perochirus* with the African *Ancylodactylus*, suggesting an alternative scenario of morphological and biogeographic evolution. Morphological similarities between *Perochirus* and *Gehyra* and *Hemiphyllodactylus* may be due to convergent evolution rather than shared ancestry. Biogeographically, this suggests a potential trans-oceanic dispersal event to Micronesia region from Africa rather than from Eurasia. This is indeed possible as another genus, *Christinus,* is hypothesized to have dispersed to Australia from Africa, potentially riding oceanic currents or using Antarctica as a stepping stone (Heinicke et al. 2014). Without further fossil and molecular evidence, however, we should interpret this relationship with caution.

The divergence between geckos and other squamates was estimated to have occurred around 130 mya (95% HPD [114.2, 147.1]), which is congruent with the fact that the oldest, widely agreed upon pan-gekkotan fossil *Hoburogekko suchanovi* was found in lower Cretaceous from around 100.5 to 125 mya (Aptian-Albian) More recent studies have placed some older fossils found in Tithonian (145–152 Mya) deposits, such as *Eichstaettisaurus*, *Ardeosaurus*, *Limnoscansor*, and *Helioscopos*, within pan-gekkota (Conrad and Daza 2015, Simões et al. 2017, Conrad 2018, Simões et al. 2018, Meyer et al. 2023). This suggests that pan-gekkotans potentially started diversifying in the late Jurassic or earlier. Even though a more complete total evidence phylogeny with intense sampling of both extant and extinct lineages is required to confidently estimate the age of diversification for pan-gekkota, current evidence, including ours, points to a potential Early Cretaceous or Late Jurassic origin of pan-Gekkota. Our age estimate for the crown Gekkota (mean = 104.2, 95% HPD [99.3, 109.7]) is significantly younger than previous gecko phylogenies (137.4 mya in Zheng & Wiens 2016; 115.2 mya in Title et al. 2024). We used *Cretaceogekko burmae* as our primary calibration point for crown Gekkota. This fossil was initially thought of as a member of the extant gecko family Gekkonidae (Arnold and Poinar 2008), but later placed within Gekkota *sensu lato* due to the fact that it’s digital scansor morphology differs from all extant geckos (Daza et al. 2014, Bauer 2019). This fossil has been used as a calibration point for crown Gekkota in previous studies as well, albeit with differing prior distributions (Nielsen et al. 2011, Skipwith et al. 2019). However, whether this fossil truly represents a crown gecko requires further review since stem geckos already show evidence of scansorial adaptations and the presence of subdigital pads (Simões et al. 2017, Fontanarrosa et al. 2018). Nevertheless, our age estimate of the crown Gekkota age agrees with the consensus that crown geckos had originated by the late Cretaceous (Simões et al. 2025).

### Diversification dynamics and mass extinction

When implementing state-agnostic analyses of diversification, we find strong evidence of rate heterogeneity across crown gekkotans. Notably, our estimate of background speciation rate (mean = 0.06 sp/my) agrees with that recovered for pygopodoids alone (Skipwith and Oliver, 2023). Some of the recovered shifts appear to coincide with biogeographic shifts as inferred by BSM analysis. The failure of BAMM to recover evidence of heterogeneity within pygopodoids concurs with the findings of Skipwith and Oliver (2023), suggesting that something fundamentally differs between this lineage and gekkonoids in lineage proliferation. BAMM consistently recovers a large shift at the mrca of Sphaerodactylidae-Phyllodactylidae-Gekkonidae, a node robustly estimated as being ancestrally Africa-Arabian in origin (Figure 4). This supports the importance of this region being an important source of diversity for the three most speciose families. While we do not explicitly test trait evolution in geckos, it is worth noting that these three families are the only geckos that lay calcareous, desiccation-resistant eggs. Future studies should explicitly test whether this novel trait is indeed a key innovation driving speciation. The rapid increase in diversification rate recovered within *Sphaerodactylus* appears to correspond with the colonization and subsequent proliferation of the West Indies from Central America. Similarly, the shift identified at the base of Asian gekkonids corresponds with the colonization of Eurasia from African ancestors. Interestingly, the shifts identified in South Asian *Cnemaspis*, Sundaland *Cnemaspis*, and *Cyrtodactylus* do not seem to be linked to major biogeographic transitions. Where ecological opportunity due to the colonization of a novel region is linked to increased speciation in *Sphaerodactylus*, the success of these other lineages may instead be linked to intrinsic traits or local environmental perturbations that facilitated proliferation within their respective centers of origin. *Cyrtodactylus* exhibited the highest speciation amongst gekkotans, which is unsurprising for the largest and most rapidly growing gecko genus in terms of newly described species (∼400 species). It is also possible that continued movement of Sundaland and the formation Wallacea drove speciation rates in *Cyrtodactylus* and Asian gekkonids *writ large*, a pattern reminiscent of body-size evolution suggested for the former genus (Oliver et al. 2014). The dozens of low-magnitude speciation rate shifts recovered by CLaDS largely, but not entirely, subtend nodes that experienced heterogeneity in BAMM. The pattern that these seem to largely indicate drops in rate fall in line with commonly proposed patterns of diversity dependence where speciation rate declines after an initial burst. However, such inferences are highly problematic and we recognize that any signal of diversity dependence from such a large phylogeny may be spurious on ecological grounds, notwithstanding the issues of model misidentification that are ubiquitous in studies relying on birth-death models on extant-only trees (Louca and Pennell 2020, Helmstetter et al. 2022). More generally, rate shifts identified by CLaDS are all clustered in relatively recently diverged lineages as none are older than 47 my in our grafted MCC tree, suggesting that different factors may have played a role in younger clades with limited distributions.

**Figure 4.**
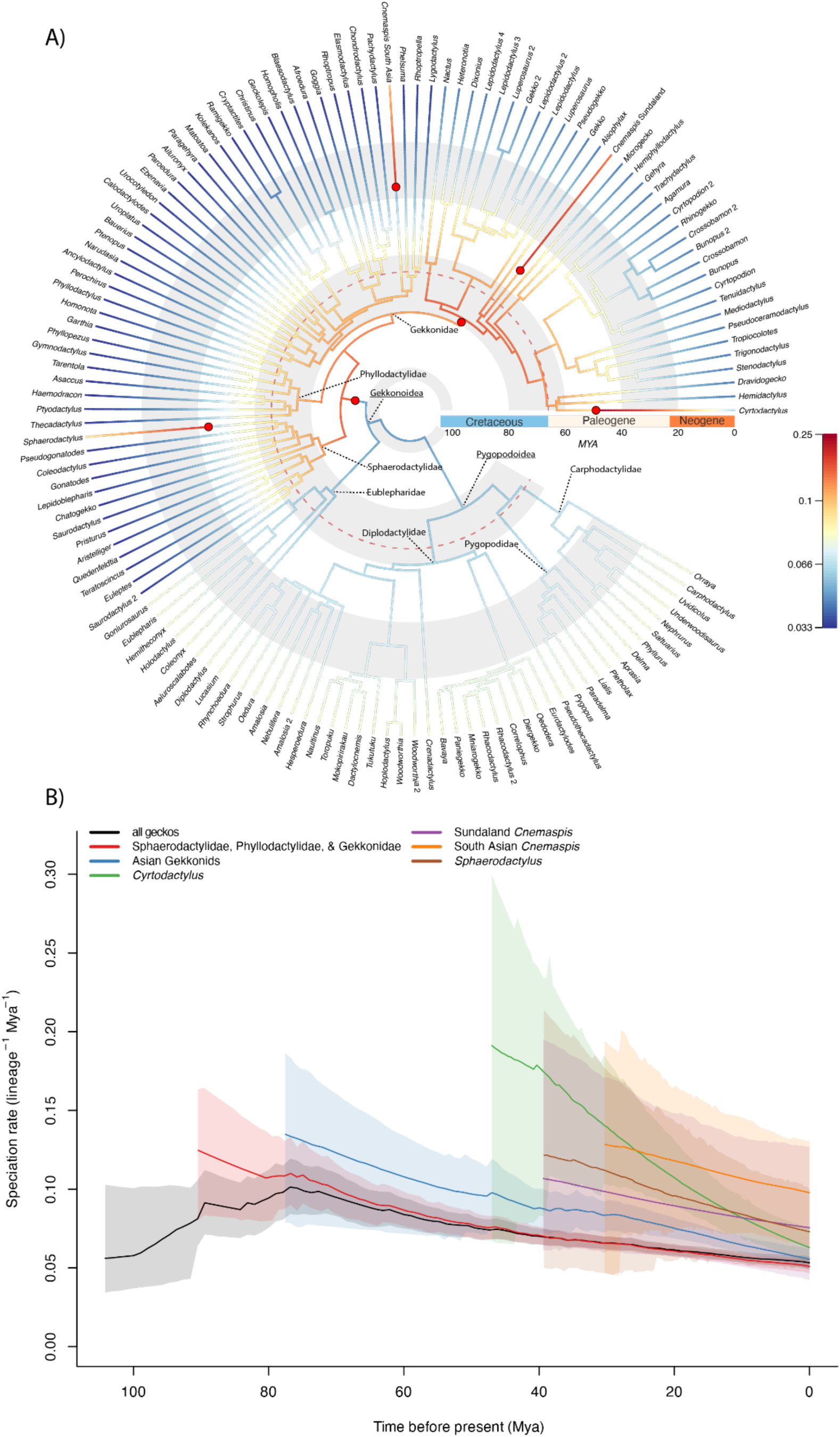
Speciation rates across geckos estimated by BAMM. (A) Time-calibrated, genus-level grafted tree (UCE-supermatrix MCC) displaying speciation rates across Gekkota from the best shift configuration (shift prior = 2). This topology simplified from the entire UCE-supermatrix tree of 1,783 species (see Figure S23). Branch color corresponds to rate and red dots approximate where shifts in speciation occurred (see Figure S23 for details). Red-dotted libne corresponds to the K-Pg mass extinction event 66 mya. (B) Rate-through-time plots of speciation from BAMM for all geckos (black) and for the clades that experienced rate shifts. The dark line inside each curve represents the mean and opaque areas represent the 95% HPDs of each clade.

Our finding that crown geckos did not experience mass extinction agrees with the findings of Skipwith and Oliver (2019) on pygopodoids alone, suggesting that neither gekkonoids or pygopodoids underwent extensive turnover. This is rather surprising as crown geckos, and all gekkonoid families had begun radiating prior to the K-Pg extinction event and all seven families were present by the early Paleogene. While extinction has likely played a major role in the diversification of geckos, it is possible the clade survived catastrophic events with no long-lasting impacts that are detectable with the rjMCMC model used here. Further analyses investigating extinction dynamics and fossil occurrences for individual gecko clades are warranted as clade-specific episodic extinction may be masked by the overwhelming signal of the whole tree.

### Biogeographic Reconstruction and Diversification

The DEC+j+w+x biogeographic model is consistently the best model regardless of the inclusion of fossils. This lends support to both the “Distance” and “Position” hypotheses, meaning that the biogeographic pattern can be explained better when we consider both the distance among the regions as well as where the regions are relative to one another. We also found that the position hypothesis (e.g. DEC+(j)+w) on average explained the geographic pattern better than the distance hypothesis (e.g. DEC+(j)+x), and often was the next best model (Table S8). Geckos are generally good overwater dispersers, with many proposed long-distance transoceanic dispersal events across the phylogeny (Gamble et al. 2011, Heinicke et al. 2011, this study, Skipwith et al. 2016). This is possibly due to their unique suite of traits such as adhesive toepads, desiccation-resistant eggs, and egg gluing behavior (Andrews 2012, Gamble et al. 2012, Choi et al. 2018, Meiri 2019), making them well suited for surviving long distance over water. The relative positions of landmasses may be more important because oceanic currents and wind patterns are important factors for passive dispersers such as geckos. This is evidenced by there being at least two dispersal events from Africa to the Caribbean region but none the other way around.

When excluding stem taxa, the ancestral state of crown geckos is estimated to be Afro-Arabian, Eurasian and Australasian, contrasting with the east Gondwanan origin hypothesis (Oliver and Sanders 2009). Importantly, the ancestral state of the crown geckos is reconstructed to have increased likelihood to be confined in only Eurasia-Australasia, or in just Eurasia when fossils are randomly added to the root (Figure 2B). This is perhaps not surprising given all of the stem-gekkotan fossils included in this study were found in Eurasia. This emphasizes the importance of considering fossils when reconstructing biogeographic areas since extinctions can bias ancestral area reconstructions (Wood et al. 2012a, Yan et al. 2021). Even though the relationships of fossil pan-gekkotans are uncertain, we can explore the effects of incorporation of the taxa on ancestral area reconstructions by randomly attaching these fossils at the root. However, without further evidence, we are unable to draw any definitive conclusions. Further, there little geological evidence that such a distribution was possible as this reconstruction suggest areas that were not contiguous at the time. The biogeographic analysis of Meyer et al. 2023 including additional stem gekkotans (including *Gobekko* and *Hoburogekko*) suggested that the ancestral area of the crown geckos is Asia (their Figure 4). However, their phylogeny supported many relationships not recovered in any molecular study, either with Sanger or genomic scale datasets (Gamble et al. 2015, Zheng and Wiens 2016, Burbrink et al. 2020, Singhal et al. 2021). Another study dealing with the biogeography of squamates as a whole using fossil taxa as geographic occurrence constraints, meaning the ancestral area of the node with fossil constraints must include the area of the fossil (Wilenzik et al. 2024). They estimated the ancestral range of Gekkota to be Pangea. Our BSM maps suggested somewhat contradictory results, where the ancestor of the crown gekkotans likely originated in the Old World, possibly in Eurasia, well after the breakup of Pangea. We did not include Antarctica in our biogeographic analyses, but it is possible that it might have acted as a corridor between other Gondwanan landmasses. The region was considerably warmer during parts of the Cretaceous, with evidence of temperate rainforests existing in Antarctica during the Late Cretaceous (Klages et al. 2020). Given the statistical ambiguity at the crown of Gekkota in our BSM analyses, we acknowledge that we cannot suggest any region as the likely center of origin for gekkotans beyond the Old World.

The ancestral ranges of the families Sphaerodactylidae, Phyllodactylidae, and Gekkonidae were all inferred to be Africa regardless if fossils were incorporated, a finding supported across BSM maps as well. Sphaerodactylids and phyllodactylids then dispersed to the New World from Africa independently five times (two times in sphaerodactylids and three times in phyllodactylids), agreeing with the estimates of Gamble et al. (2011). Our reconstruction generally agrees with their second hypothesis when it comes to the origin of the New World Phyllodactylidae, where there is back dispersal from South America to Africa in the ancestor of the *Tarentola*, although this is not supported in our BSM maps or MuHisse analysis (Figure S9-10, S25). Within Gekkonidae, extant species have a predominantly Old-World distribution, but ancestral centers of diversity can be loosely summarized in terms of two clades. One clade diversified *in situ* within Africa and Madagascar (e.g. *Pachydactylus* clade, *Lygodactylus*, *Phelsuma* clade, etc.) with one major dispersal event from Madagascar to the Indian subcontinent leading to the diversification of the Indian *Cnemaspis* clade. The sister of this predominately African clade is a speciose clade that colonized Eurasia around 80 mya from Africa, with many subsequent invasions to Sundaland, Australasia, Indian subcontinent and the New World. Further, this predominately Asian clade gave rise to lineages that would go on to recolonize Africa (e.g. *Hemidactylus*, Palearctic bent-toed geckos). Despite the fact that early stem gecko fossils are known from Eurasia, the ancestors of the majority of the extant gekkonoid families did not originate in this region. Rather, they are very likely representing an out-of-Africa diversification pattern. The only exception are the predominately Eurasian eublepharids (e.g. *Goniurosaurus*) which originated and diversified in Eurasia. Not only was Africa a potential place of origin for three out of four extant gekkonoid families, but our chord diagram (Figures S11-12) also indicates that it was an important source for dispersals. There are dispersal events out of Africa (emigration) to six other regions, more than any other regions (Figure S11). However, the total number of events overall is low compared to Eurasia which has about twice as many emigration events per BSM estimation (Table S11). Similarly, the dispersal events into Africa (immigration) are also diverse (from four other regions), but the total number of events is about 1.5 times less than Eurasia. Due to low emigration and immigration rates in conjunction with the fact that Africa has one of the most diverse gecko faunas (∼21.5% of all species) and one of the lowest diversification and turnover rates (Figure 5, S21), we argue that Africa has acted as a biodiversity “refugia” for geckos. While supporting a comparably diverse gecko fauna (∼20.2% of all species), Eurasia has both a higher number of dispersal events compared to Africa and has some of the highest diversification and turnover rates. This was also suggested by our state-agnostic diversification analyses where we find strong support for Eurasia acting as a major engine of diversity as evidenced by elevated rates being detected throughout the core Asian gekkonid clade as inferred by BAMM and ClaDS. These patterns suggest that Eurasia may act as both a major dispersal corridor and a diversification hotspot for geckos. It is very likely that the close geographic connection between Africa and Eurasia combined with very different diversification patterns have influenced global gecko diversity through cycles of persistence, diversification, and dispersal. Since “hotspots” have both high speciation and extinction rates, reconstructing biogeographic history may be potentially be biased towards “refugia” (Vasconcelos et al. 2022). This is possibly evident here where incorporating stem fossils into biogeographic reconstruction led to the estimation of the ancestral range of crown Gekkota to be more biased towards “hotspots” (Eurasia) but not “refugia” (Africa), albeit with high uncertainty.

**Figure 5.**
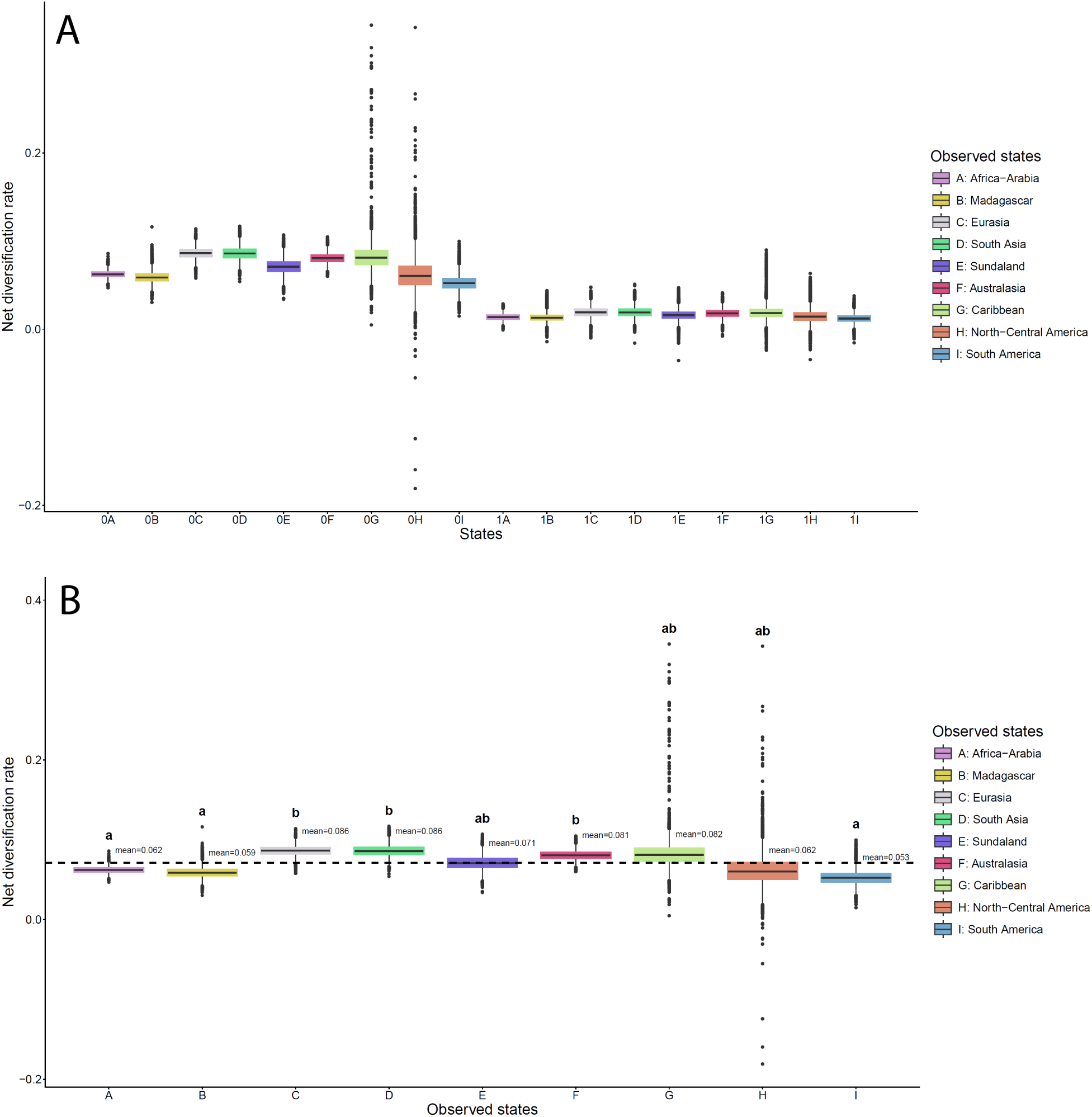
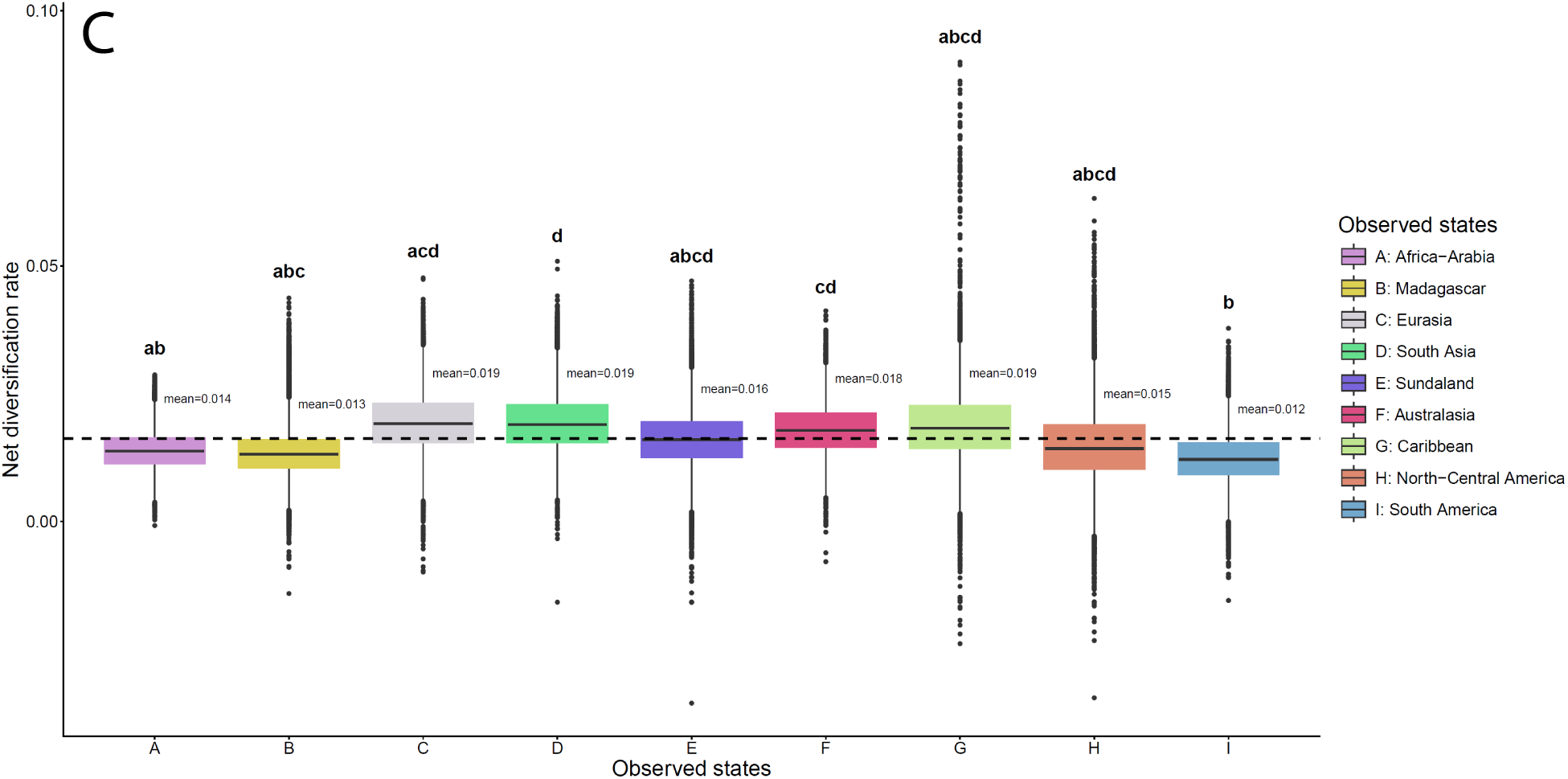
Boxplot showing posterior estimates of the net diversification rate (speciation rate – extinction rate) from MuHisse model with two hidden states. A: boxplot of all observed state and hidden state combinations together showing that species in hidden state 0 have higher net diversification rate than species in hidden state 1. This difference is significant (see also Figure S11). B & C: boxplot of net diversification rate estimates within hidden state 0 and 1. Letter above the box indicating group, with different letters indicating significant differences (see also Figure S12). The thick dash line indicates the mean rate of this hidden state. Observed state coding: A-Africa-Arabia; B-Madagascar; C-Eurasia; D-South Asia; E-Sundaland; F-Australasia; G-Carribean; H-North-Central America; I-South America.

## Conclusions

By leveraging a supermatrix dataset, we use the largest estimate of gecko relationships to date for inference in diversification dynamics and biogeographic history. Our findings contribute to growing evidence that the cosmopolitan gekkonoids truly do stand apart from pygopodoids, which have received more attention in recent macroevolutionary studies, in diversification dynamics. Further, we demonstrate biogeographic history has had major, long-term impacts on gecko diversification. Under this paradigm, we find little evidence for a strictly Gondwanan origin but more generally one that is very likely Old-World in nature while Africa and Eurasia may have acted as refugia and diversification hotspots/dispersal corridors, respectively.

## Acknowledgement

We thank Dr. Paul Oliver for providing many interesting ideas for the divergence dating analyses. We thank Dr. Jenna M. McCullough for the big help with the use of BEAST2 software. We thank Dr. Rosana Zenil-Ferguson for generously providing the example script of the MuHisse analysis and for the tremendous help with the RevBayes software. We thank Dr. Aaron Bauer for many insightful and helpful comments.

## Author Contributions

Both authors contributed equally to the conceptualization and the design of the study. YL has led the data analyses while PL contributed to the diversification dynamic analyses. YL has led the writing of the manuscript while PL wrote the introduction section. Both authors contributed equally to the reviewing and revising the drafts and both authors approved the final version.

## Data Availability

Raw genomic from Skipwith et al. (2019) are available on the Mendeley Data Platform (doi: 10.17632/zstmhjfkw5.1). Additional information on analyses are available in the supplement of this manuscript.

