## Supplementary Material for "Global biogeography of geckos (Squamata: Gekkota) reveals Old-World origins and region-specific heterogeneity in diversification"

Taxon sampling and phylogenetic analyses procedures

To make sure that the sequences we acquired were correctly identified, several precautions were taken: 1) The original publication of the sequences was checked to see if the sequence accession numbers were actually present in the study and in the phylogeny, and if the sequences were correctly labeled. 2) Each accession number on GenBank was attached to a species name and this name reflected the taxonomic information at the time. However, many species names have gone through changes since the first publication with more information available (e.g. changes of the genus names and/or the recognition of cryptic species within a species complex). Often times, the names on GenBank do not reflect this change. To make sure the sequences in our study were correctly identified with the most up-to-date species names, we cross-referenced the references on GenBank with information on The Reptile Database (Uetz et al. 2025) and peer-reviewed publications using Google Scholar. The names were updated if they appeared in a latter publication and were revised.

Previous gecko phylogenetic studies, ranging from species descriptions to large squamate supermatrix phylogenies, have relied heavily on Sanger data. Therefore, we focused solely on and mined Sanger sequence data from GenBank. Species with phylogenomic data (e.g. UCEs, AHEs) but without Sanger data were excluded in our initial phylogeny building (e.g. Wagner et al. 2014, Ramírez-Reyes et al. 2020). To align protein coding loci, we first adjusted the reading frame using Coding_Translation_Tests.py module in superCRUNCH 1.3.2 (Portik and Wiens 2020). We then used the Align.py module in superCRUNCH with *-a macse* flag to run MACSE 2.07 alignSequences program (Ranwez et al. 2018), and with both *--pass_fail* and *--accurate* flags turned on. Coding_Translation_Tests.py script occasionally returns two sets of sequences, one set that passed the translation test (high quality set) and was correctly adjusted, and one set that failed the translation test (lower quality set). The --*pass_fail* flag was used to automatically adjust aligning algorithm of alignSequences program based on sets quality. That is, the frameshift and stop codon costs for the lower quality set are different than those used for high quality set. Turn on *--accurate* flag sets the both the *-local_realign_init* and *-local_realign_dec* flag to 0.9 in alignSequences program, meaning the program will perform a more through refinement strategy to improve alignment in their refinement loop steps.

Effects of missing data

Our initial supermatrix had a high percentage of missing data of 95.5% (Table S1). To test if the topology of our phylogenetic inferences were significantly affected by missing data, we first trimmed our 71-gene dataset to a 20-gene dataset by first pruning genes with the highest percentage of missing taxa. This resulted in a 20-gene alignment with 86.9% missing data. These datasets contained the same species as the 71-gene datasets. The same tree search parameters (as in the main text) were applied, and the analysis was repeated three times. To further test the impact of missing data, we further trimmed genes in the matrix to reduce missing data, again starting from genes with highest percentage of missing taxa. This resulted in a 5-gene dataset (1693 species, 69.9% missing data), a 4-gene dataset (1597 species, 61.7% missing data), and a 3-gene dataset (1477 species, 51.3% missing data). The same tree search parameters were applied to each dataset, and each analysis was ran three times in IQTREE 2.2.2.6. We measured information-based generalized Robinson–Foulds distance (Smith 2020) among tree topologies returned using these matrices using function ClusteringInfoDistance() in R 4.4.3 (R Core Team 2025) package TreeDist v2.7.1.9003 (Smith 2020). To control for species missingness, we kept only species present in the 3-gene dataset when comparing across datasets. We normalized the distance against the maximum value, defined in (Smith 2020), which is half of the information content of all the splits in both trees in comparison.

The effects of missing data were minimal (Figure S2, Table S2). When reducing percentage missing data from 95.5% to 86.9% (71 genes to 20 genes), the normalized distance between these topologies is only 2.2%. After reducing the missing data to 69.9% (71 genes to 5 genes), the distance increases drastically to 9.9%. Nonetheless, the normalized distance reached only 14.2% when reducing to 3 genes with 51.3% missing data. In addition, with visual inspection of the topology across datasets, the major monophyletic groupings and non-monophyletic groupings of the genera described in the main text remained largely consistent. It is important to note, however, that this could also be due to the fact that majority of the phylogenetic signals come from a few genes. For example, the ND2 gene has the least missing data and therefore was kept throughout the different datasets. For future analyses, using datasets without ND2 gene (e.g. using nuclear loci) could further explore the effect of missing data across mitochondrial vs. nuclear datasets.

Fossil and secondary calibration distributions and justifications

*Initial three fossil calibration points*

Notes: All calibration distributions used the scale of 100 million years.

1. Crown Gekkota: fossil calibration using lognormal distribution using *Cretaceogekko burmae* as calibration fossil. It was estimated to be 97-100 million years old, from the amber beds that were estimated to be in the Upper Albian (Arnold and Poinar 2008). Therefore, we are setting the lower bound to 97 my (offset = 0.97). We set the mean and the standard deviation so that the mean is about the boundary of Albian (~113 mya) and the 97.5% quantile is within Cretaceous (~145 mya).

$$Lognormal(\mu=-1.94, \sigma=0.62, offset=0.97)$$

2. The split between the Dominican *Sphaerodactylus difficilis* and the Puerto Rican *S. nicholsi* and *S. townsendi*: fossil calibration using gamma distribution using *Sphaerodactylus ciguapa* and *S. dommeli* as calibration fossil (Böhme 1984, Daza and Bauer 2012, Daza et al. 2013a). Thees fossils were estimated to be late Early Miocene to early Middle Miocene of age. Since the age bracket is relatively broad, a gamma distribution was set so that the minimal bound is the end of the Miocene (~5.3 mya, offset = 0.053), and the majority of the density (between 2.5% and 97.5% quantile) is between the start of the Miocene (~23 mya) and Middle Miocene (~12.6 mya).

$$Gamma\left( \alpha=20, \beta=168, offset=0.053 \right)$$

3. Crown age of the New Zealand diplodactylids: fossil calibration using lognormal distribution using a Miocene cranial material from New Zealand (Lee et al. 2009) as calibration fossil. This material was estimated to be Lower Miocene in age. The lower bound is set as the start of the age Burdigalian (~16 mya, offset = 0.16). We set mean and standard deviation so that 97.5% quantile is within Oligocene (~33.9 mya).

$$Lognormal(\mu=-2.5, \sigma=0.37, offset=0.16)$$

*Initial three secondary calibration points*

To be able to better compare with previous studies, some secondary calibrations were also taken from previous publications and using timetree.org websites (adjusted times were used).

4. Age of crown Lepidosauria (root): Uniform distribution ~ ($\min=2.356,\max=2.631$). Previous morphological and molecular clock estimates put the split between Sphenodontia and Squamata somewhere in the Late Permian about 256-260 Mya (Simões et al. 2022, Brownstein et al. 2023, Simões et al. 2025). The range in the Timetree database closely follows these previous studies, we therefore choose this range as our prior on the root.

5. Age of Squamata: Normal distribution $\sim(mean=1.89, \sigma=0.03)$. Previous large-scale phylogenies estimated the age of the Squamata to be between about 173 – 205 mya (Pyron and Burbrink 2014, Zheng and Wiens 2016). Timetree database gave a similar range where it summarized to be between 179 – 204 mya. We choose a distribution close to these ranges where the 2.5% quantile is 183 mya and the 97.5% quantile is 195 mya.

6. Age of Diplodactyloidea: Normal distribution $\sim(mean=0.6, \sigma=0.05)$. Previous phylogenomic study estimated the crown age for Diplodactyloidea is between 65-86 mya with a mean of 76 (Skipwith et al. 2019). The Timetree database summarized a much younger time range of 56-71 mya. Without definite evidence showing which range is closer to reality, we choose a distribution that overlaps both ranges where the 5% quantile is 50 mya and 97.5% quantile is 70 mya.

*Incongruence of time estimation between nuclear and mitochondrial loci*

It has long been recognized that mitochondrial loci generally evolve more rapidly than nuclear loci (Brown et al. 1982, Simon et al. 2006). Because of this faster evolution and potentially earlier saturation, it perhaps not surprising that the divergence dates estimated using mitochondrial loci might be different than those estimated using nuclear loci. But whether those differences are significant enough to produce drastically different evolutionary scenarios, or whether those differences are heterogeneous across nodes, is still worth studying. Indeed, studies have suggested that, overall, mitochondrial loci tend to infer divergence times significantly older compared to nuclear loci, especially towards the tips of the phylogeny (Zheng et al. 2011, Zadra et al. 2021). In addition, some of the estimates do not overlap in their confidence interval, indicating potentially very different evolutionary scenarios, especially for the downstream macroevolutionary analyses such as biogeographic reconstruction and diversification rate estimations. Even though we do not have sufficient sample size to make a robust conclusion about the mutation rate returned from our Bayesian analyses, the estimates of the mean mutation rate (Calculated by $mean substitution rate \times mean clock rate$ for each partition) of our mitochondrion partition are ~ 15.6 to 17.1-fold faster than the mean mutation rate of our nuclear partition, which is comparable to that estimated for various gecko genera in Allio et al. (2017). In that study, they estimated the ratio of mutation rate between mitochondrion and nuclear genes in various gecko genera to be between 17.8 and 34.2.

Given such dramatic difference in substitution rate, we set out to test if there are incongruences of divergence time estimations between our mitochondrial and nuclear loci. We performed the divergence dating analysis with only nuclear loci in BEAST 2.7.7 following the same procedures described in the main text. This dataset (“nuclear-only dataset”, henceforth) contains 1052 species with 85.9% missing data. We pruned the 20-gene ML topology to only include species that have nuclear data. We constrained the topology to be this pruned ML topology by turning off topology operators so only the node dates were estimated in BEAST 2 runs. All six calibration nodes in the initial dataset (“full dataset”, henceforth) were present in this pruned topology. We used the same site model as previously estimated for the nuclear partition. We then ran with five independent chains in parallel each with the same chain length, sample frequency, and brin-in generation, and with the same posterior tree sampling strategy as the previous run with the full dataset.

To compare node estimates between datasets, we preserved the nodes that were present in both the 20-gene ML topology and the pruned topology. We then fitted the linear model using the lm() function in R (R Core Team 2025) using the log-transformed dates estimated in the full dataset as the dependent variable, using the log-transformed dates estimated in the nuclear-only dataset as the independent variable, and using the different datasets as the categorical variable. We initially included the interaction term and tested with type III ANOVA using the Anova() function in the package car 3.1-5 (Fox and Weisberg 2019) to determine if the slopes and the intercepts were significantly different. If there was no interaction, we then tested with type II ANOVA to provide more statistical power. The assumptions were visually checked using the check_model() function in the R package performance 0.18.2 (Lüdecke et al. 2021). Visual diagnostics confirmed that all assumptions were satisfied with negligible deviations.

Overall, the inferred dates using nuclear-only dataset were significantly younger compared to those of the full dataset for mean, lower and upper 95% Highest Posterior Density (HPD) (figure S3; table S3). This was especially prominent for the nodes closer to the tip of the phylogeny. Additionally, lower 95% HPD boundaries of the dates estimated by the nuclear-only dataset tended to be very close to 0 for nodes near the tips (figure S3C). Due to the faster rate of mutation in mitochondrial loci, we might expect them to underestimate the number of mutations and return overall younger dates compared with the nuclear loci. However, we observed the exact opposite pattern, similarly to previous studies (Zheng et al. 2011, Zadra et al. 2021). The underlying reasons for these patterns were unknown, and we believe this could be an interesting topic of future research. Nonetheless, we do believe that the drastically different mutation rate between nuclear and mitochondrial loci had contributed, at least partially if not entirely, to these patterns. Furthermore, since there are no ways for us to determine which date estimates are “more correct”, choosing one set of dates versus another could also introduce biases to the downstream analyses. For the above reasons, we opted to include additional secondary priors in our full dataset analyses for many relatively deep nodes based on our nuclear-only date estimates as well as previous publications, following the assumption that these dates could be closer to reality due the slower mutation rate of the nuclear loci. This way, we hope to combine the advantages of the slow-evolving nuclear loci on estimating those deeper nodes while keeping the mitochondrial loci so the shallower nodes could be reliably estimated.

*Addition eleven fossil calibration points*

Next, we picked 12 additional calibration points based on node age estimates of our nuclear only dataset, as well as ranges summarized in timetree website. In the remaining texts, we will term this dataset as full dataset with additional priors. The 10 out of the 12 calibrations points are relatively deep nodes. We argue that node ages estimated at the deep nodes using nuclear data may be closer to reality compared to those that relied on the dataset with mitochondrial loci, due to the relatively slower evolutionary rates of nuclear loci. The potential problem of shallow calibration points has also been noted before (Duchêne et al. 2014). In addition, the date estimated by the nuclear-only dataset on the shallow nodes could be unreliable due the low rate of evolution of the nuclear loci. These node age estimates are also supported in other publications.

13-17. Age of divergence (diversification) of *Uroplatus*: Normal distribution $\sim(mean=0.35, \sigma=0.1)$. Dates estimated from our nuclear datasets: 16-33 mya.

Age of divergence of *Phelsuma*: Normal distribution $\sim(mean=0.35, \sigma=0.1)$. Dates estimated from our nuclear datasets: 14.6-28 mya.

Age of divergence of *Ebenavia*: Normal distribution $\sim(mean=0.35, \sigma=0.1)$. Dates estimated from our nuclear datasets: 20-42 mya.

Age of divergence of *Paroedura*: Normal distribution $\sim(mean=0.35, \sigma=0.1)$. Dates estimated from our nuclear datasets: 22-39 mya.

Age of divergence of clade containing *Geckolepis* & *Blaesodactylus*: Normal distribution $\sim(mean=0.35, \sigma=0.1)$. Dates estimated from our nuclear datasets: 19-46.7 mya.

These dates for Madagascar clades are largely congruent with timing of divergences in other clade such as Gemsnakes (Burbrink et al. 2019), Vangas bird (Reddy et al. 2012), and the divergence of many dune beetle tribes (Miraldo et al. 2011). These studies inferred the start of diversification from early Miocene to Late Eocene.

18. *Gekko*-*Lepidodactylus* split: Normal distribution $\sim(mean=0.6, \sigma=0.1)$. This date largely agrees with other phylogenies of South-East Asian geckos (Heinicke et al. 2012, Siler et al. 2012). Dates estimated from our nuclear datasets: 40-71 mya.

It has been noted that the use of secondary calibration points tends to produce much younger dates (Sauquet et al. 2012, Schenk 2016). Indeed, if the secondary calibration distribution is too narrow, it may give a false sense of precision and shift the estimated age to be younger (Schenk 2016). However, we would argue that our secondary distributions are not too narrow where most of the distributions have 95% quantile values that span over 20 million years, which mostly covers, if not extends, the 95% HPD of previous studies (e.g. Agarwal et al. 2014b, Lajmi and Karanth 2020, Tamar et al. 2021). In addition, our rationale of choosing these secondary calibrations is slightly different than previous studies where secondary calibrations are used when the primary fossil evidence is absent or depauperate (See sections below). Whereas our main purpose of choosing secondary calibrations points is to reconcile nuclear dataset and full dataset with an assumption that, in deeper nodes, age estimates produced by nuclear dataset might be closer to reality due to their slower evolutionary rate and a lower propensity for saturation (Allio et al. 2017).

This is indeed the case where the date estimates of the full dataset with additional priors are not significantly different than the nuclear only date estimates (ANCOVA: *Sum Sq* > 0.2, *F* > 2.1, *p* > 0.1) except for the lower 95% HPD boundaries (ANCOVA: *Sum Sq* = 238.02, *F* > 497.68, *p* < 0.0001) (Supplementary table S3-S4). However, adding additional priors led to significantly younger date estimates for the mean, lower and upper 95% HPD compared with the full dataset with six initial priors (ANCOVA: *Sum Sq* > 39, *F* > 40, *p* < 0.0001). Overall, the full dataset with six initial priors produced the oldest age estimates across almost all nodes, for upper 95% and mean estimates.

Biogeographic regions (see also figure S5)

1) South America, north from Colombia, Venezuela to Argentina in the south. This includes islands or island nations that are close to South America continent geologically, including Leeward Antilles island nations such as Aruba and Curaçao, and Trinidad and Tobago. South America was separated from North-Central America during most of its geological history, until the formation of the Isthmus of Panama around 3 million years ago (O’Dea et al. 2016). The border between the country of Colombia and Panama closely follows the boundary between the Isthmus of Panama and South America. Therefore, it is justified to use Colombia as the start of the South America continent. We recognize that Trinidad and Tobago are sometimes classified as within Lesser Antilles in the geological literatures (Garmon et al. 2017), we follow conventions from biological literatures where they are usually excluded from Lesser Antilles islands (Henderson and Powell 2009, Losos 2009).

2) North-Central America, from USA, Mexico to Panama. This includes islands that are close to Central America geologically, such as Bay islands of Honduras. Although Central America and North America consists of many different tectonic plates and have different ways of origin, their close physical position as well as close biological connections during the last 100 million years justifies coding them as one region (Whitmore and Stewart 1965, Estes and Báez 1985, Scotese 2021). In addition, combining North and Central America can also help to reduce the total number of regions as the total combinations can get very large.

3) Caribbean, including Greater Antilles, Lesser Antilles and the Bahamas. Although the way and timing of origin of these islands varied drastically, they share some similarities in terms of their “biogeographical properties”. They are all islands of various sizes, and they form a chain extending from North America to South America through the Caribbean region. They potentially served as stepping stones for species to move between North and South America. Considering their common characteristics, we therefore code these islands in one region.

4) Africa-Arabia, including the African continent, Arabian Peninsula, and part of Iran and Turkey that is south of the Zagros Mountain ranges. The Arabian plate was part of the African continent until it split off about 25 million years ago, forming the Red Sea and Gulf of Aden (Stern and Johnson 2010), and Zagros Mountain ranges on the north. Coding Arabian and African plates together therefore reflects the long historical connection between these two regions. This region also includes Comoros and Seychelles archipelago due to geological proximity to mainland Africa continent as well as a long period of isolation (especially so in the case of Seychelles).

5) Eurasia, from Europe on the west to Japan islands on the east and Peninsular Malaysia on the south. This includes areas that are north of the Zagros Mountain ranges.

6) South Asia, or Indian subcontinent, including India, Pakistan, Bangladesh, Nepal, Bhutan and Sri Lanka. Indian subcontinent (along with Madagascar) separated from the Antarctica around 112 mya (Jokat et al. 2021) and it separated from Madagascar around 94-84 mya (Gibbons et al. 2013), becoming an isolated landmass until collision with Eurasia at around 40-50 Mya (Najman et al. 2010, Bouilhol et al. 2013). Coding South Asia as its own region reflects the unique geological history and long period of isolation of this region throughout Earth’s history. We do recognize that using country borders as geological borders between the Indian subcontinent and Eurasia in this case is only a rough proximation, but given the resolution of gecko records in these countries (or in general), we argue it would be sufficient.

7) Sundaland, includes Borneo, Sumatra, other Sunda Shelf islands and the Philippines, eastwards to the Lydekker line. Although Borneo, Sumatra, Bali and Java are part of the Sunda shelf connecting to the mainland Eurasia, they have been isolated in the Indo-Pacific Ocean for the majority part of last 200 million years, or at least been through many cycles of isolation (Scotese 2021). This is similar to conditions of other islands in Southeast Asia that have very weak or no connections to any major landmasses. Therefore, it is justified to include Borneo, Sumatra, Bali and Java as part of the Sundaland region in this study. Andaman and Nicobar Islands are also in this region due to geological proximity.

8) Australasia, eastwards of the Lydekker’s line, including New Guinea, Australia, New Zealand, New Caledonia and various islands and island nations in South Pacific Ocean, including Micronesia.

9) Madagascar. This region includes Réunion and Mauritius islands. Madagascar have separated from Africa continent around 170 to 140 mya (Rabinowitz et al. 1983, Gaina et al. 2015), and have separated from Indian subcontinent around 94-84 mya (Gibbons et al. 2013). Both events have shaped the landscape of Madagascar (Clementucci et al. 2025). These results suggest the uniqueness and prolonged isolation of this region, therefore justifying to code it as a separate region.

Biogeographic analyses

To determine the region to which a species belongs, range maps and occurrence records were viewed and integrated using The International Union for Conservation of Nature (IUCN) Red List of Threatened Species (IUCN 2025), Map of Life (Jetz et al. 2012) and publications including species descriptions, checklists and books (Supplementary material).

*Distance hypothesis*

This hypothesis states that the longer the distance between two areas, it is less likely for species to disperse between them. Distances between regions were defined as the shortest edge-to-edge distance between regions, assuming species dispersed through the potentially least resistant route. The default power function was used to modify dispersal probability in which dispersal probability (*d* parameter) was multiplied by${distance}^{x}$ (Van Dam and Matzke 2016). The power parameter *x* was set to “free” so the value can be estimated based on the data. If two or multiple regions are connected, distances among them were set to 1. Since 1 raised to the power of any number is still 1, the dispersal probability was then unchanged. The assumption here is that there were no constraints on dispersal within connected regions, that is within-region dispersal probability is equal to among-region dispersal probability. The measured distance has a unit of 100 km to prevent really large numbers (Table S5). Because regions in this case were continents, many distances among them represented the distances for direct trans-oceanic dispersals.

*Position hypothesis*

This hypothesis states that distance itself did not influence dispersal directly, rather, it is the position of the region relative to others that influences dispersal. Adjacent regions are defined as regions that were in close proximity to one another with a distance less than the distance of Africa and South America at that point in time. Many geckos are very good dispersers and there are several proposed cases of trans-oceanic dispersal in this clade (Gamble et al. 2011, Heinicke et al. 2014). Among them, there are multiple examples of trans-Atlantic gecko dispersal from Africa to South America throughout the Mesozoic and Cenozoic, some within the last 15 million years (Gamble et al. 2011). This suggests that the distance between Africa and South America did not pose a significant barrier for dispersal. We therefore use this distance as a criterion to define the manual dispersal multiplier modifiers. If regions are adjacent, we set the manual dispersal multiplier to 0.5 (Table S5).

Non-adjacent regions are defined as two regions with distance greater than the distance of Africa and South America, or with one or more regions in between. This definition suggests that species would be much more likely to use the intervening region as a “stepping-stone” when dispersing rather than direct dispersal to a non-adjacent region. For example, Indian subcontinent separated from Madagascar around 88 million years ago and therefore they became adjacent regions with ocean in between. In this case, African continent would be non-adjacent to Indian subcontinent since it is more likely for species to disperse to Madagascar first and then disperse to Indian subcontinent rather than direct dispersal from African continent to Indian subcontinent. In addition, before Indian subcontinent separated from Madagascar, they would have the dispersal multiplier were set to 1 because they are connected. However, even in this case, African continent is still considered non-adjacent to Indian subcontinent with the same reason as above. If regions are non-adjacent, we set the manual dispersal multiplier to 0.001.

We recognized that these dispersal multiplier can be very arbitrary, so we included an additional free parameter “*w*” so that the dispersal probability (*d* parameter) was multiplied with a power function ${(manual dispersal multiplier)}^{w}$, and that the *w* parameter can be estimated with data. When two regions are connected and manual dispersal multiplier is set to 1, the dispersal probability is again not scaled.

*Model Averaging*

To infer the nature of cladogenetic and anagenic events across the gecko tree, 100 biogeographic stochastic mappings (BSM, Dupin et al. 2017) was performed for each fitted phylogeny using the runBSM function within BioGeoBEARS package. To determine the input parameters of BSM, we used a model averaging approach instead of choosing the most supported model. This approach integrated parameters in each model proportional to how much the model was supported based on AICc weight ($w_{i}$).

We recognized that choosing the best model based on AICc weight values could be subjective sometimes when these values are similar to each other. In our results, even though the best and the second-best supported models are consistent across runs, the AICc weight value for them ranges from 99.7% to 62% and 0.2% to 38% respectively. Picking one best model could be fine in a few cases but model averaging would be less subjective in the majority of the cases. Therefore, for the final parameter estimates and the parameter inputs to the downstream stochastic mapping, we took the model averaging approach. To model averaging, we multiplied parameter estimates in each model with the AICc weight value of this model in model comparison, and sumed across all models. For models with fixed parameters, these parameters are left with the default values. For example, DEC model has two free parameters, dispersal (*d*) and extinction (*e*), but DEC*+j* model has three free parameters, dispersal (*d*) and extinction (*e*), and founder (*j*). If DEC model is supported with 60% of the AICc weight with $d_{1}=2$, $e_{1}=1$, $j_{1}=0$ (default value) while DEC*+j* model is supported with 40% of the weight with $d_{2}=1$, $e_{2}=0.5$, $j_{2}=1$, the final values would be: $d_{final}=2\times0.6+1\times0.4=1.6$; $e_{final}=1\times0.6+0.5\times0.4=0.8$, and $j_{final}=0\times0.6+1\times0.4=0.4$. This way, the influences of all models could be accounted for. The model averaged parameter estimates for each model of each run can be found in table S6.

Integrating fossils into biogeographic analyses

The gekkotan fossil record is sparse due to the small size, fragility, and isolated nature of many squamate paleoassemblages (Daza et al. 2014). In addition, phylogenetic studies including gekkotan fossils are rare, and relationships among stem taxa are unresolved (Daza et al. 2013b, but see Meyer et al. 2023). Nevertheless, some fossils can be confidently assigned to extant genera, especially for those restricted to the Neogene (Daza et al. 2014).

Fossil lineages can be used in the biogeographic analyses because they have distribution data associated with them. Although most of the time we do not know who their closest related living species is, sometimes we have a fairly good estimate of which taxonomic group it resides in or closely related to. Therefore, we picked gecko fossil lineages that can confidently assign to extant clades or have previously been included in phylogenetic analysis.

Two Cretaceous fossils, *Hoburogekko suchanovi* Alifanov (1989) and *Gobekko cretacicus* Borsuk-Białynicka (1990), were included in phylogenetic analyses in previous studies and were found to be part of a basal polytomy outside of crown Gekkota (Daza et al. 2013b). Recent phylogenetic hypotheses, however, showed that *Gobekko cretacicus* could possibly be sister to Pygopodoidea (Meyer et al. 2023). While interesting, this study implemented very sparse sampling of crown geckos and returned a morphological phylogenetic hypothesis with numerous relationships not recovered in any molecular study (Burbrink et al. 2020, Singhal et al. 2021). Therefore, in this study, we conservatively placed *G. cretacicus* outside of the crown Gekkota following (Daza et al. 2013b). Finally, *Cretaceogekko burmae* (Arnold and Poinar 2008) can be assigned to Gekkota but could not confidently placed to any extant families. We therefore treated this species to also be outside of the crown Gekkota as well (Detailed reasons below). We included four additional fossil lineages that could be confidently assigned to extant genera: *Euleptes gallica* (Müller 2001), *Pygopus hortulanus* (Hutchinson 1997), *Sphaerodactylus dommeli* (Böhme 1984), and *Sphaerodactylus ciguapa* (Daza and Bauer 2012). In the end, a total of seven fossil lineages were included (Table S7).

To attach fossil to the phylogenetic tree, we use the bind.tip() function in the R package phytools 2.5-2 (Revell 2012). To attach stem fossil lineages outside the crown gecko clade, we first attached *H. suchanovi* to the root node, sister to all crown gecko lineages. Both the starting time (the attach point) and the terminating time of this branch are sampled from a uniform distribution, bounded by the lower and upper time boundary of the formation(s) in which the fossil was found, following the latest version of GSA geologic time scale (Walker and Geissman 2022), or by the time ranges estimated in the original article of description. We then sequentially attached two other stem fossil lineages *Cretaceogekko burmae* and *Gobekko cretacicus* to the phylogeny. Firstly, we randomly picked a node outside of the crown gecko, which could be a previously attached fossil tip(s), the node of the crown geckos, or any node that was produced by previous attachments. We then attached the fossil lineage to the branch leading to this node at a random point sampled with a uniform distribution. This uniform distribution is bounded by the age of the chosen node, and the age of the parent node of the chosen node. Again, the terminating time of this branch are sampled from a uniform distribution bounded by the upper and lower boundary of the formation(s) in which the fossil was found, or by the time ranges estimated in the original article of description. For fossil lineages that could be confidently assigned to extant genera, we attached them to the most inclusive clades to which they could be assigned with confidence. We first identified a set of nodes within the clade that were older than the minimum age of the relevant deposit. Then we selected a random node within this node set and attached the fossil lineage to the branch leading to this node at a random point sampled with a uniform distribution. Similarly, this uniform distribution is bounded by the age of the chosen node, and the age of the parent node of the chosen node. To make sure the fossil branches are positive in length and to prevent very short branches, we constrained all fossil branches so that their starting time must be greater than the terminating time and that the branch length should be greater than 1 million years.

Overall, the geological distribution of the fossils that could be assigned to extant genera are largely consistent with the geological distribution of the genus (*Euleptes* and *Pygopus*), so adding them randomly did not produce alternative geographical hypotheses, at least for the clade in which they are nested. Within the genus *Sphaerodactylus*, adding fossil *Sphaerodactylus dommeli* could potentially produce different hypotheses and may increase estimated dispersal events in the phylogeny. This is because *S. dommeli* can be placed into *Sphaerodactylus* *sensu lato*, but have not confirmed to have morphological similarities to any extant species or clades (Daza et al. 2013a). *Sphaerodactylus ciguapa*, on the other hand, was suggested to have phenetic similarities to extant Hispaniolan species *S. difficilis* and *S. shrevei* (Daza and Bauer 2012). Even though there is not enough evidence to put *S. dommei* into any extant *Sphaerodactylus* lineage, it was found in the same geological area and time period as *Sphaerodactylus ciguapa* (Daza et al. 2013a, Daza et al. 2014)*.* Therefore, for biogeographic analyses, we restricted *S. dommei* to being in the same group as *S. ciguapa*, which is a group found in Hispaniola. To attach these two fossils to the phylogeny and avoid alternative geographical hypotheses, we restricted them to the Caribbean clade of *Sphaerodactylus*. Even though there is one colonization event back to Central America (*S. rosaurae* and *S. alphus*), it is much younger than both fossils that the fossil will never get attached to the branch leading to this node.

It should be noted that some fossils were used both as a calibration prior in the node time calibration analyses and as a fossil branch attached to the phylogeny in the biogeographic analyses. Due to the randomness of attaching fossils to the phylogeny, this can sometimes create seemingly conflicting hypotheses. For example, we used *Cretaceogekko burmae* as a calibration prior for the crown Gekkota, essentially treating it as a crown gecko species. However, in the biogeographic analyses, we attached it to the stem lineages, treating it as a stem species. But ultimately, we do not know where it falls due to the lack of osteological features of this fossil. Therefore, it is reasonable to explore how it affects crown Gekkota range reconstruction when *C. burmae* is a stem species, assuming all the divergent time estimates are correct. We avoided attaching *C. burmae* to any extant lineages to avoid producing unnecessary alternative geological hypotheses within extant clades, which could potentially cause more harm than good. Essentially, we wanted to introduce variance only on the node of crown Gekkota, but not too much variance to drown out the signals within the crown clades, which are the signals we are looking for. The same reasoning applies for constraining *Sphaerodactylus dommeli* to be within the Caribbean clade of the genus *Sphaerodactylus*. Similarly, we can explore the effect of including fossils to the overall ancestral state reconstruction without producing unnecessary noise on the transition rates and event frequencies.

Additional null models

It has long been recognized, perhaps very intuitively, that the best supported model might not be the “correct” model if the “correct” model is not even included in the comparison (Beaulieu and O'Meara 2016). Furthermore, as Beaulieu and O’Meara 2016 pointed out, rejecting the simpler model does not mean that all the assumptions of the more complex model are reasonable. It might very well be the case in our study where we compared models without time-stratification (DEC and DEC*+j*), against models with time-stratification (model included modifier matrices). Because of the over simplicity of these non-time-stratified models, it might be expected that they will have very low support almost all the time, failing to compete with the time-stratified models. Therefore, the low support of the non-time-stratified models does not tell us anything about the validity of including the modifier matrices or whether the entry of these matrices fit better than some random numbers. To try to create null models with same parameter numbers, in other words, same complexity, and to test if the entry of our modifier matrices explains data better than some random numbers, we created distance matrices and manual dispersal multipliers matrices populated with random numbers.

To create random distance matrices, we first fit all the distances in our original distance matrices to a set of univariate distributions using *model_select* function in r package univariateML (Moss 2019) to determine the best distribution that fit the data. Then we generated random numbers using the best-fitted distribution. We also tallied the number of ones (meaning two regions are connected without any constraints to dispersal) in our original distance matrices. Together with the randomly generated distance, they are put randomly into a new distance matrix for each time periods. The procedure for generating random manual dispersal multipliers is similar except that we generated multiplier numbers (0.001, 0.5, and 1) based on their proportion within the original matrices. We then fitted the same models as in the main text, except this time we used these randomly generated matrices. We again compared all models using AICc criteria.

The null models with randomly generated matrices performed significantly worse compared to models with nonrandom matrices (table S9, figure S13). However, the null model with the highest parameter number (DEC*+j*_ran_wx model: with jump dispersal and both random distance and manual dispersal multipliers matrices) performed three or four magnitudes better than simple DEC model, and performed a little worse, but comparable, to the DEC*+j* model as well.

We concluded that our original, nonrandom distance and manual dispersal multipliers matrices explain data better than some random numbers, showing that our original matrices could represent some biological realities. In addition, we concluded that the simpler DEC and DEC*+j* model failed to compete with more complex models in model comparison situation with AICc weights as criteria, even when the matrices in the complex models are filled with random numbers. This suggests that these simpler models will likely always get rejected when more complex models are included, therefore rejecting them does not provide much more valuable information. Of course, whether this conclusion can be generalized to broader phylogenetic tree topologies and biological systems requires more empirical and simulation studies.

Some taxonomic notes

For family-level relationships, the Pygopodoidea is sister to all other geckos. Within Pygopodoidea, Diplodactylidae is sister to Carphodactylidae and Pygopodidae, which is also supported by both coalescent (Skipwith et al. 2019, Singhal et al. 2021) and concatenated approaches using genome-wide data (Title et al. 2024). In our phylogeny, Eublepharidae is sister to the clade uniting Sphaerodactylidae, Phyllodactylidae and Gekkonidae. This relationship was not supported by coalescent-based topologies in Singhal et al. (2021), or by both coalescent and concatenated based topologies using genome-wide data (Streicher and Wiens 2017, Burbrink et al. 2020, Title et al. 2024: topology produced by their constrained approach). Interestingly, our three-gene (including the loci ND2, C-mos, and pdc) supported the topology where Eublepharidae is sister to all geckos while adding more loci returned the topology where Eublepharidae is sister to Sphaerodactylidae, Phyllodactylidae and Gekkonidae. (Supplementary material). This suggests that at least some loci supported the former relationships but most of them supported the latter relationships. Nonetheless, considering the limitation of our methodology (e.g. we only considered concatenation), the relationship of Eublepharidae with other gecko families still needs further study and may be unresolvable to existing genetic data and analytical methods.

Our topology supported monophyly for the majority of genera with some noticeable exceptions. Within New Caledonian diplodactylids, *Mniarogekko chahoua* returned to be nested within *Eurydactylodes* and was sister to *E. symmetricus*. This contrasts to previous large-scale Diplodactylodae phylogeny (Skipwith et al. 2019) which used ultraconserved elements (UCEs). They supported *Mniarogekko* as nested within *Rhacodactylus* species, which are morphologically similar to *Mniarogekko* (Bauer 1990). They argue that the close association between *Mniarogekko* and *Eurydactylodes* is driven by mitochondrial genome. Another large-scale supermatrix phylogeny that included traditional locus sets returned a sister relationship between *Mniarogekko* and *Eurydactylodes* (Title et al. 2024), further supporting this claim. We believe that the relationships in Skipwith et al. 2019 are probably more reasonable and reliable.

Another paraphyly in our phylogeny is regarding the *Strophurus taenicauda*, which is nested within clade *Amalosia* and is sister to *A. robusta*. Same relationships were returned across other supermatrix studies (Zheng and Wiens 2016, Title et al. 2024). This is not a surprise since similar data were used to construct our phylogeny as well. This spurious position of *S. taenicauda* is likely stemmed from a single mitochondrial ND2 sequence (NCBI genbank accession number: AY369006). This sequence is present in all of the above datasets, including ours (from Zheng and Wiens 2016 dataset where it is the only sequence sampled for this species). In its original paper (Melville et al. 2004), *S. taenicauda*, represented only by this sequence, is supported to be outside of the *Strophurus*, although molecular data did not reject the possibility of it being inside of the genus. Title et al. 2024 included two additional sequences (RAG1 and 16S) from Nielsen et al. (2016) for this species, which supported the placement of *S. taenicauda* to be within *Strophurus* in their original paper. However, Title et al. 2024 still returned the same sister relationship of *S. taenicauda* and *A. robusta*, suggesting this ND2 sequence potentially masked the effects of these two other sequences. This points to the potential limitations of the supermatrix approach that one or few spurious sequences in certain species can lead to erroneous relationships. Therefore, nesting *S. taenicauda* within *Amalosia* should be considered spurious without further evidence.

Within Sphaerodactylidae, *Saurodactylus fasciatus* is sister to *Euleptes europae*. It has been recognized that *Saurodactylus* is not monophyletic (Rato and James Harris 2008) and was supported by another study with different sets of sequences (Gamble et al. 2012, but see Gamble et al. 2015). In addition, the morphology of *S. fasciatus* is somewhat distinct from all other *Saurodactylus* gecko (Javanmardi et al. 2019), providing additional evidence for the paraphyly of this genus. We do believe that this could be a true paraphyly and it is currently being resolved (Daza et al. in prep).

Within the Gekkonidae, the relationships among *Lepidodactylus*, *Luperosaurus*, and *Pseudogekko* have long been complicated (Brown et al. 2012, Oliver et al. 2018). The majority of our sequences for these groups came from a few previous studies (Heinicke et al. 2012, Oliver et al. 2018, Eliades et al. 2021), but there are some novel relationships that are worth noting. *Lepidodactylus ranauensis* is sister to *Lepidodactylus* *lugubris* group + *L. guppyi* group + *L. listeri* + *Luperosaurus* + *Pseudogekko*, instead of to the *L. novaeguineae* group (Oliver et al. 2018). This suggests that *L. ranauensis*, which inhabits Mount Kinabalu in Sabah, Malaysia, is sister to other species of this taxonomic complex that occupied similar biogeographic region, namely Sundaland plus the Philippines. Our phylogeny also suggests a sister relationship of *Luperosaurus joloensis* + *Gekko* (*Luperosaurus*) *sorok* to *L. listeri* instead of to the *Pseudogekko* (Oliver et al. 2018, Fukuyama et al. 2022). This in turn suggests a closer grouping of these two *Luperosaurus* species with the *L. lugubris* group which, like *Pseudogekko*, also found across the Philippines. Interestingly, our reduced representative (4-gene and 3-gene) datasets do support the previous relationship, but it changes to the current relationship when more genes are added (e.g. 16S in 5-gene dataset, Supplementary material). Previous UCE phylogeny found similar groupings with some important exceptions (Wood et al. 2020). In their phylogeny (Figure 2 and 3), *Luperosaurus cumingii* is sister to *Lu. joloensis* while *Lu. cumingii* is grouped with *Lu. angliit* + *Lu. macgregori* in our phylogeny. This is partially supported by the biogeographic evidence where the species of *angliit*, *macgregori*, and *cumingii* all occur in the northern and northeastern Philippines on the island of Luzon and its offshore islands. On the other hand, the species of *Lu. joloensis* and *sorok* are found in the southern Philippines island of Jono and in the Malaysian Borneo respectively, which share a close geographic proximity as well. Again, biogeographic proximity does not necessarily warrant a phylogenetic proximity, especially in geckos where trans-oceanic dispersals are relatively common. Nonetheless, we believe our phylogeny hypothesis provides a more parsimonious biogeographic scenario and therefore comes with some support.

The relationships among Palearctic naked-toed geckos are problematic and poorly resolved despite decades of hard work among taxonomists (Bauer et al. 2013, Machado et al. 2019). Our phylogeny certainly does not provide a definitive answer to this problem, instead it provides yet another potential taxonomic hypothesis. All genera are monophyletic except *Cyrtopodion*, *Bunopus*, and *Tropiocolotes*. *Cyrtopodion* are being split into three groups. One is the ‘*Cyrtopodion*’ *aravallense* group (Agarwal et al. 2014a) which is sister to the clade including *Crossobamon*, *Bunopus*, and *Rhinogecko*. Second group consisting of *C. gastrophole*, *C. agamuroides* and *C. persepolense* (Hosseinian Yousefkhani et al. 2019) that is sister to *Tenuidactylus*. Lastly, the third group consists of the rest of the *Cyrtopodion* and it is sister to the second group + the *Tenuidactylus*. Looking at out reduced representative datasets, in all of them, second and third group become sisters and together is sister to *Tenuidactylus*. *Crossobamon eversmanni* is nested within *Bunopus* and is sister to *B. blanfordi* + *B. tuberculatus*. However, in a later study with broader sampling both in terms of number of loci and species distribution, *C. eversmanni* is sister to all *Bunopus* (Pola et al. 2024), even though with relatively low support on the node. Despite the fact that the topology of Pola et al. (2024) is likely more reasonable, the monophyly of *Bunopus* is still probably problematic because our phylogeny supports *Rhinogecko misonnei*, which was not included in the previous paper, to be nested within *Bunopus*, sister to *B. crassicauda* + *B. orientali*. This nesting of *Rhinogecko* within *Bunopus* is supported in all our reduced representative datasets, suggesting the result might be somewhat robust. Lastly, *Tropiocolotes algericus* does not group with the rest of the *Tropiocolotes* species but is sister to them plus *Trigonodactylus* and *Stenodactylus*. Again, all of our reduced representative datasets support this topology. Most of our sequences for *Tropiocolotes* came from Machado et al. (2019) which specifically addressed the question of monophyly of this genus. They indeed found a support for the monophyly of the genus despite with some uncertainty. It is perhaps not a surprise that increase taxon sampling is important for accurately infer and resolve the relationships (Hedtke et al. 2006, Wiens and Tiu 2012), but whether our topology represents the truth or not still requires a lot more scrutiny.

Another genus worthy of note is *Calodactylodes*. This South Asian lineage is nested within an Africa-Madagascar clade including *Urocotyledon*, *Ebenavia*, and *Paroedura* with high bootstrap support value (BS=90). This nesting is supported with high values across reduced gene topologies as well (BS>80) (supplementary material). These species all possess similar external toepad morphologies where they are leaf-padded with a single or double (only in *Calodactylodes*) row(s) of terminal pads. This also suggests a dispersal event from Africa to the Indian subcontinent. This is not unreasonable given the close proximity of these two land masses throughout history, and other inferred dispersal events from Africa to India within the genus *Eublepharis* and from Madagascar to India in the ancestor of the South Asian *Cnemaspis* clade (Figure S7, S8).

Figure S7. Dated phylogeny returned from BEAST2 analysis with full + moreprior dataset. Reconstructed ancestral areas with time-statified DEC+j+w+x model fitted to the phylogny and with fossils randomly attached to the phylogeny are summarized across ten runs and are plotted on nodes. Fossil taxa are removed before plotting. The texts on the nodes have the format of “maximum likelihood state: number of runs that this state is the maximum likelihood state”. For example, A: 6 B: 4 means that in six out of ten runs, state A is the maximum likelihood state for this node, while in the other 4 runs, state B is the maximum likelihood state for this node. State coding: A-Africa-Arabia; B-Madagascar; C-Eurasia; D-South Asia; E-Sundaland; F-Australasia; G-Carribean; H-North-Central America; I-South America.

Figure S8. Dated phylogeny returned from BEAST2 analysis with full + moreprior dataset. Reconstructed ancestral areas with time-statified DEC+j+w+x model fitted to the phylogeny and without fossils randomly attached to the phylogeny are summarized across ten runs and are plotted on nodes. The texts on the nodes have the format of “maximum likelihood state: number of runs that this state is the maximum likelihood state”. For example, A: 6 B: 4 means that in six out of ten runs, state A is the maximum likelihood state for this node, while in the other 4 runs, state B is the maximum likelihood state for this node. State coding: A-Africa-Arabia; B-Madagascar; C-Eurasia; D-South Asia; E-Sundaland; F-Australasia; G-Carribean; H-North-Central America; I-South America.

Figure S9. Dated phylogeny returned from BEAST2 analysis with full + moreprior dataset. Both cladogenetic and anagenetic events realized in Biogeographic Stochastic Mapping (BSM) analysis with model-averaged parameters and with fossils randomly attached to the phylogeny are summarized across 100 BSM maps each run across ten runs, and are plotted on nodes and branches. Fossils are removed prior to plotting. A certain type of events is saved in each run only if they happened in more than 80 out of 100 BSM maps in this run. Vicariance events are saved if they happened in more than 60 out of 100 BSM maps. We also filtered out events that happened in less than 10 out of 100 BSM maps in each run. The events on the nodes and branches have the format of “event text: (average number of BSM maps in each run that this event happened in more than 80 BSM maps)/(number of runs this event happened in more than 80 BSM maps)”. For example, CF−>CF,F: 80.5/4 means that a subset type of event happened in this node where one of the daughter lineages inherits the ancestral range CF while another daughter lineage inherits part of the ancestral range F. And this event happened averaged 80.5 times across four runs that it happened in more than 80 BSM maps. Relevant node states are also plotted on the nodes and these are summarized from BSM analyses rather than ancestral area reconstructions. For anagenetic events, even though our model does not allow direct range switch (parameter a in BioGeoBEARS is fixed to be 0), but a range switch type event can still happen by a range expansion event followed by a range contraction event. For example, C−>CD; CD−>D means that the species expand to include state D as its range along with its ancestral range C, but later a range contraction events led to the loss of its ancestral range C, resulting in the range that is different than its ancestral range, thus a range switch. For events for the Crown node, refer to Figure 3 in the main text. State coding: A-Africa-Arabia; B-Madagascar; C-Eurasia; D-South Asia; E-Sundaland; F-Australasia; G-Carribean; H-North-Central America; I-South America.

Figure S10. Dated phylogeny returned from BEAST2 analysis with full + moreprior dataset. Both cladogenetic and anagenetic events realized in Biogeographic Stochastic Mapping (BSM) analysis with model-averaged parameters and without fossils randomly attached to the phylogeny are summarized across 100 BSM maps each run across ten runs, and are plotted on nodes and branches. A certain type of events is saved in each run only if they happened in more than 80 out of 100 BSM maps in this run. Vicariance events are saved if they happened in more than 60 out of 100 BSM maps. We also filtered out events that happened in less than 10 out of 100 BSM maps in each run. The events on the nodes and branches have the format of “event text: (average number of BSM maps in each run that this event happened in more than 80 BSM maps)/(number of runs this event happened in more than 80 BSM maps)”. For example, CF−>CF,F: 80.5/4 means that a subset type of event happened in this node where one of the daughter lineages inherits the ancestral range CF while another daughter lineage inherits part of the ancestral range F. And this event happened averaged 80.5 times across four runs that it happened in more than 80 BSM maps. Relevant node states are also plotted on the nodes and these are summarized from BSM analyses rather than ancestral area reconstructions. For anagenetic events, even though our model does not allow direct range switch (parameter a in BioGeoBEARS is fixed to be 0), but a range switch type event can still happen by a range expansion event followed by a range contraction event. For example, C−>CD; CD−>D means that the species expand to include state D as its range along with its ancestral range C, but later a range contraction events led to the loss of its ancestral range C, resulting in the range that is different than its ancestral range, thus a range switch. For events for the Crown node, refer to Figure 3 in the main text. State coding: A-Africa-Arabia; B-Madagascar; C-Eurasia; D-South Asia; E-Sundaland; F-Australasia; G-Carribean; H-North-Central America; I-South America.

Figure S25. Ancestral state probability reconstructions on the node for the MuHisse model in RevBayes. The pie charts on the nodes are the proportion of the probability for each estimated states for the nodes. Hidden states are 0 and 1 while observed states are from A to I. State coding: A-Africa-Arabia; B-Madagascar; C-Eurasia; D-South Asia; E-Sundaland; F-Australasia; G-Carribean; H-North-Central America; I-South America.

Figure S26. Ancestral hidden state probability reconstructions on the node for the MuHisse model in RevBayes. The pie charts on the nodes are the proportion of the probability for each estimated states for the nodes. Hidden states are 0 and 1.

Figure S23. Speciation rates across geckos estimated by BAMM (shift prior = 2) on time-calibrated tree (UCE-supermatrix MCC) displaying all 1,783 tips (see Fig. 4A in the main text). Branch color corresponds to rate and red dots approximate where shifts in speciation occurred.

Figure S24. Ultrametric phylogeny of 1,783 gecko species with node numbers listed. Meant to be used in conjunction with table S2.

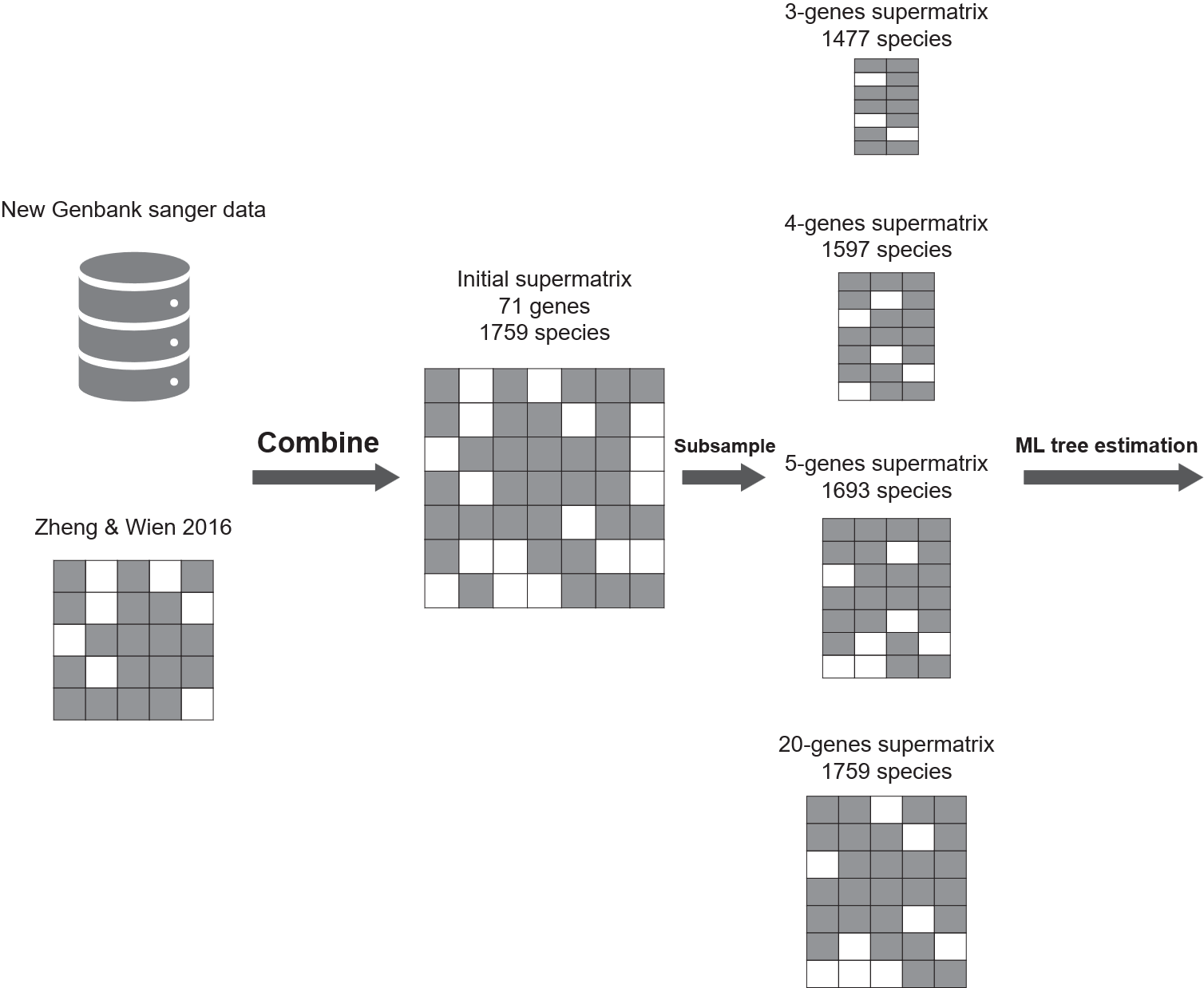

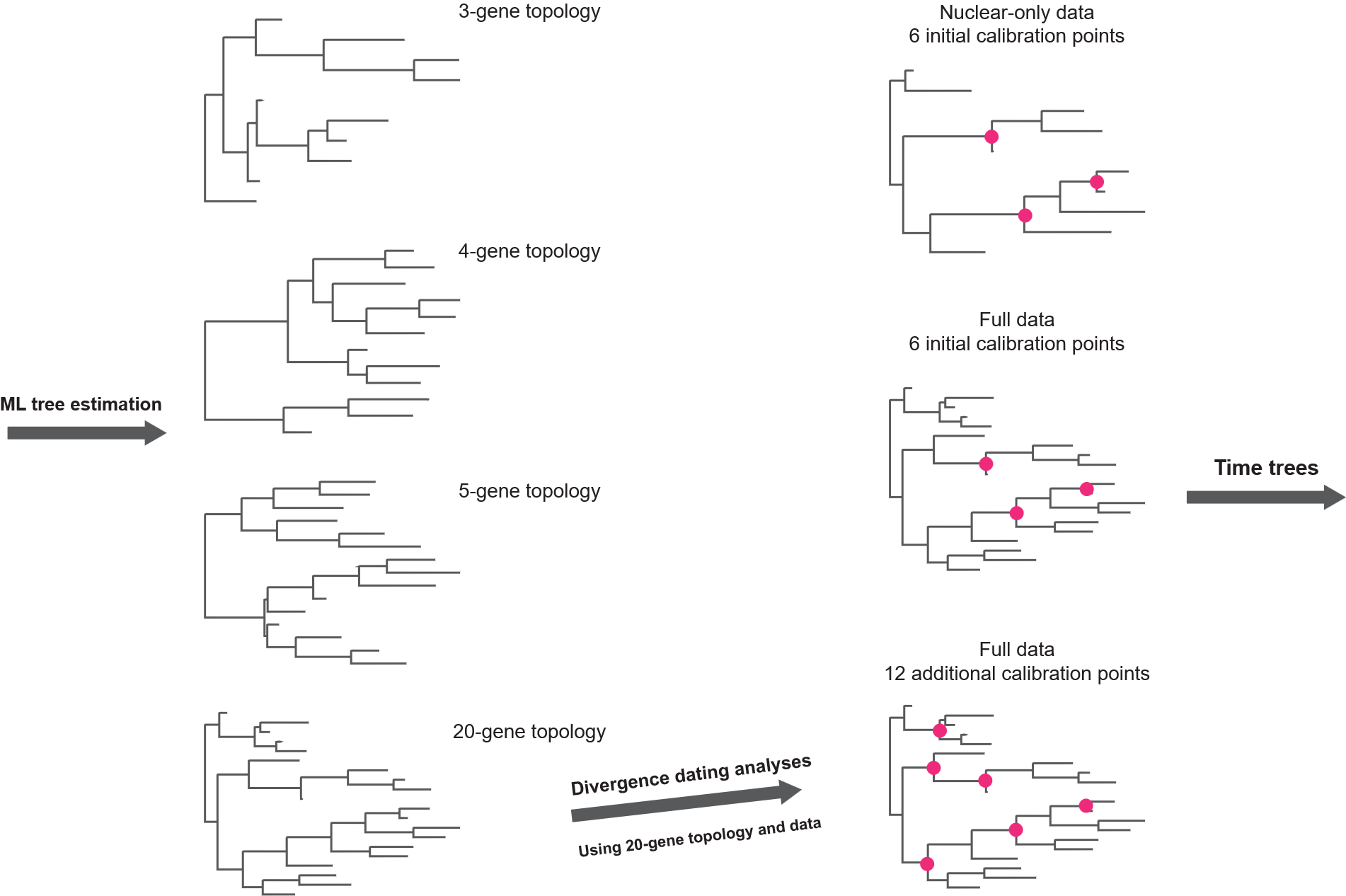

**A**

**B**

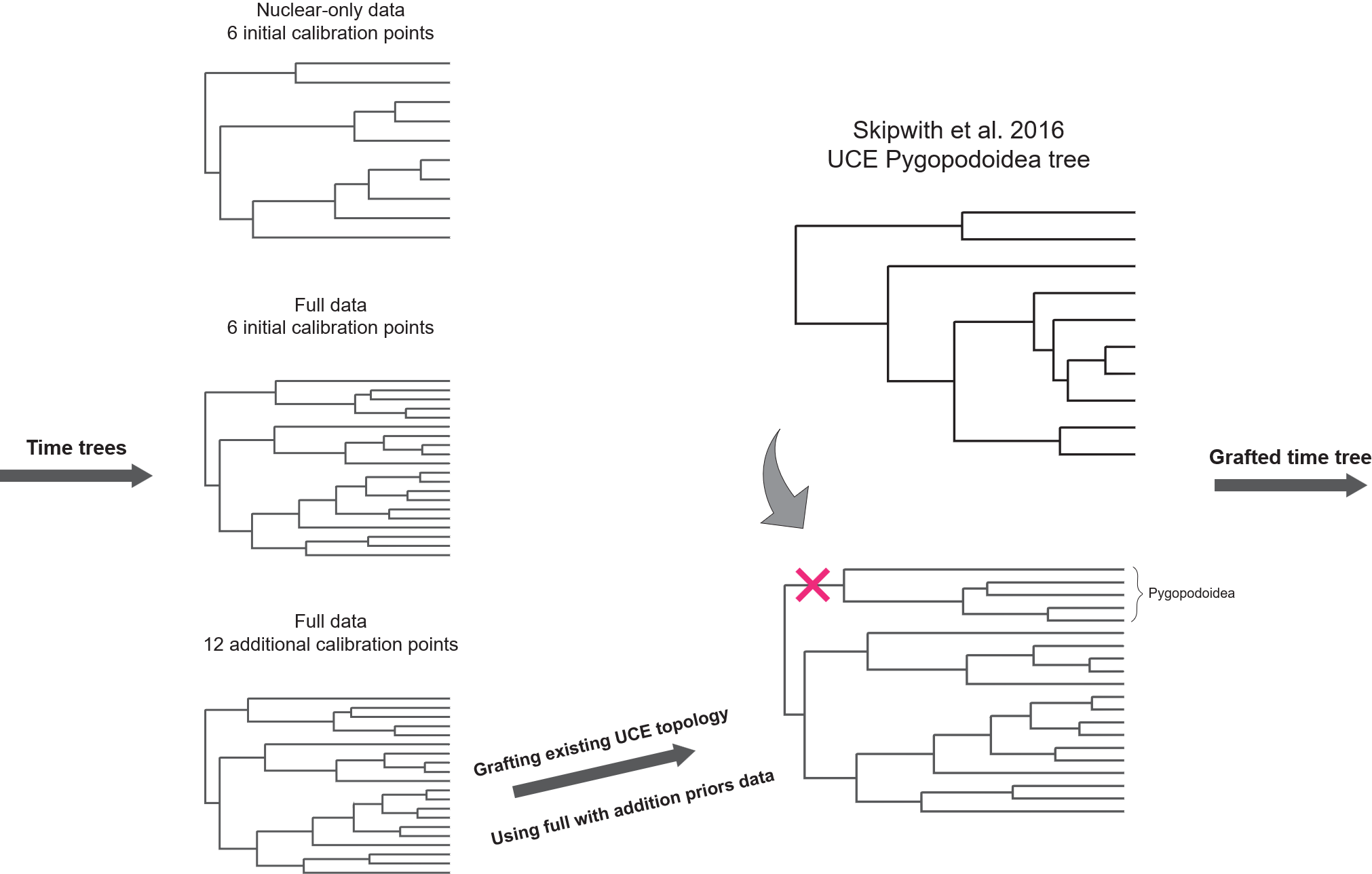

**C**

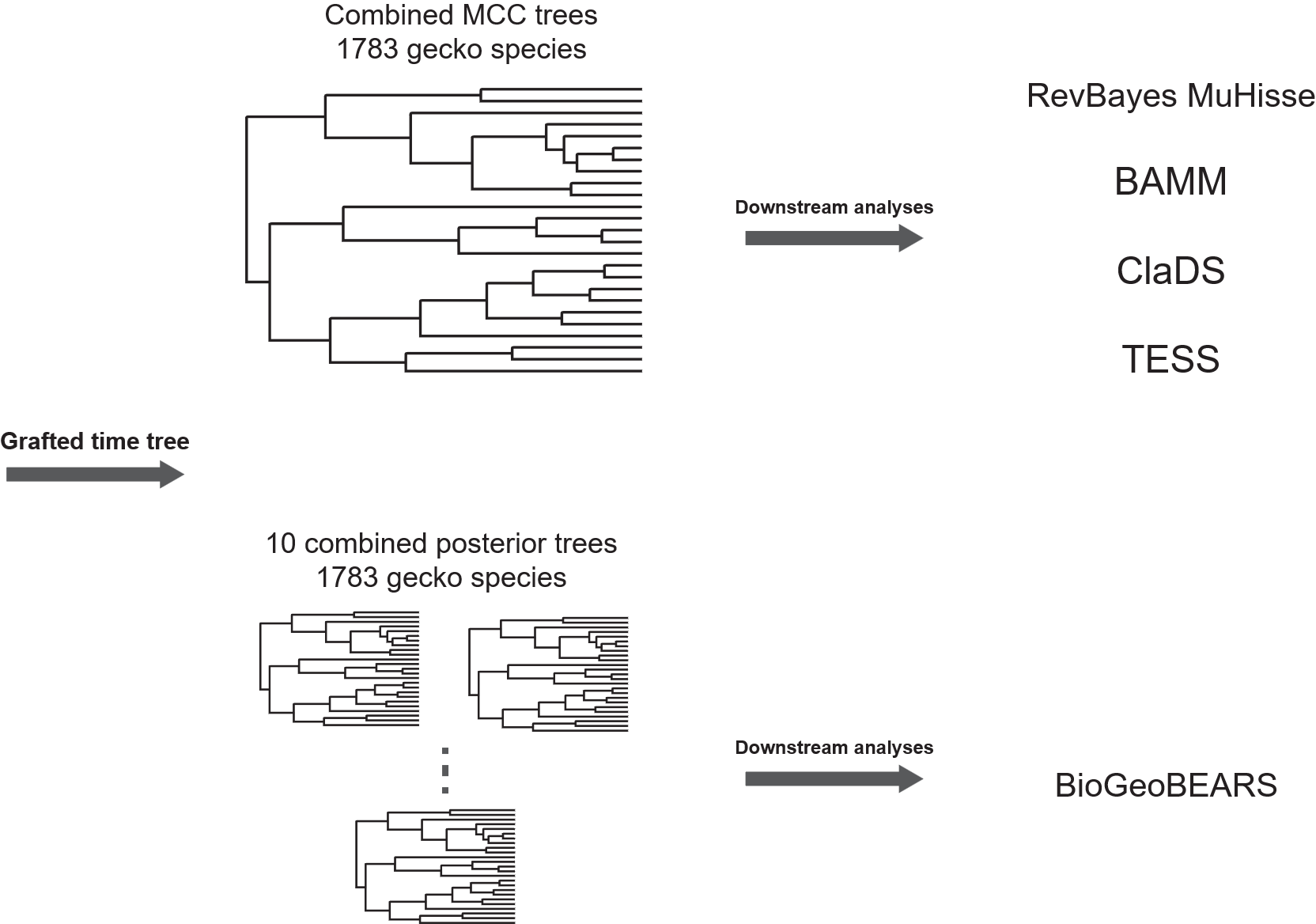

**D**

Figure S1. Flow chart describing the procedures and methods used in this study.

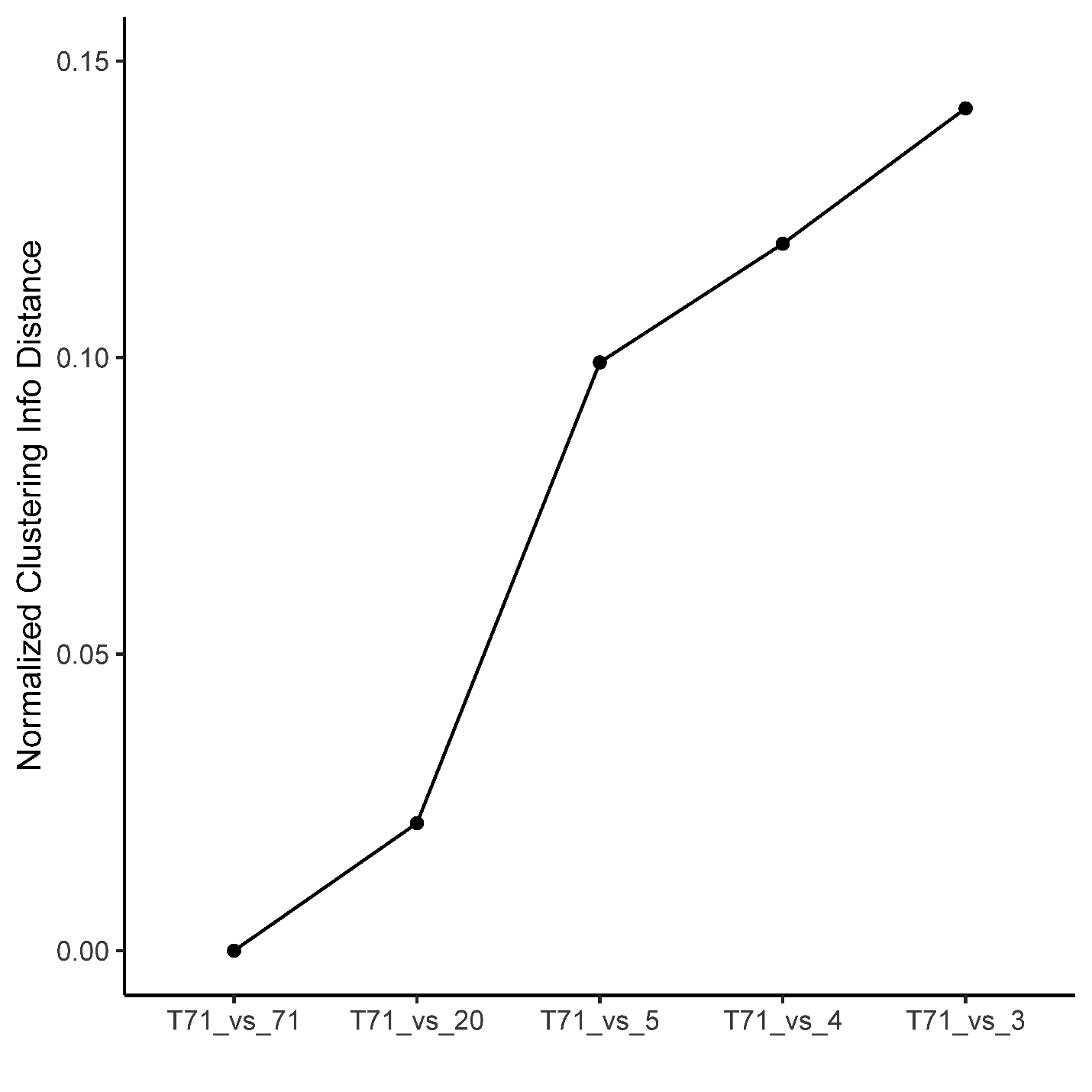

Figure S2. Information-based generalized Robinson–Foulds distances based on clustering information. “T71_vs_71” means 71-gene MASCE topology compared with itself. “T71_vs_20” means 71-gene MASCE topology compared with 20-gene MASCE topology, and so on. Y axis is normalized clustering distance which is based on the clustering information criteria in (Smith 2020). The value is normalized against the maximum which is defined as half of the information content of all the splits in both trees (Smith 2020). To control for species missingness, only species present in the 3-gene dataset were kept when comparing across datasets.

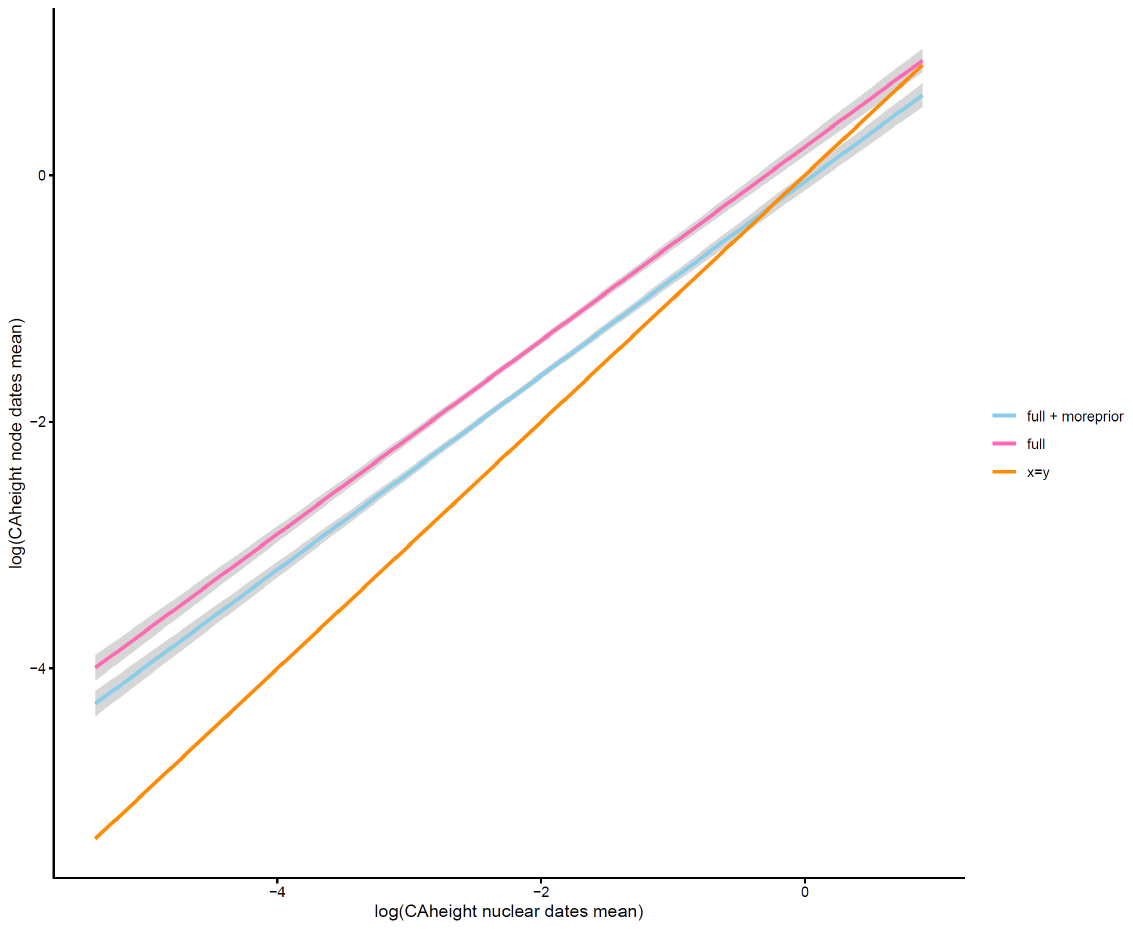

**A**

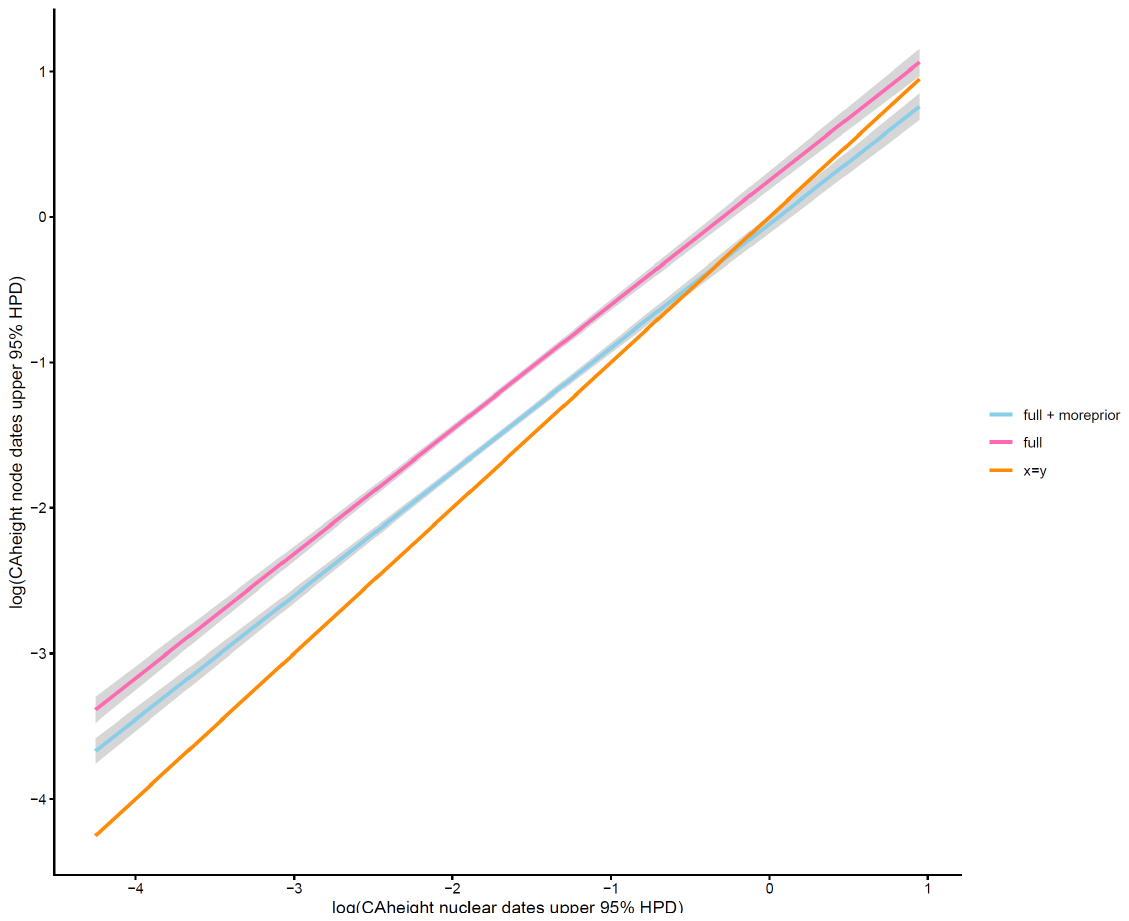

**B**

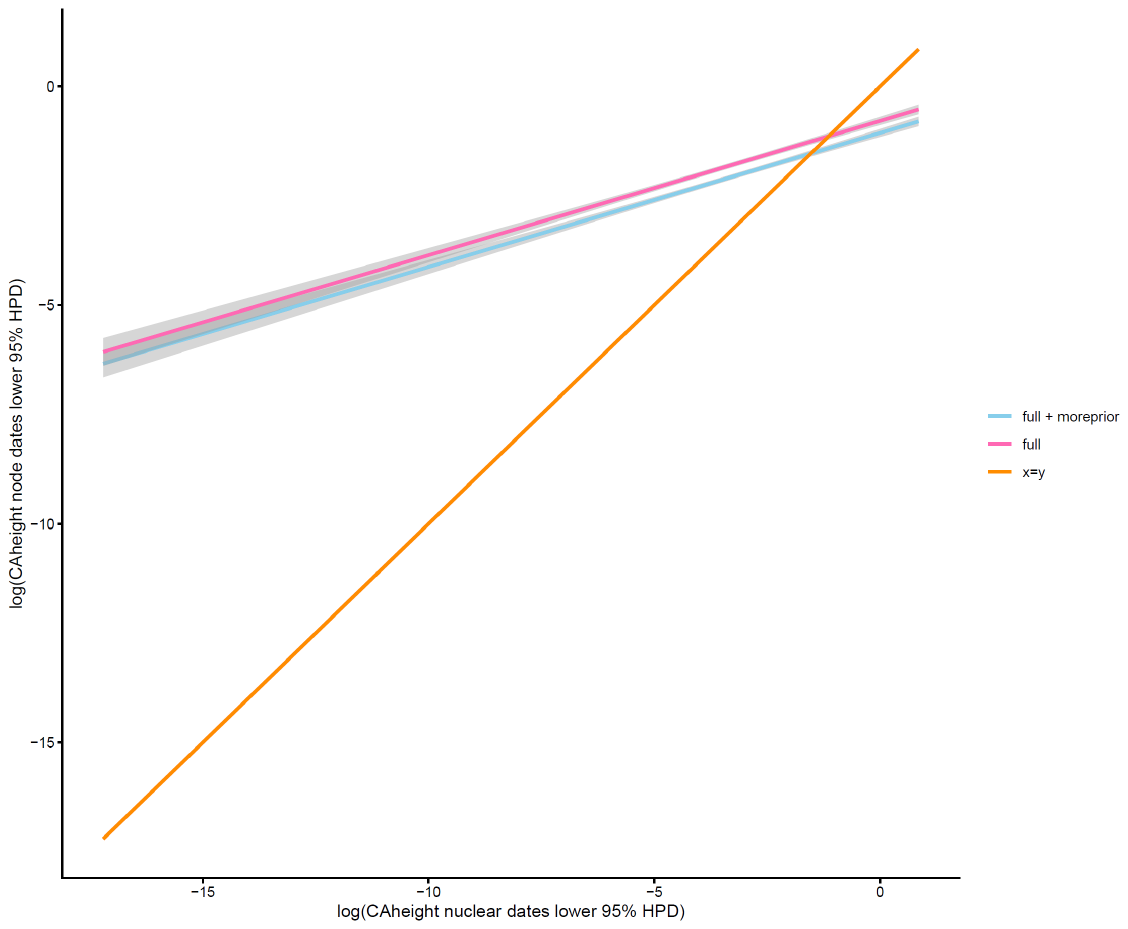

**C**

Figure S3. Comparisons of the node age estimations (A: mean estimates; B: upper 95% HPD estimates; C: lower 95% HPD estimates) between nuclear only datasets and datasets including both nuclear and mitochondrial data with initial six prior (full) and with additional twelve priors (full + moreprior). Full: dataset with initial 6 node calibration points but allowing each partition to have different clock rates. Full + Moreprior: dataset with 18 calibration points and allow each partition to have different clock rates. Node age estimations were log transformed before plotting. The lines represent the fitted linear model with formula y ~ x using “lm” method within the *geom_smooth()* function of ggplot 2.

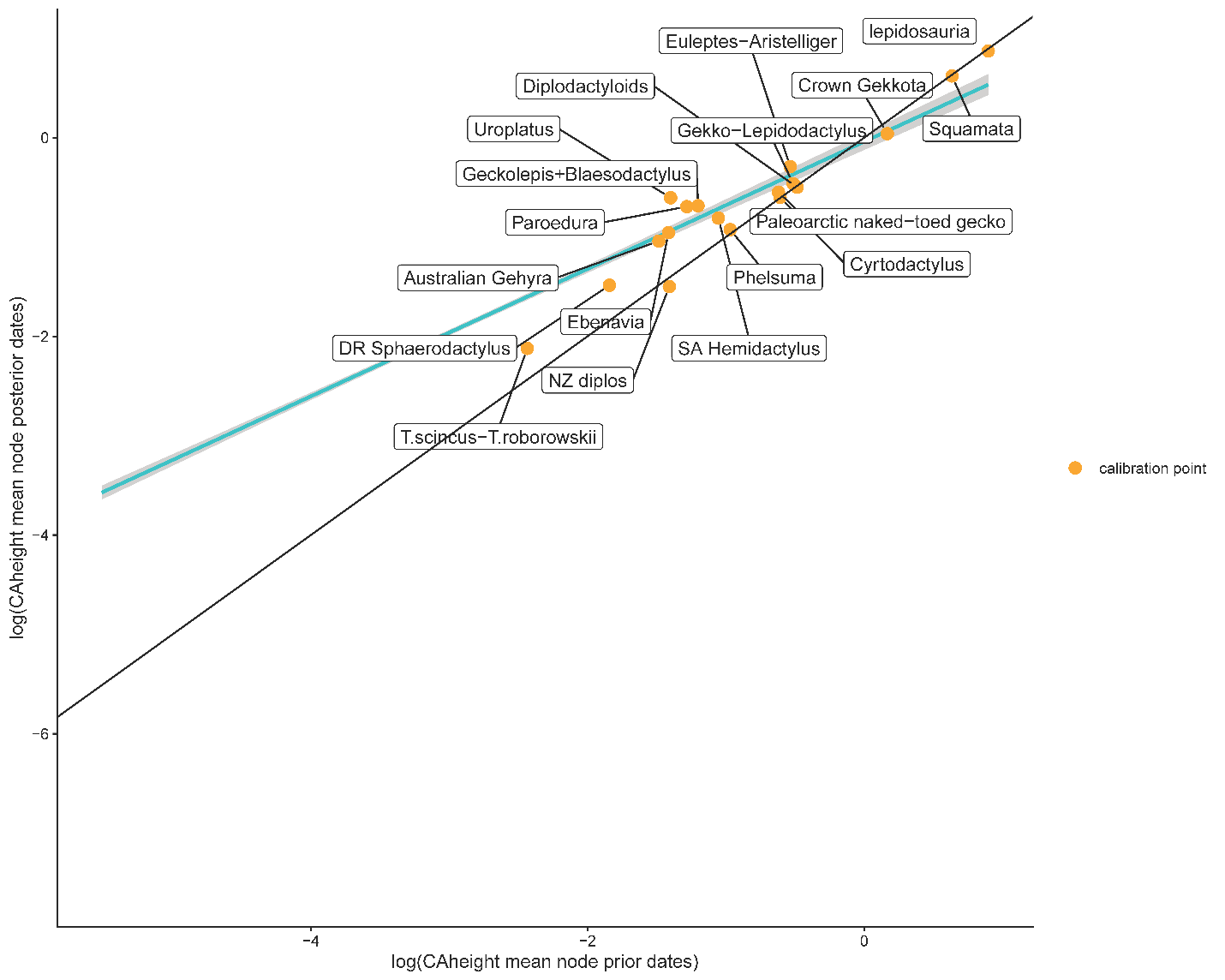

**A**

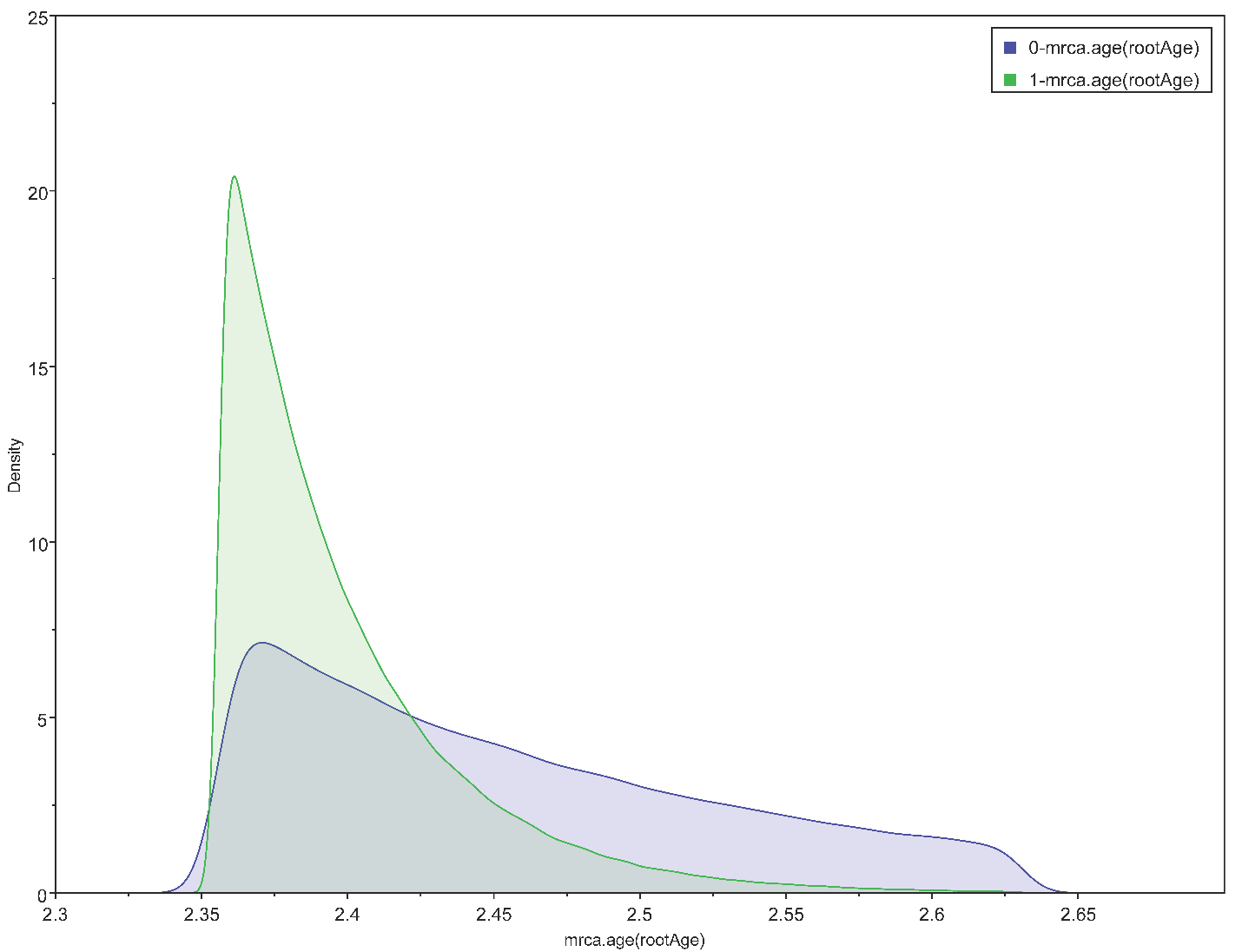

**B**

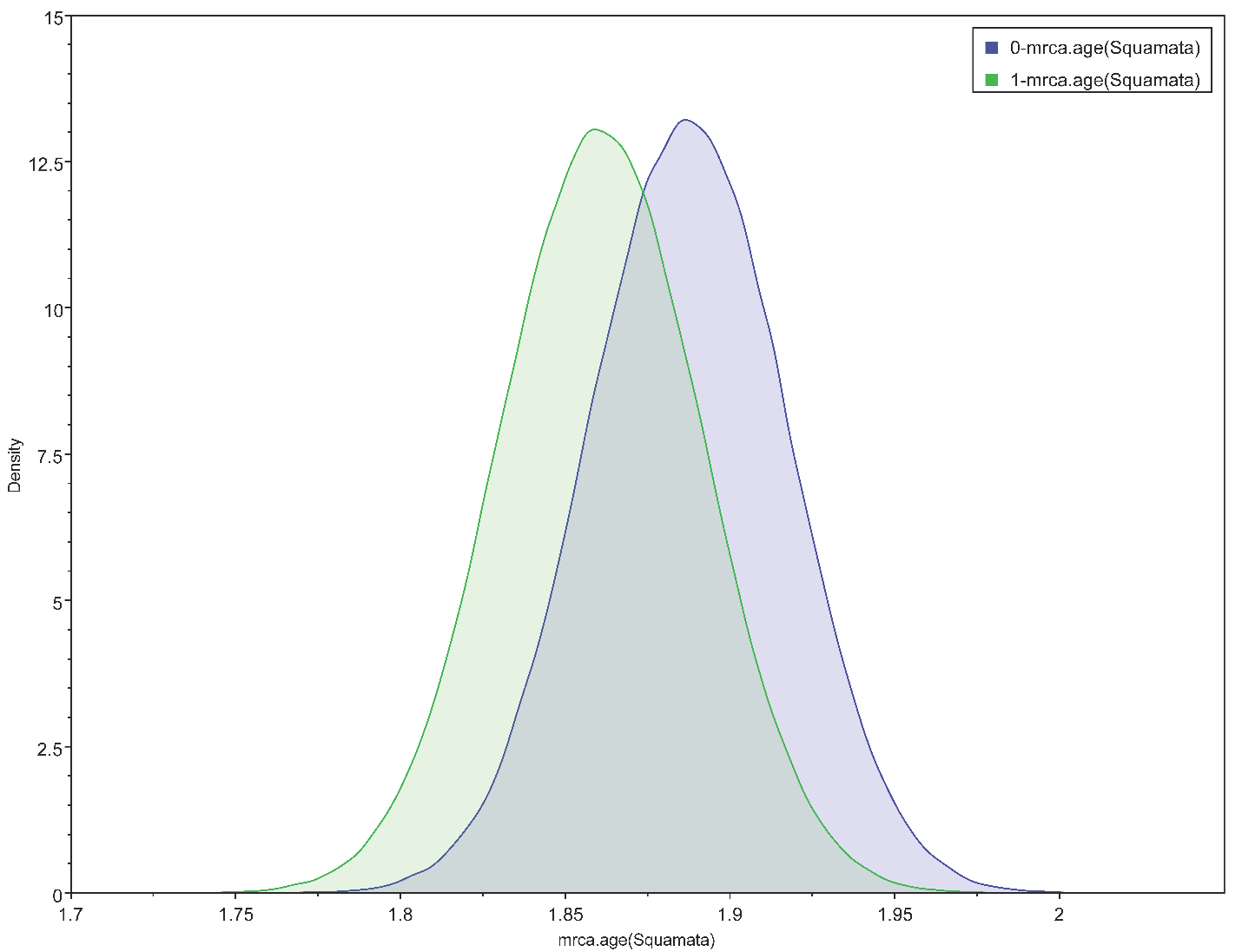

**C**

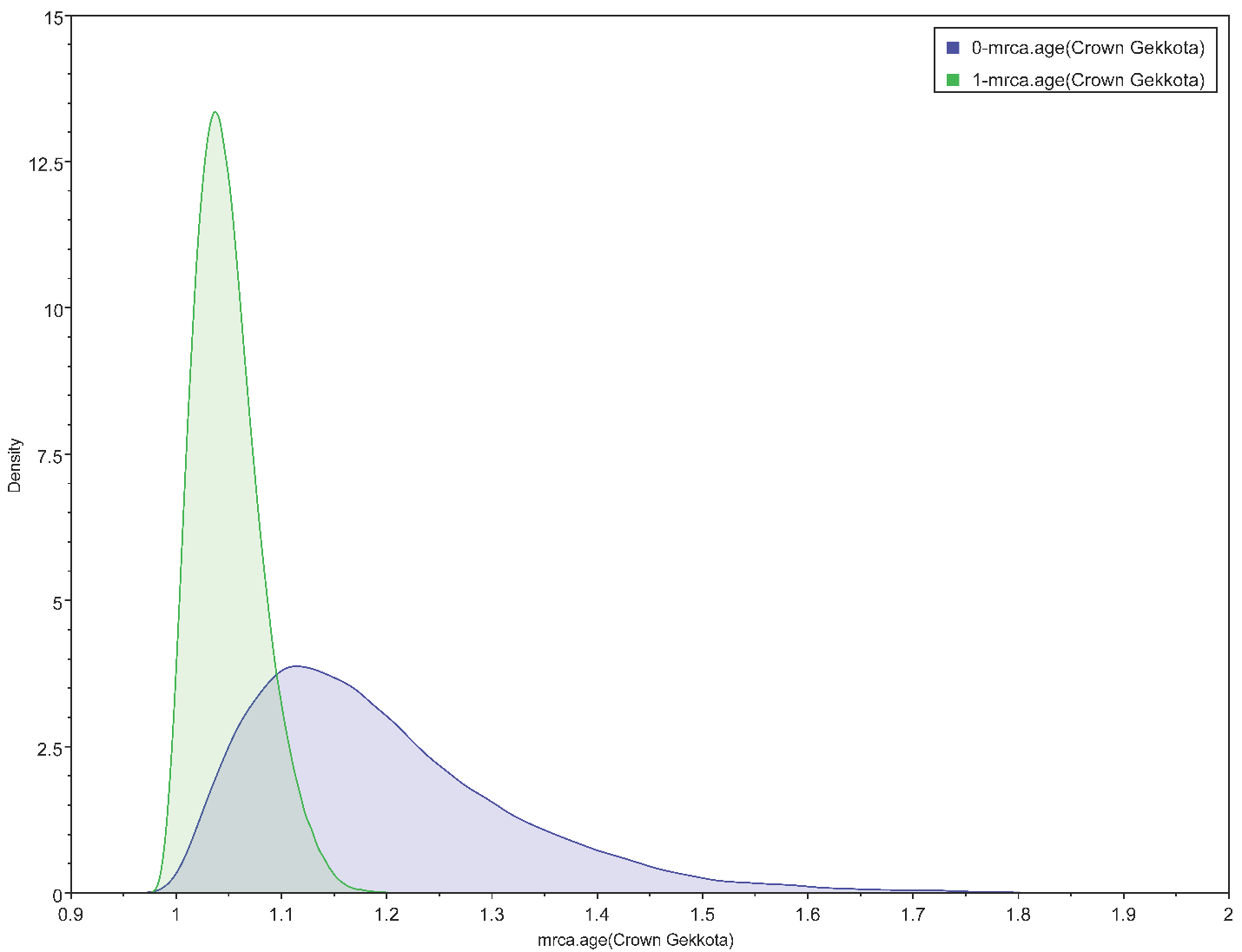

**D**

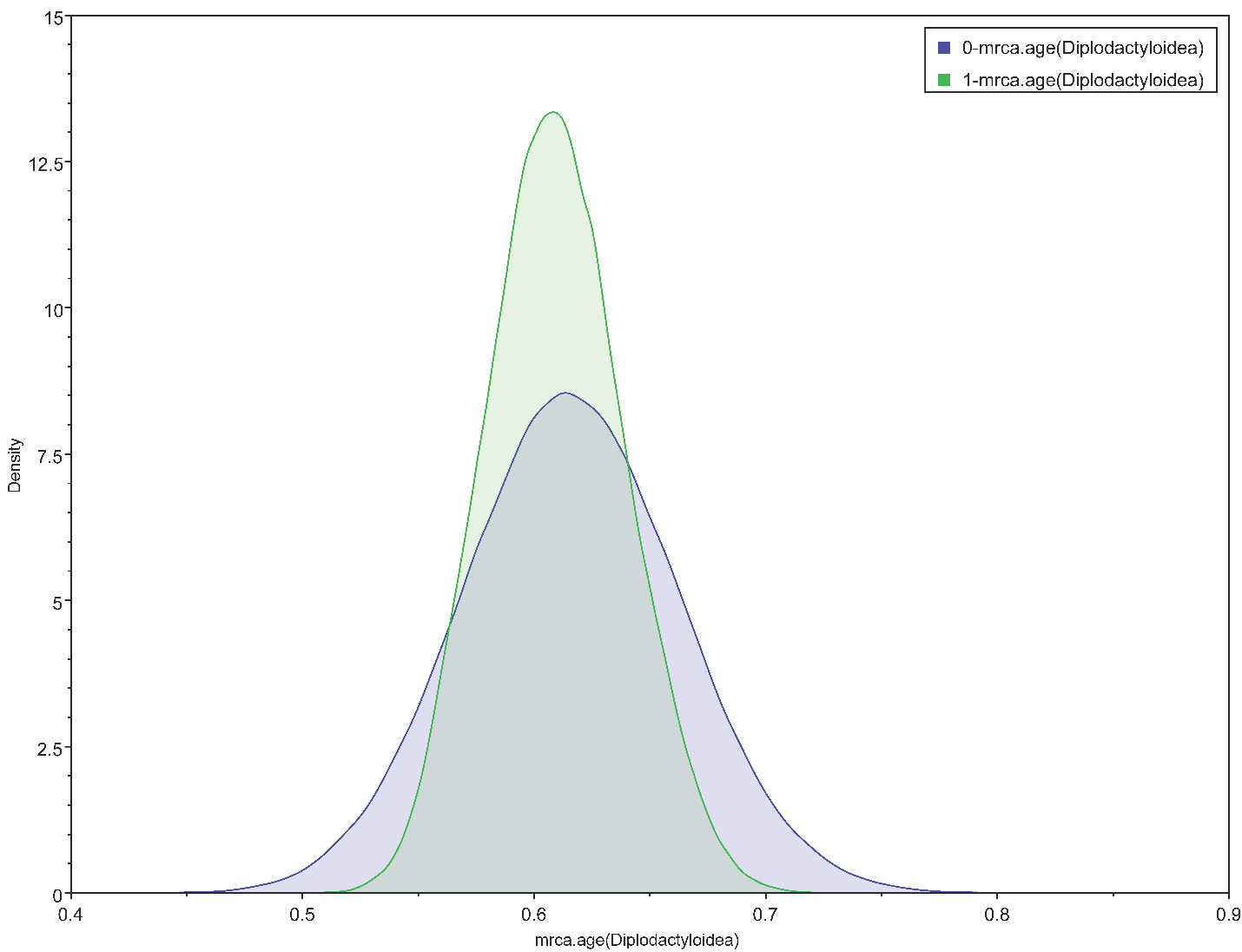

**E**

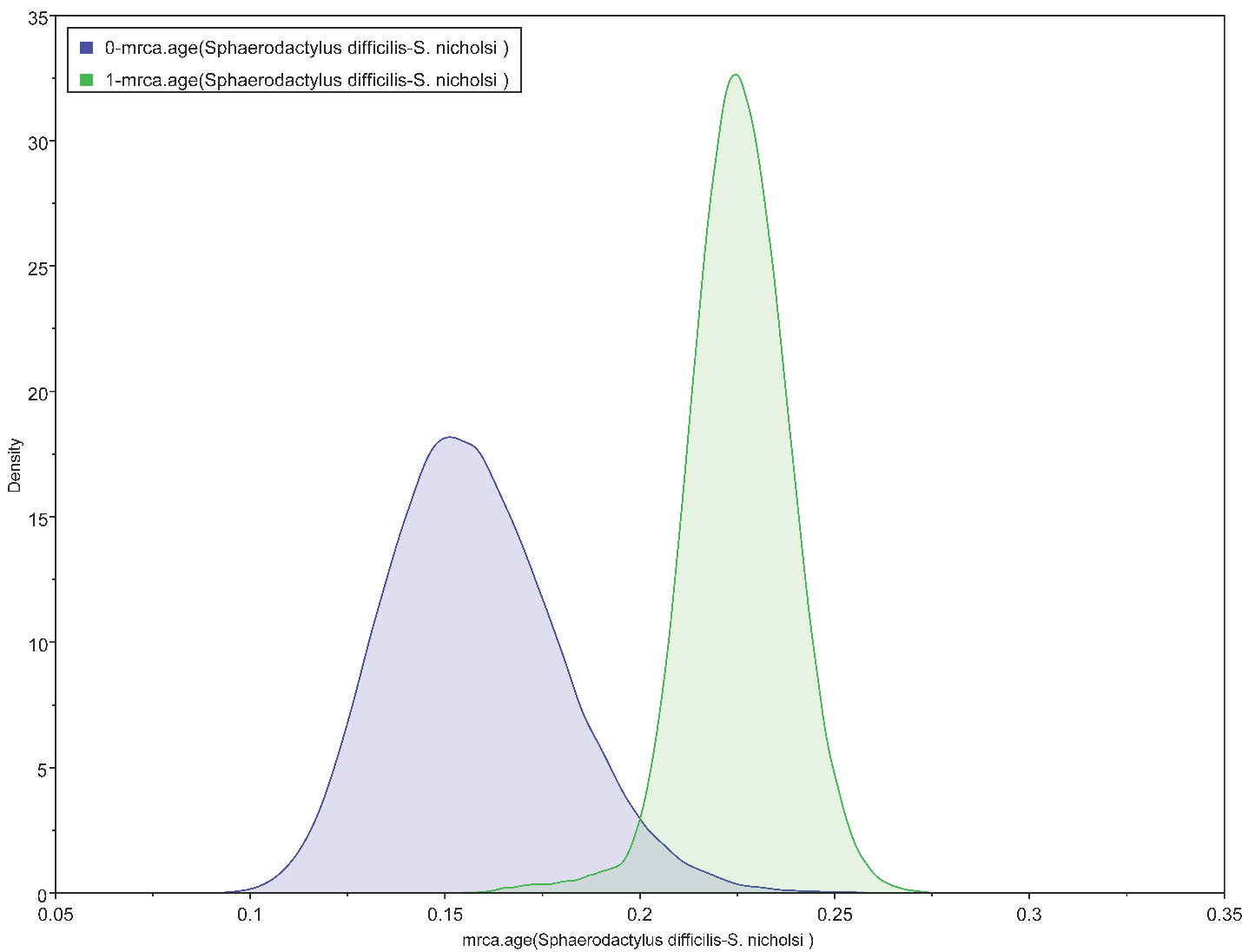

**F**

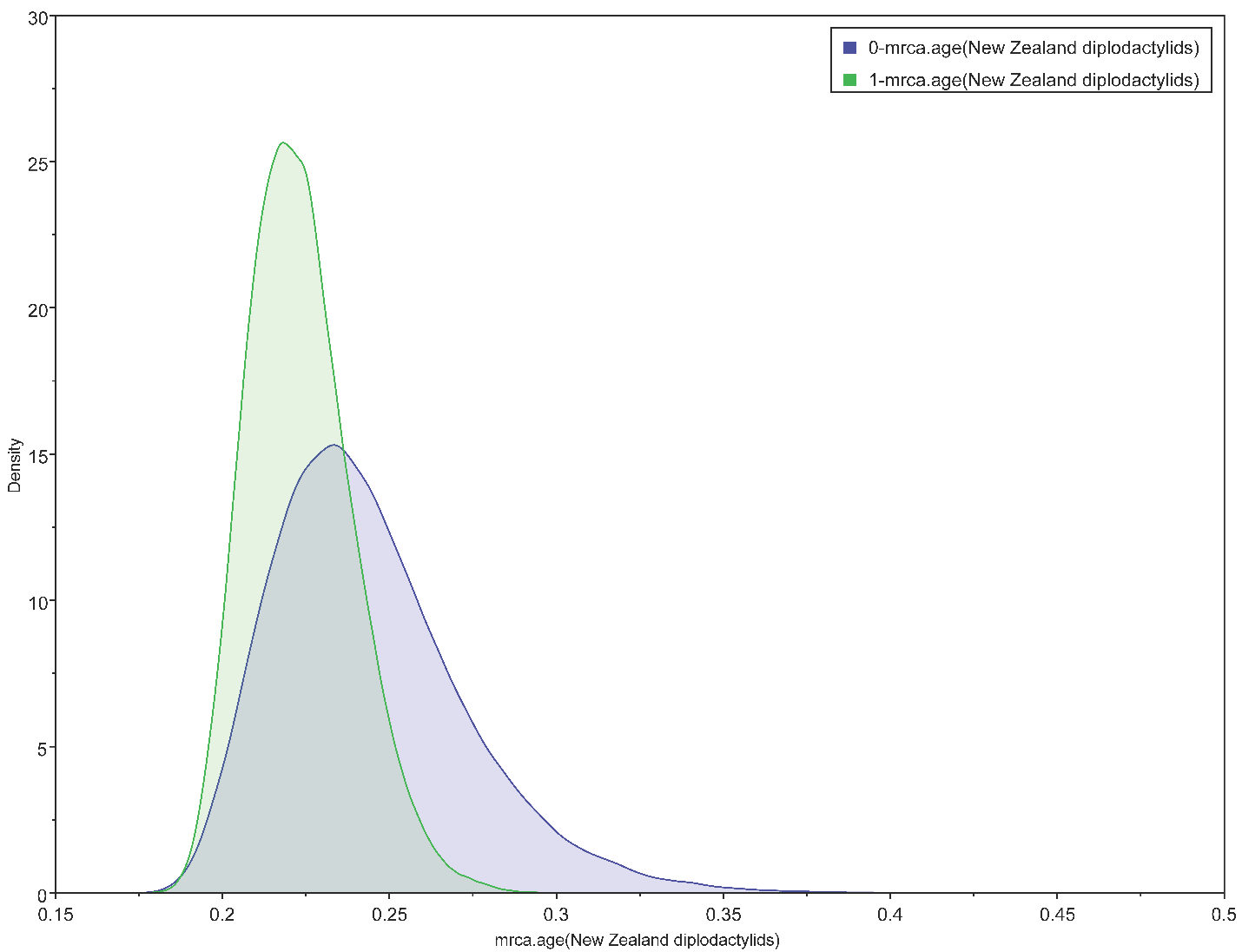

**G**

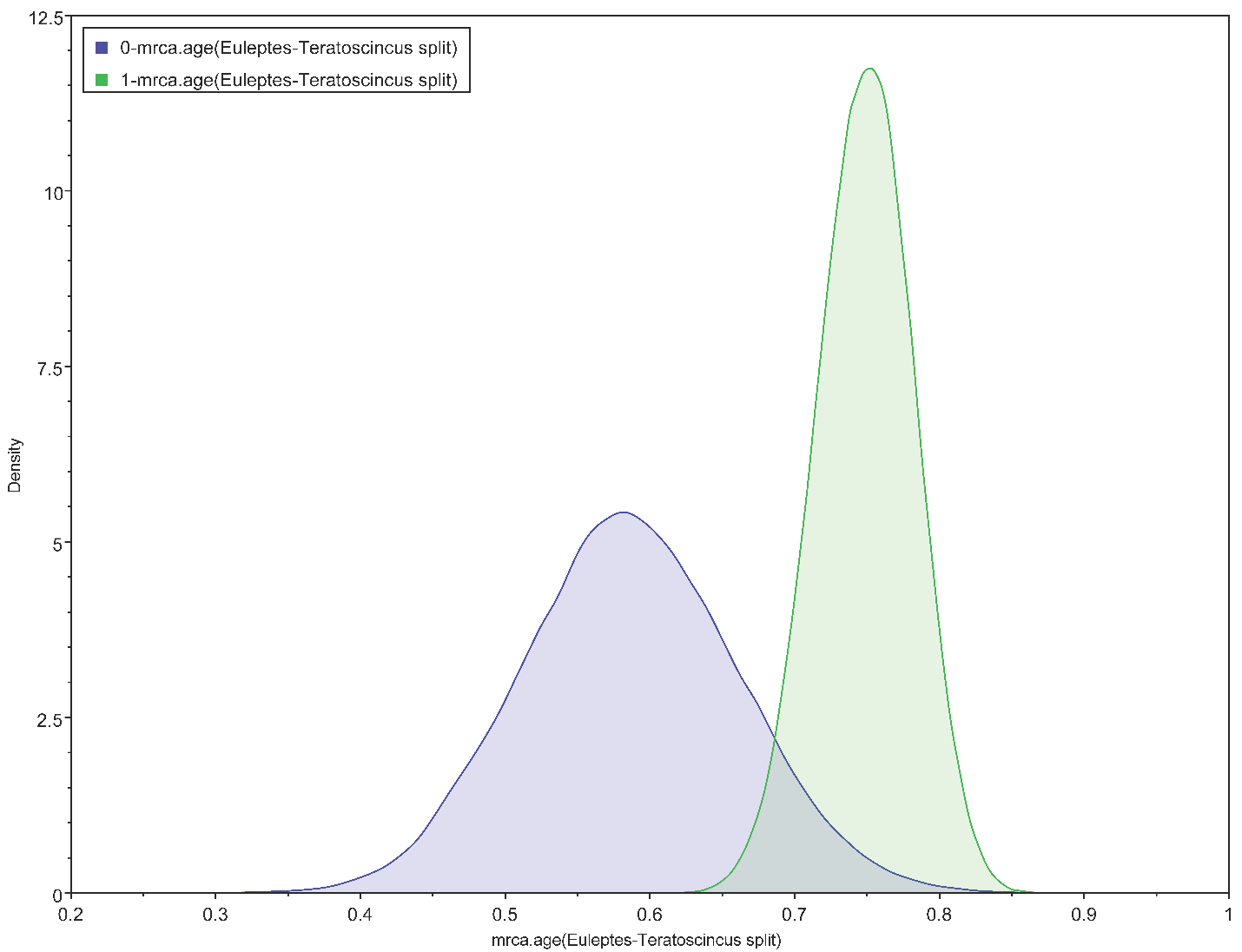

**H**

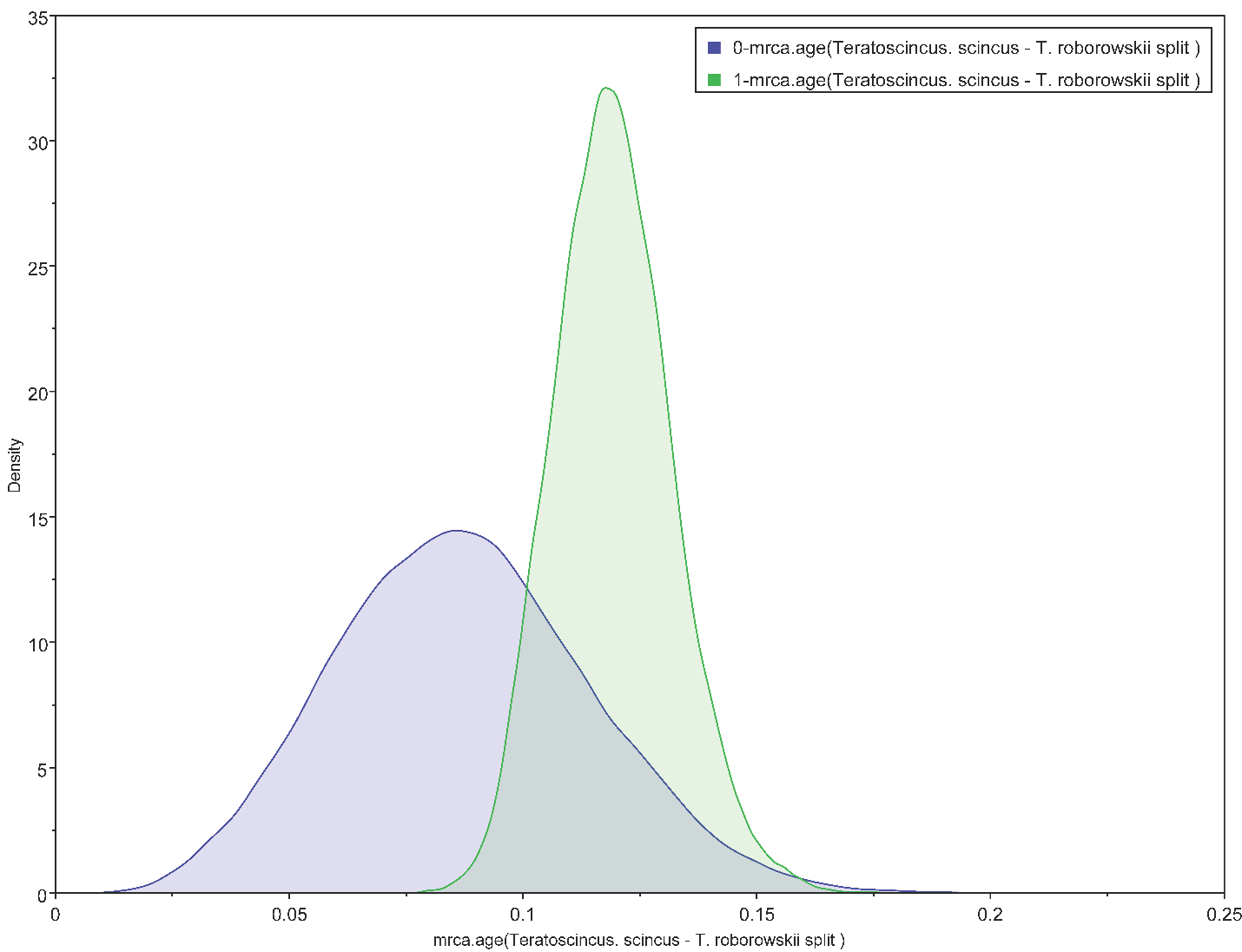

**I**

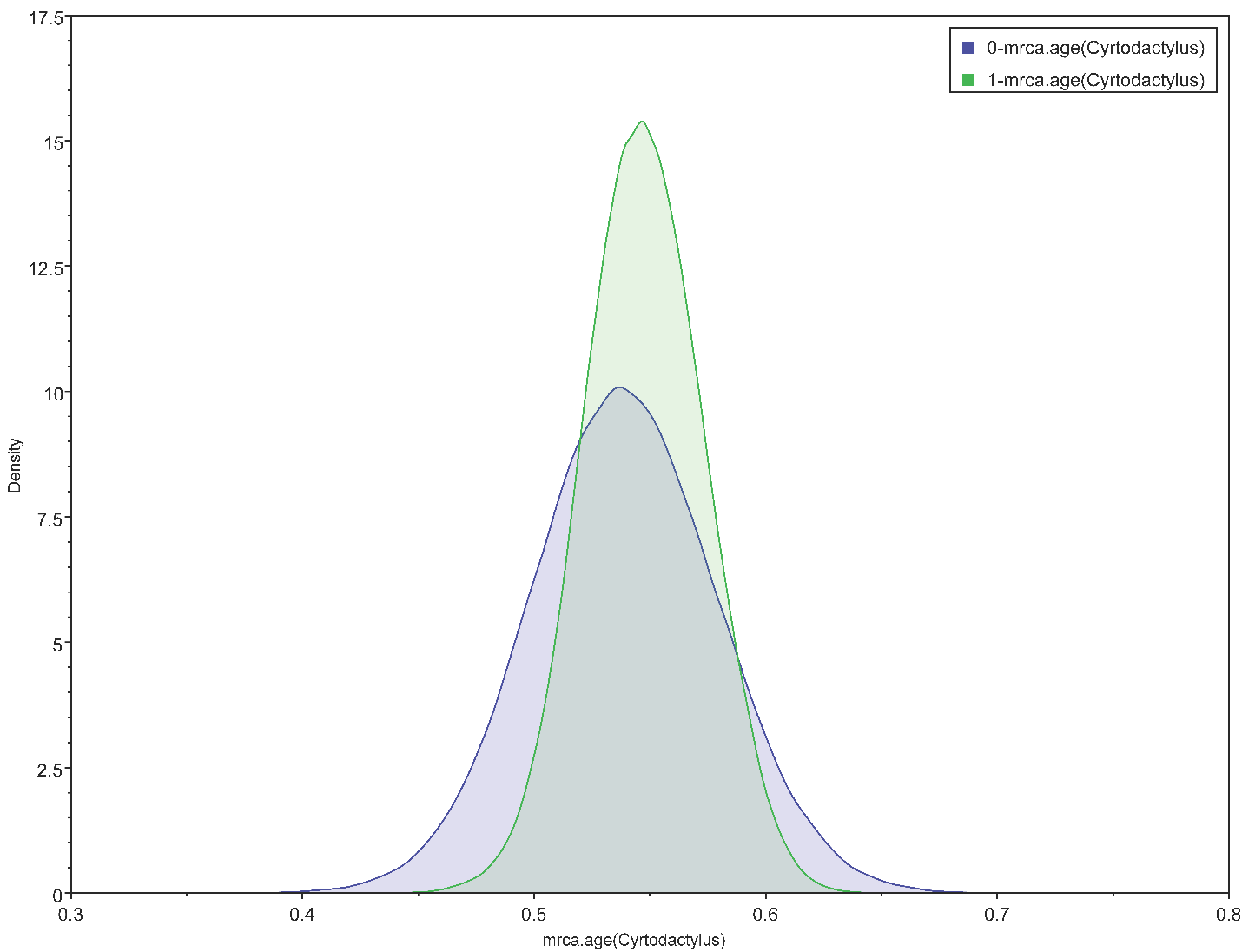

**J**

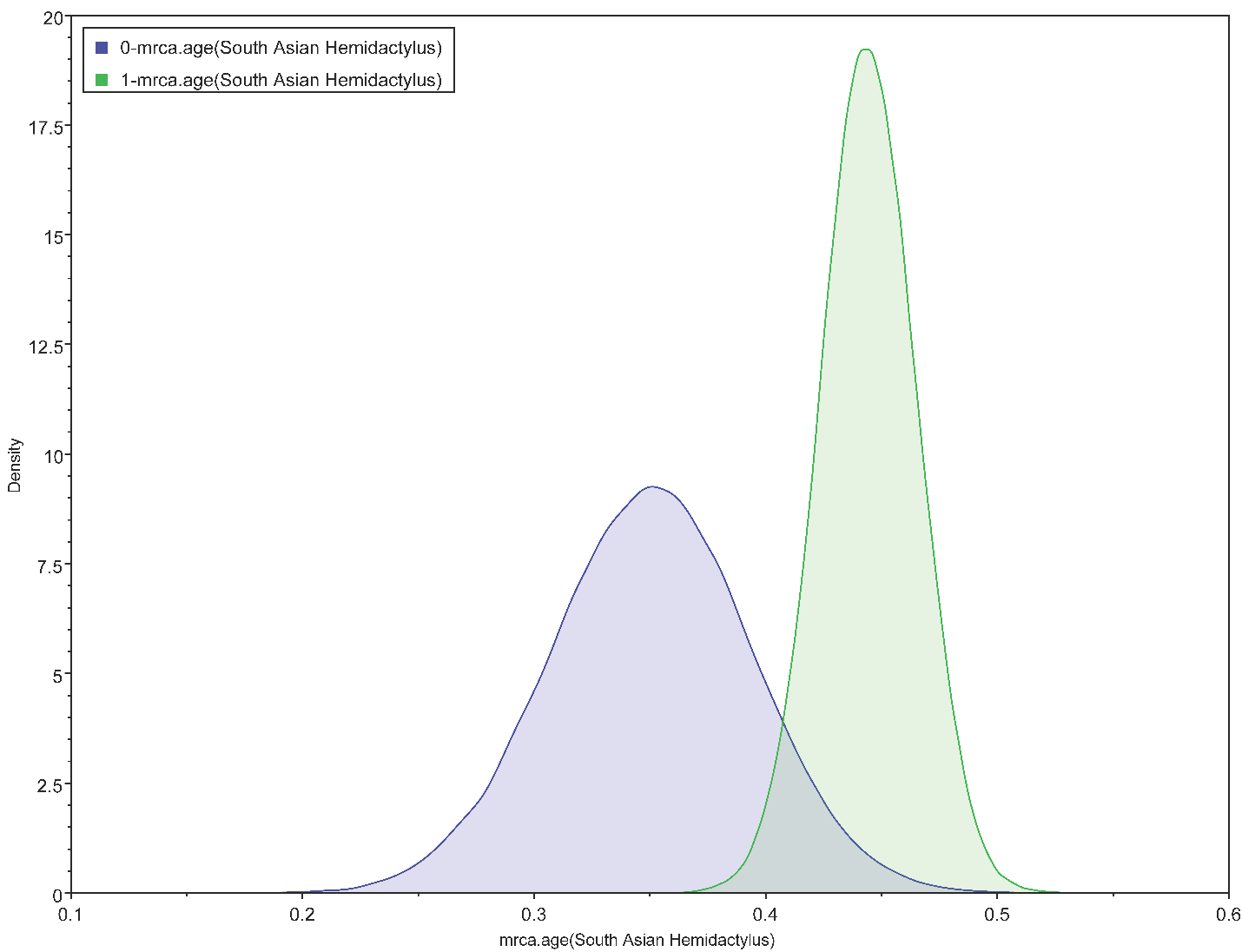

**K**

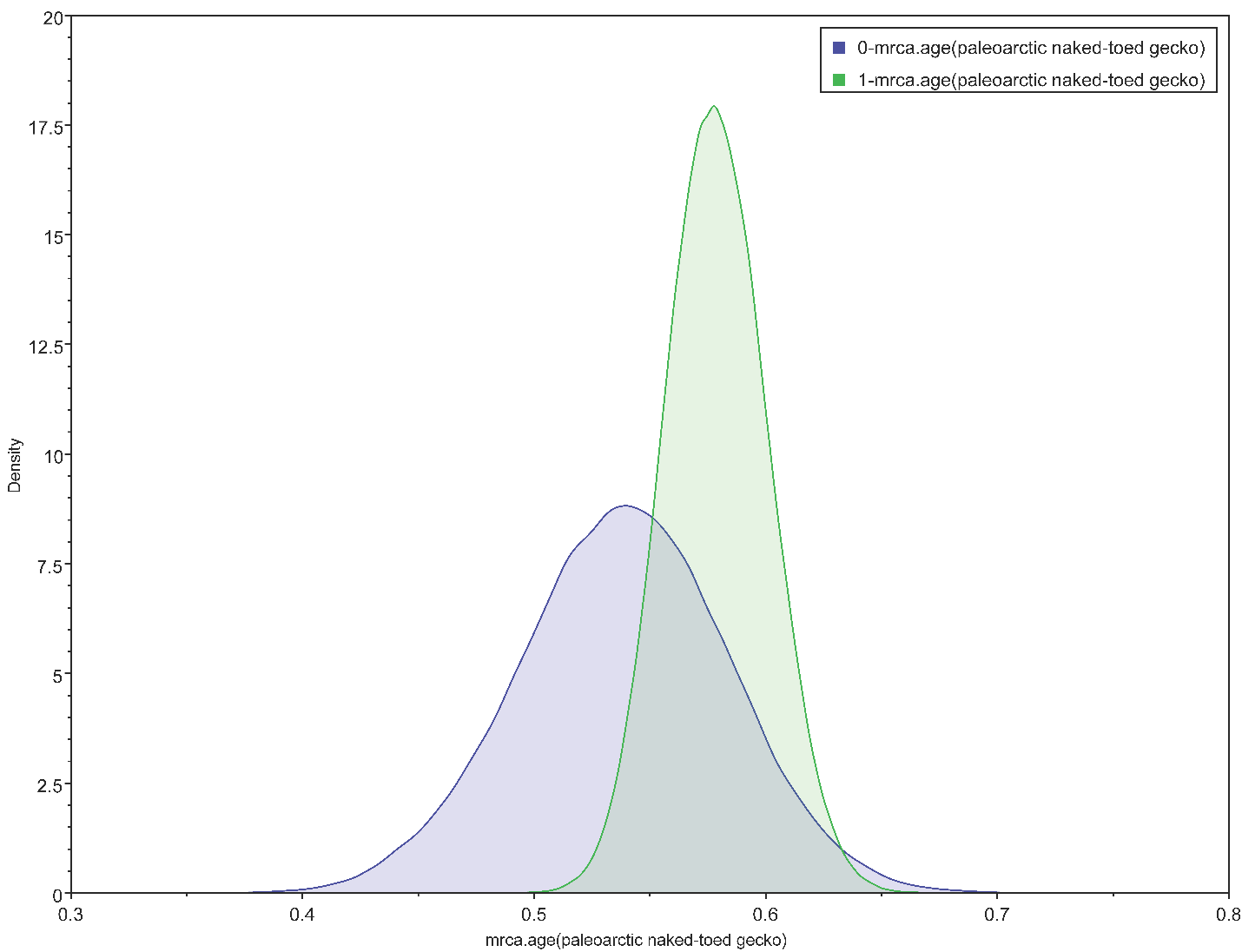

**L**

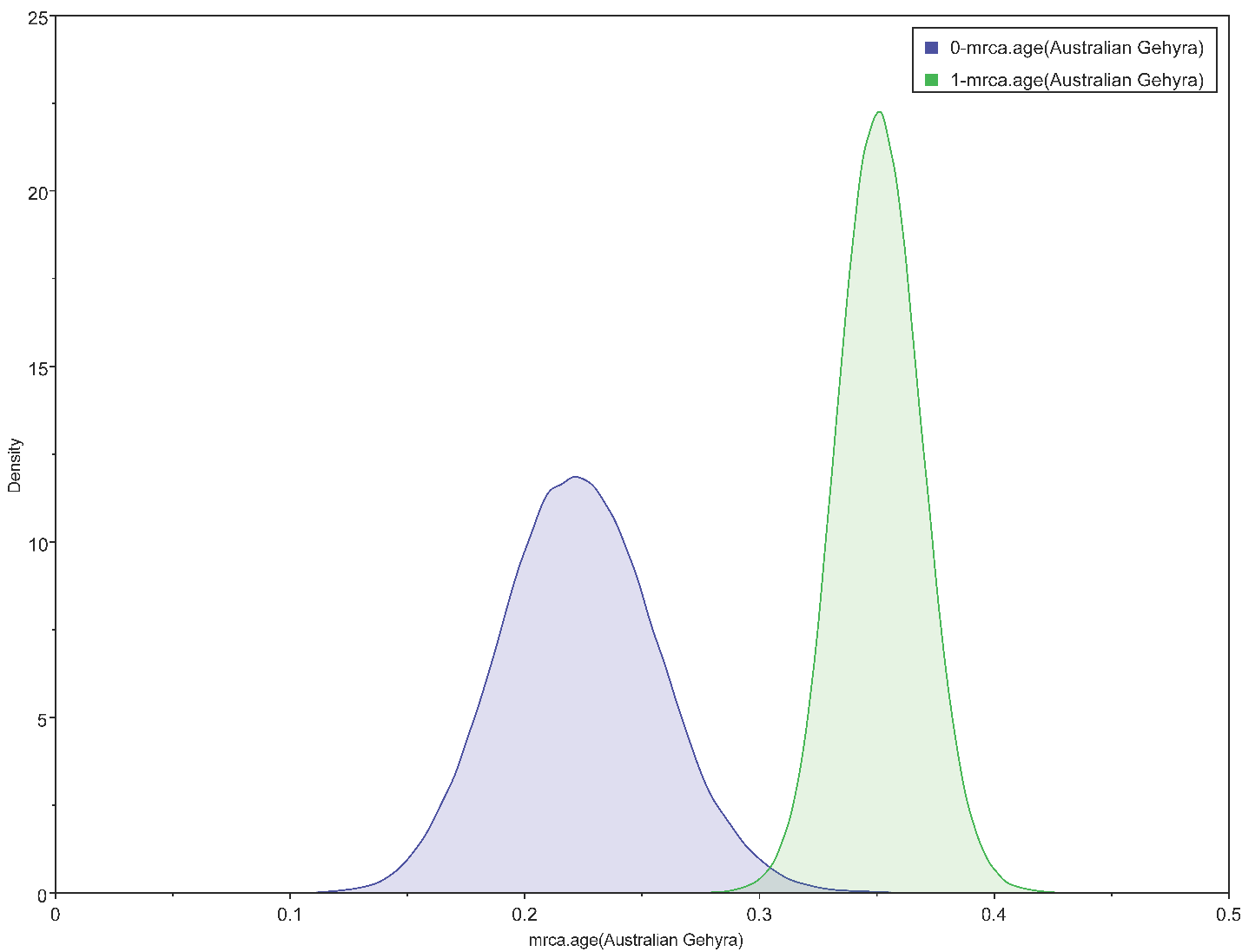

**M**

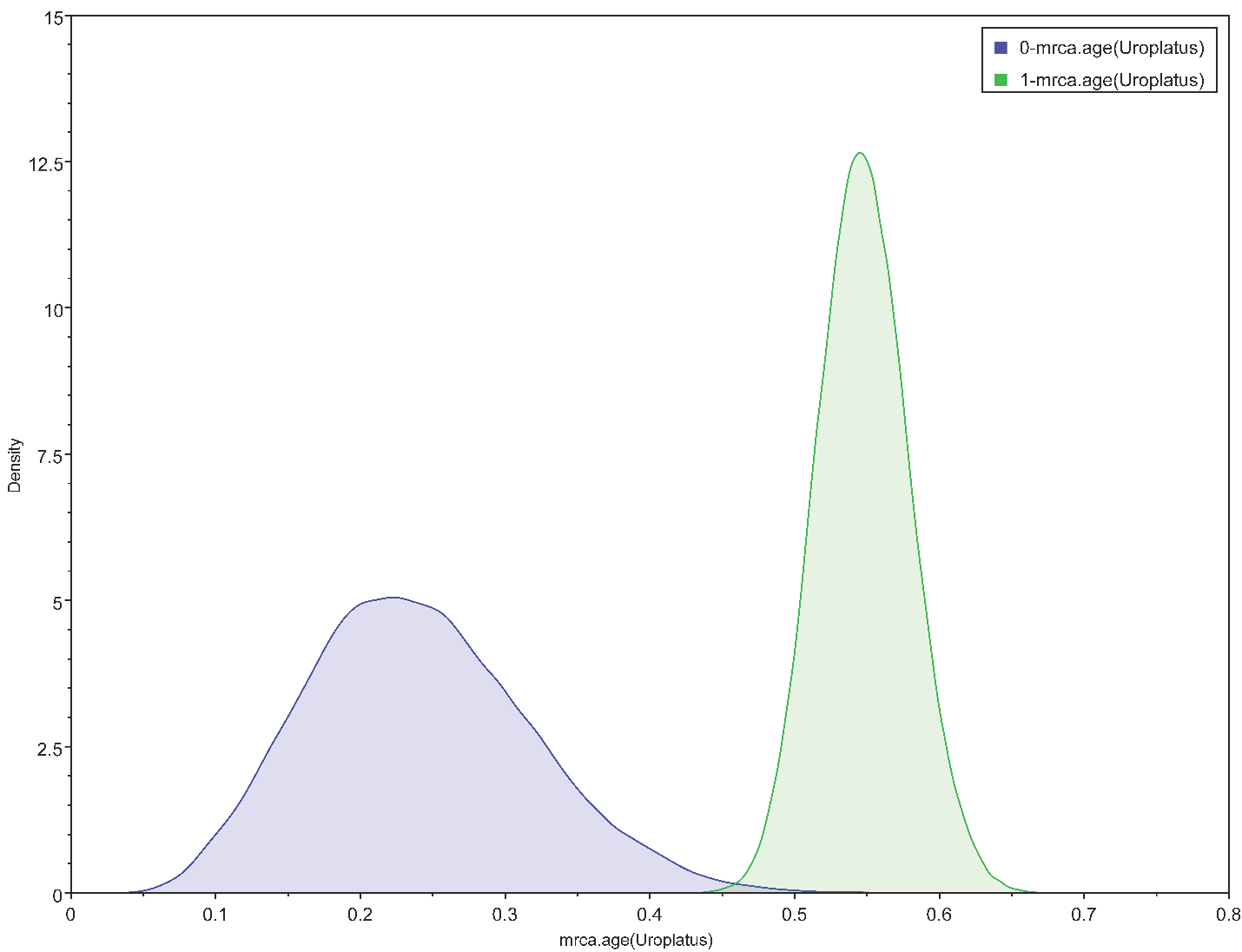

**N**

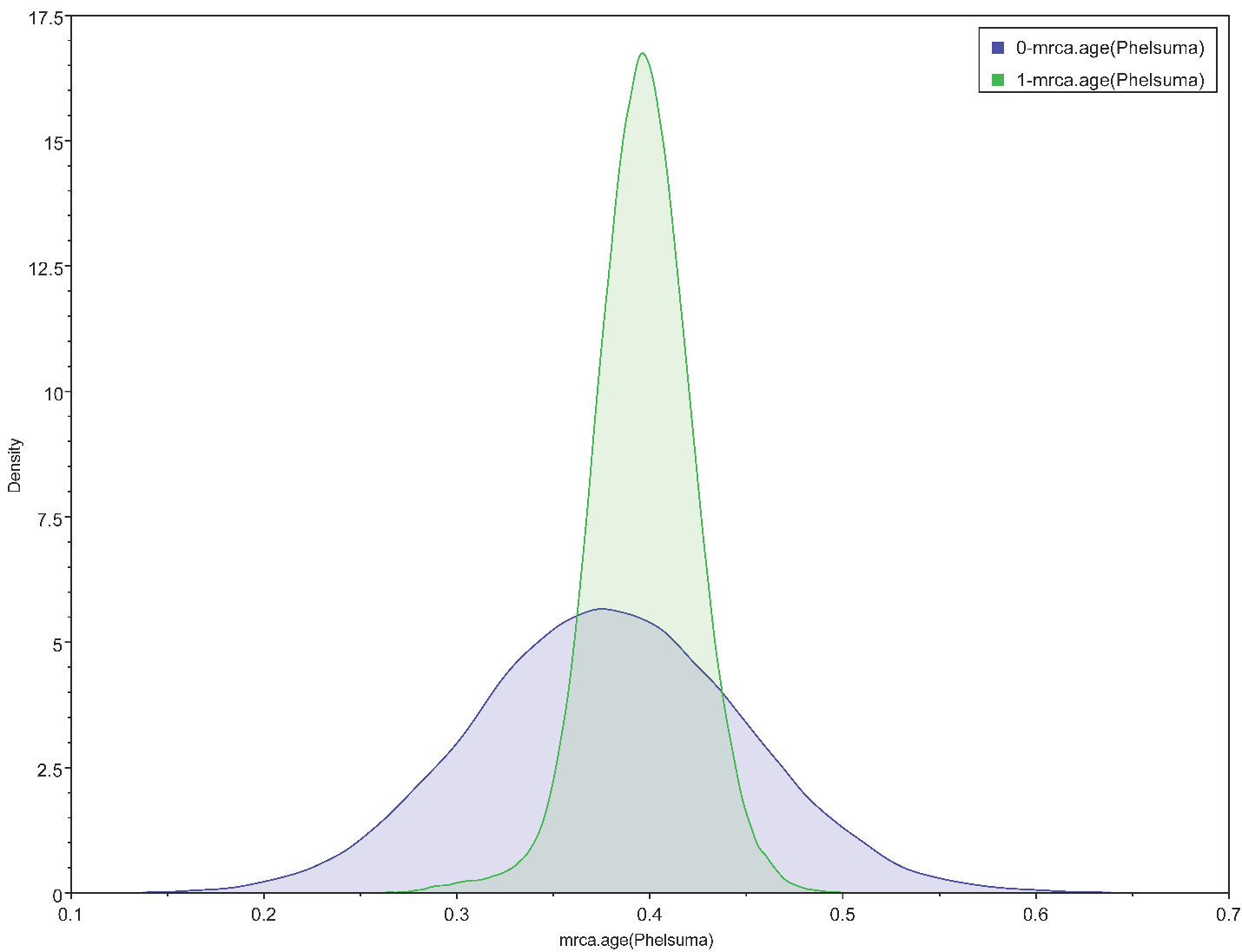

**O**

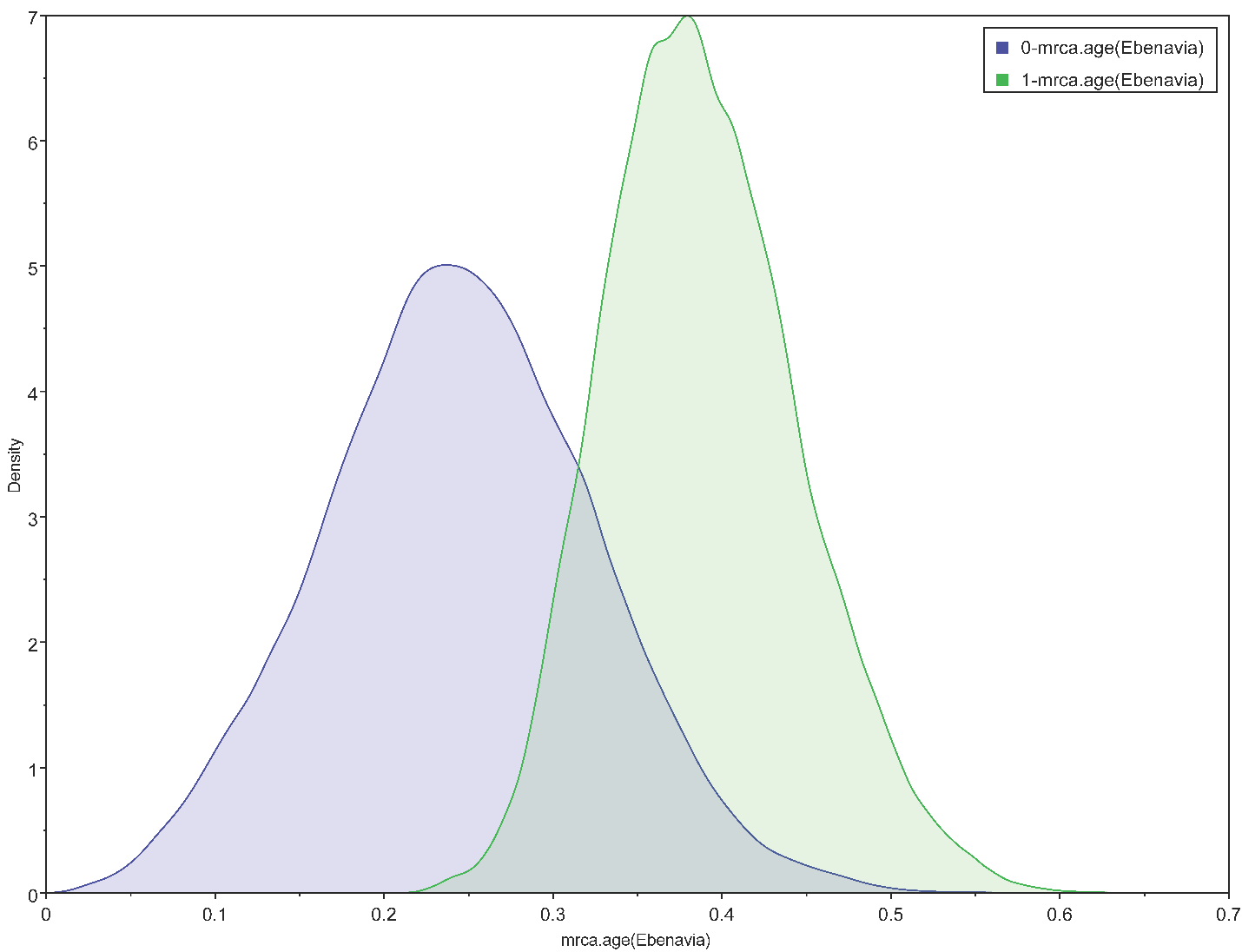

**P**

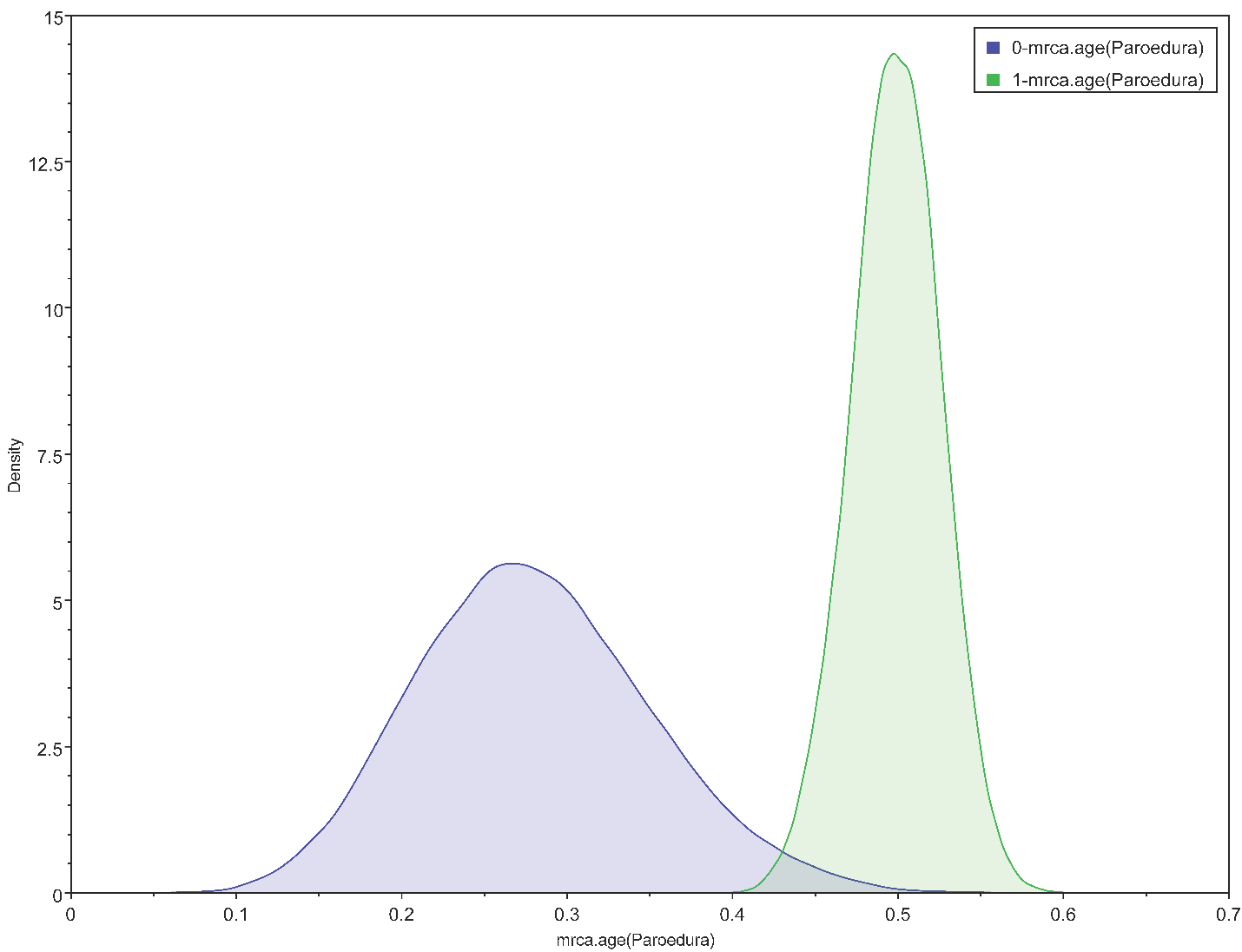

**Q**

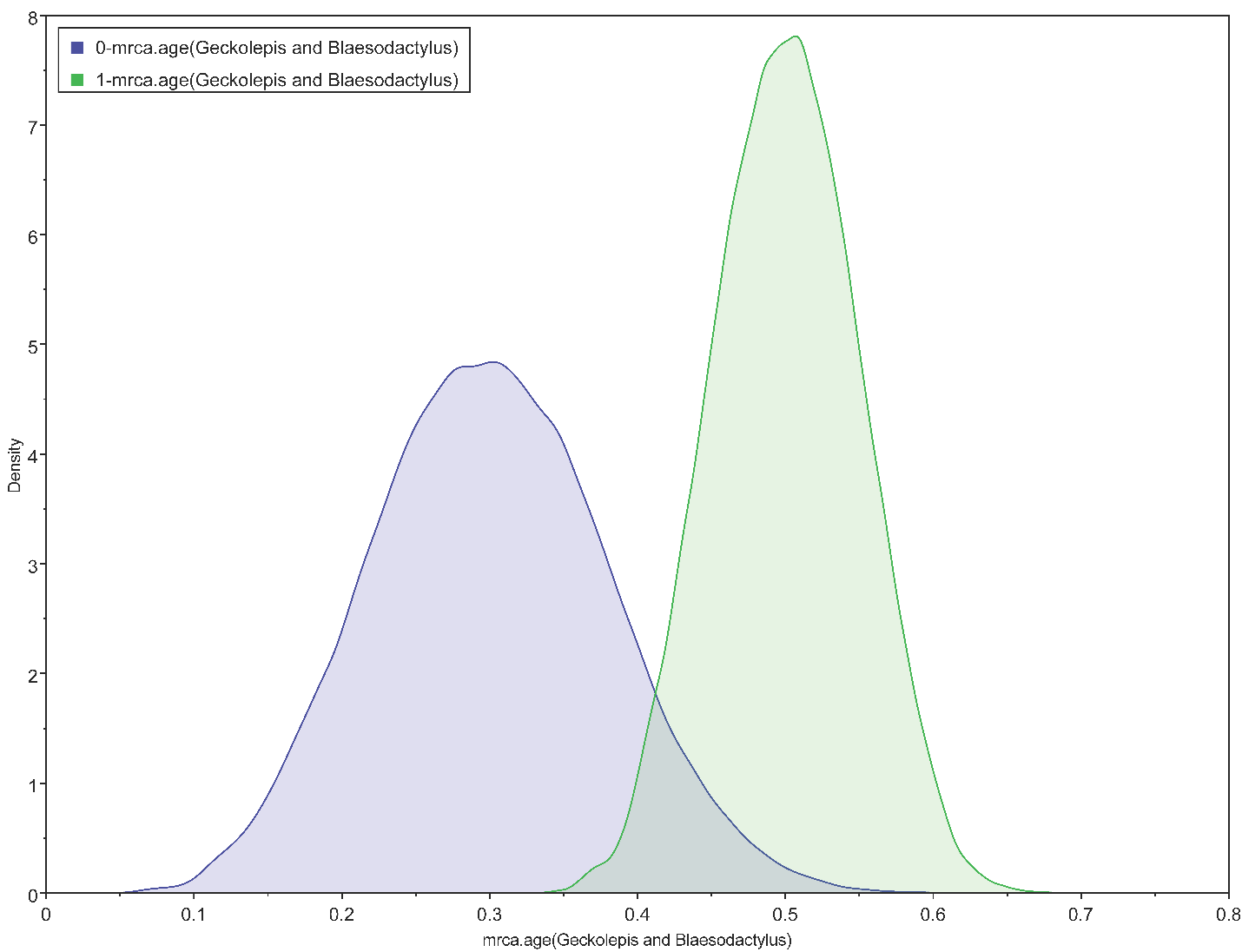

**R**

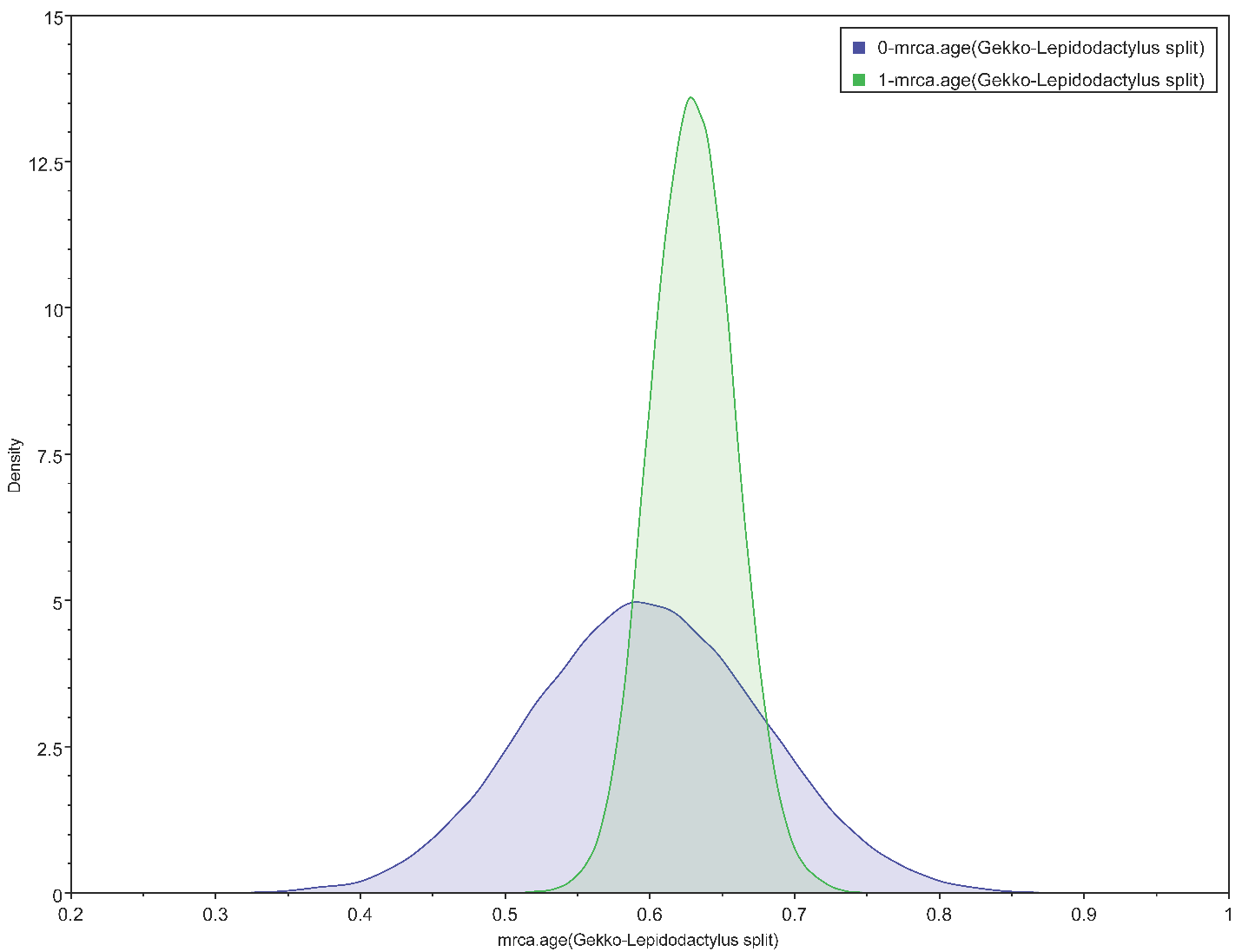

**S**

Figure S4. A: Comparing mean node prior ages to mean node posterior ages of the full + moreprior dataset. Thin black line is the x=y line. Thick cyan line represents the fitted linear model with formula y ~ x using “lm” method within the *geom_smooth()* function of ggplot 2. B-S: Kernel density plot in Tracer for the calibration points of the phylogeny. 0 in legend refers to the prior density while 1 in the legend refers to the posterior density.

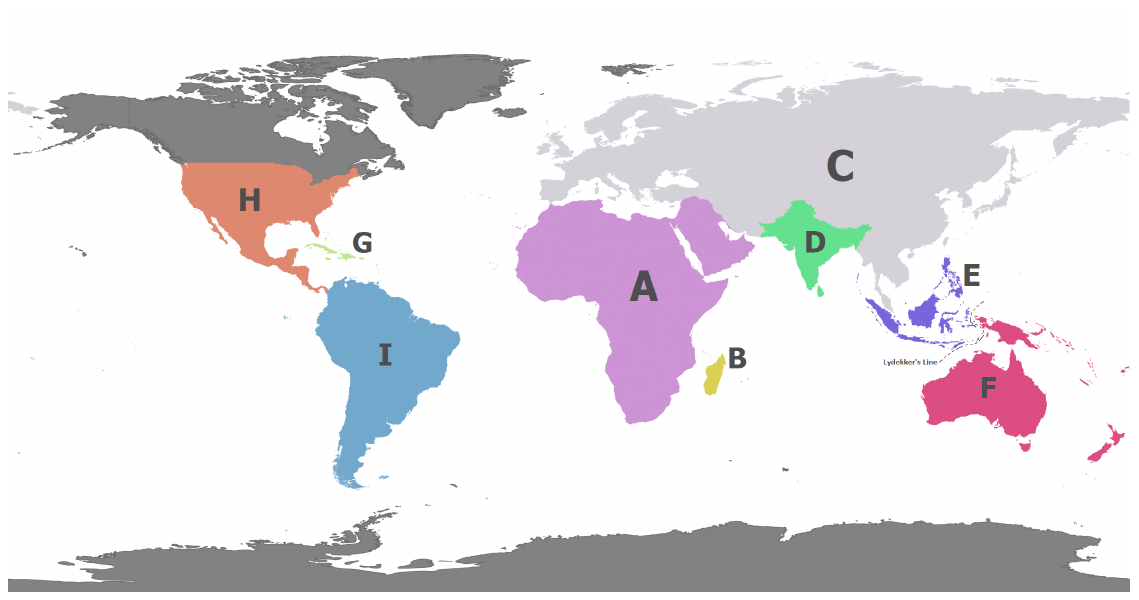

Figure S5. World map colored based on biogeographic regions defined in this study. The color is consistent with colors used in plotting. This map is produced using ArcGIS Pro. State coding: A-Africa-Arabia; B-Madagascar; C-Eurasia; D-South Asia; E-Sundaland; F-Australasia; G-Carribean; H-North-Central America; I-South America.

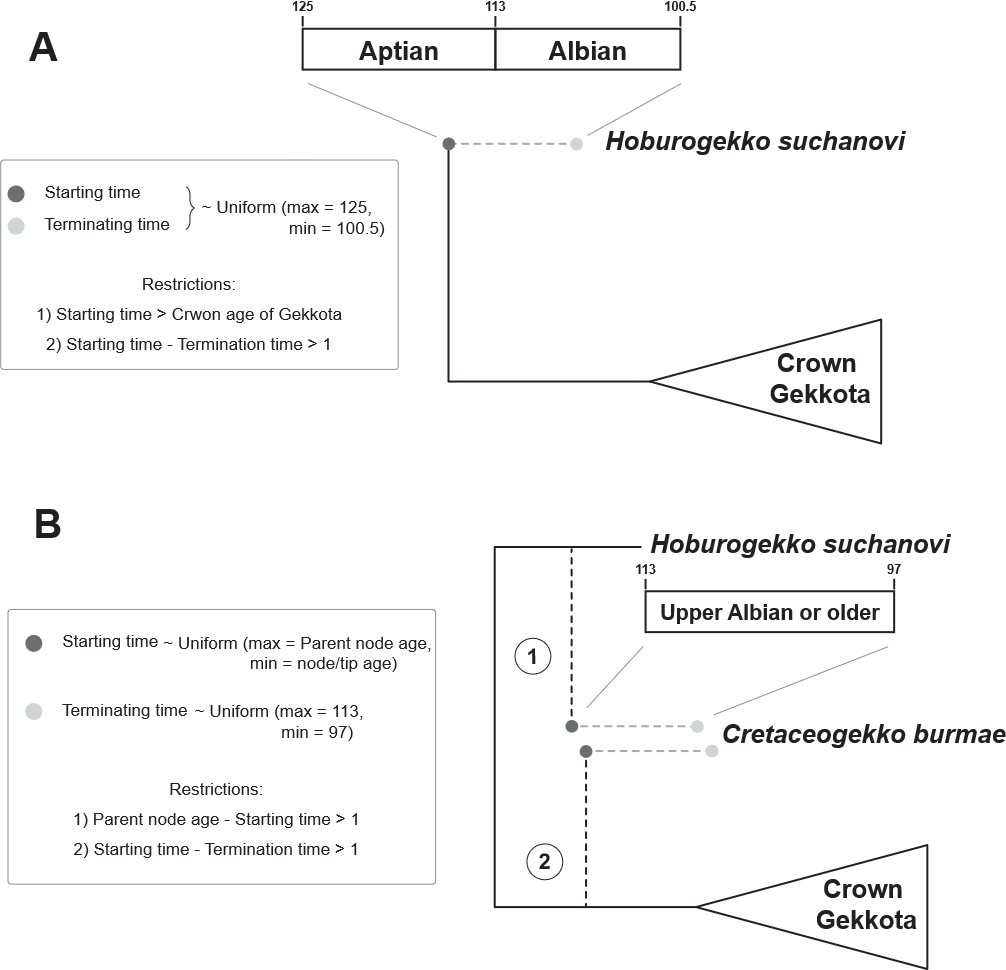

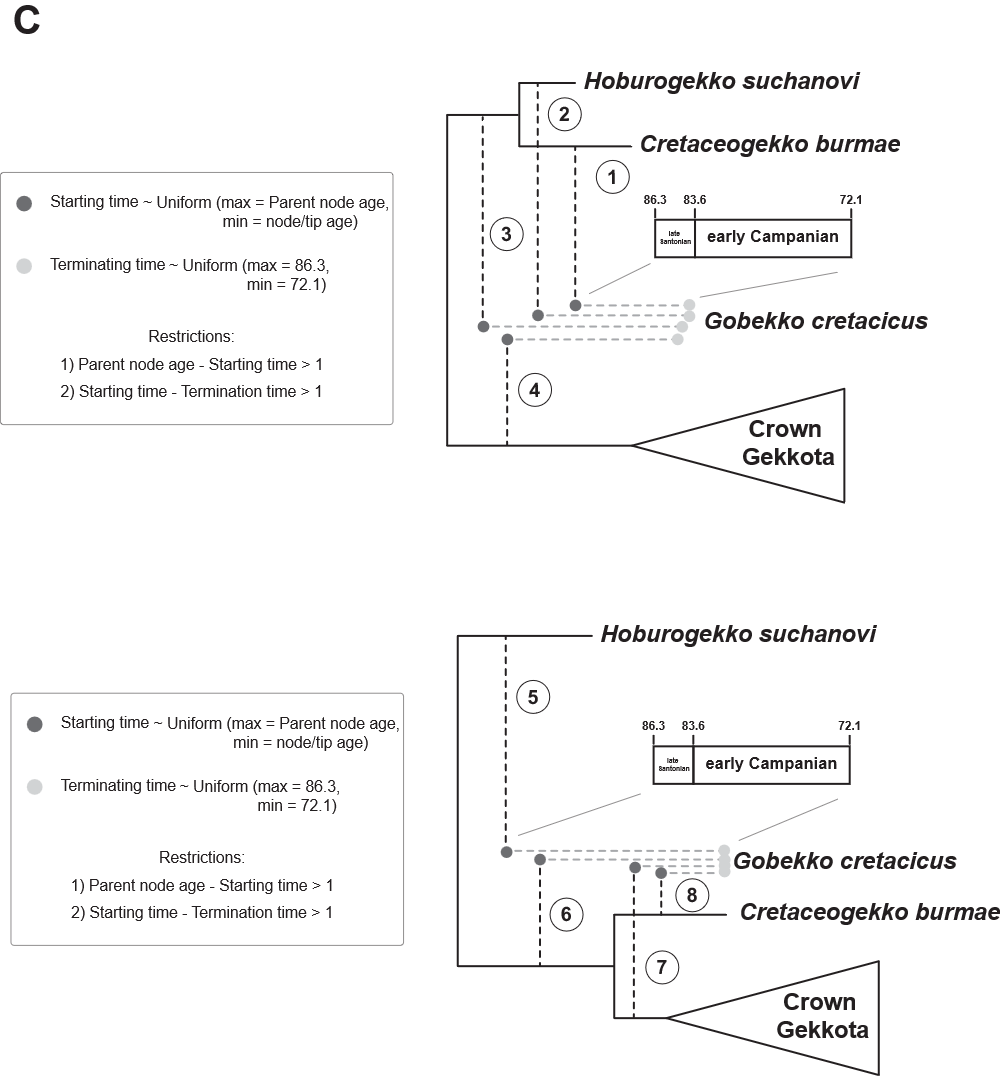

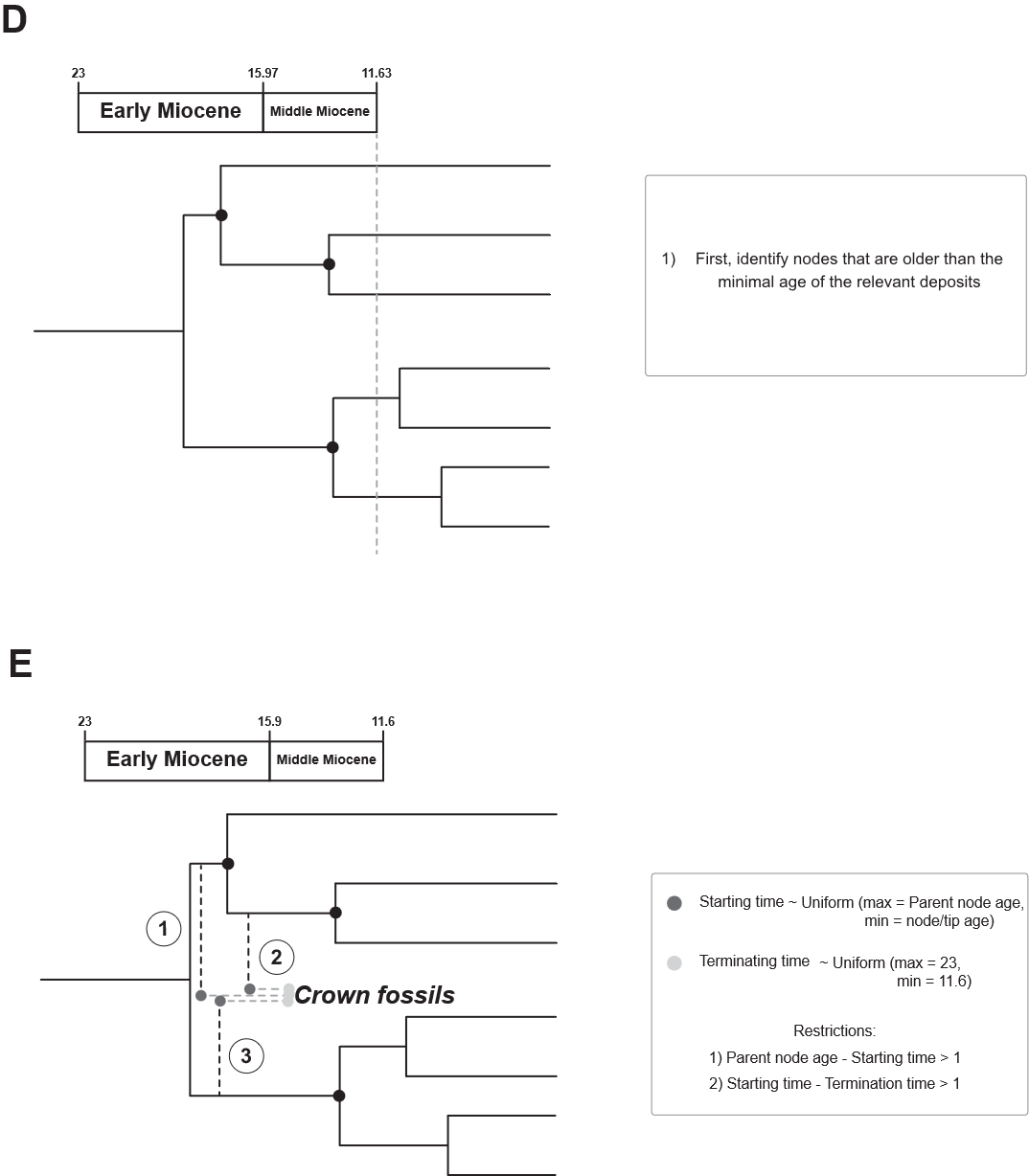

Figure S6. Procedures for attaching fossils to the phylogeny. A-C: Procedures for attaching stem fossils. D-E: Procedures for attaching crown fossils.

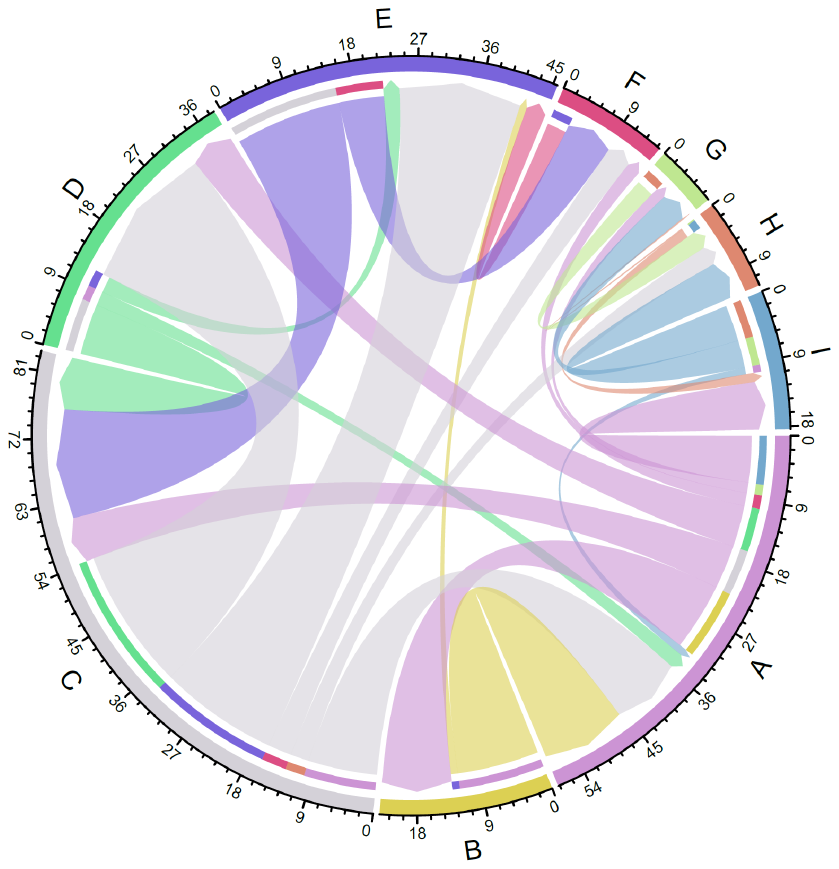

**A**

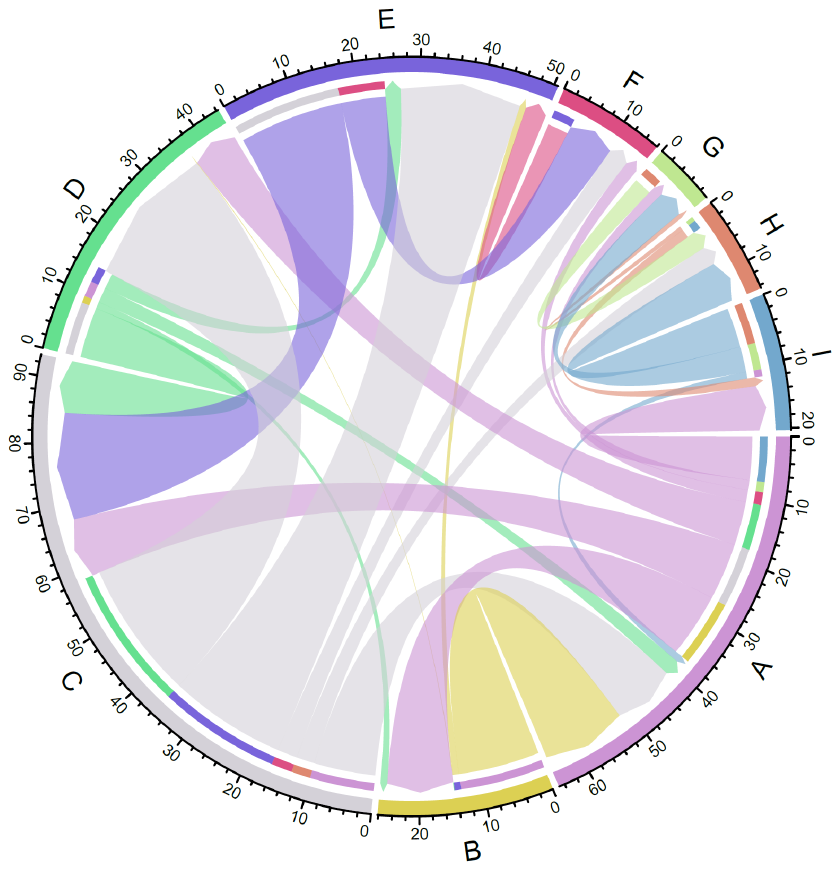

**B**

Figure S11. Chord diagram showing average number of transitions across 100 Biogeographic Stochastic Mapping (BSM) maps across 10 runs, each run with averaged model parameters. These transitions include both anagenetic transitions (range expansion (d)) as well as cladogenetic transition (founder-speciation (j)). A: Analyses with fossils randomly attached to the phylogeny. B: Analyses without fossils. The inner ring indicates the range it disperses to, and the thick arrow shows the range it disperses from. State coding: A-Africa-Arabia; B-Madagascar; C-Eurasia; D-South Asia; E-Sundaland; F-Australasia; G-Carribean; H-North-Central America; I-South America.

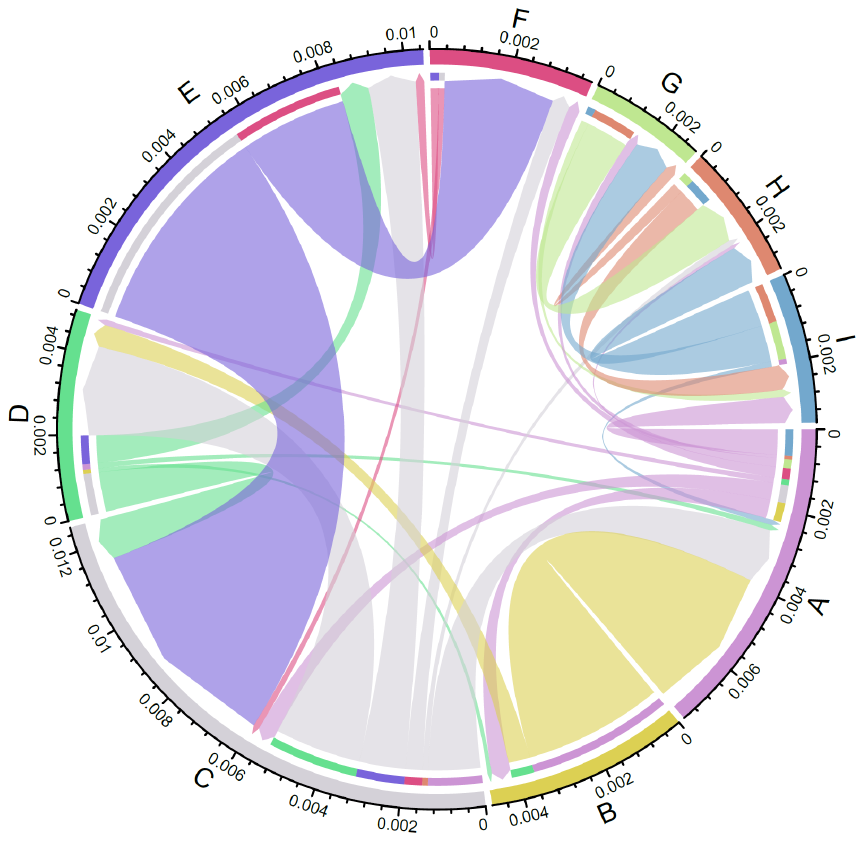

Figure S12. Chord diagram showing posterior estimated transition rate between states in MuHisse model. The inner ring indicates the range it disperses to, and the thick arrow shows the range it disperses from. State coding: A-Africa-Arabia; B-Madagascar; C-Eurasia; D-South Asia; E-Sundaland; F-Australasia; G-Carribean; H-North-Central America; I-South America.

**A**

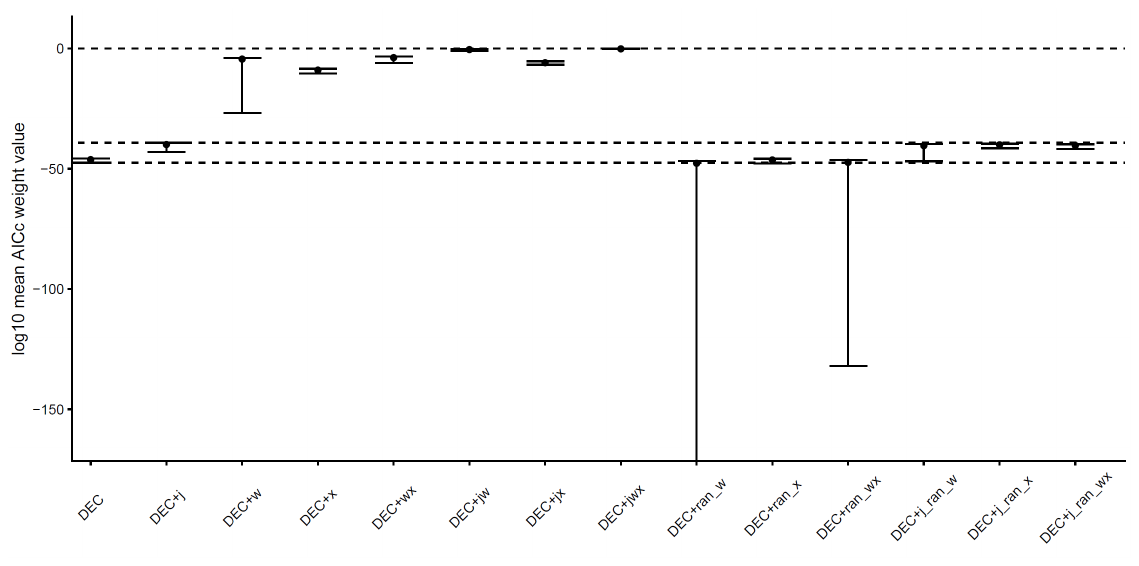

**B**

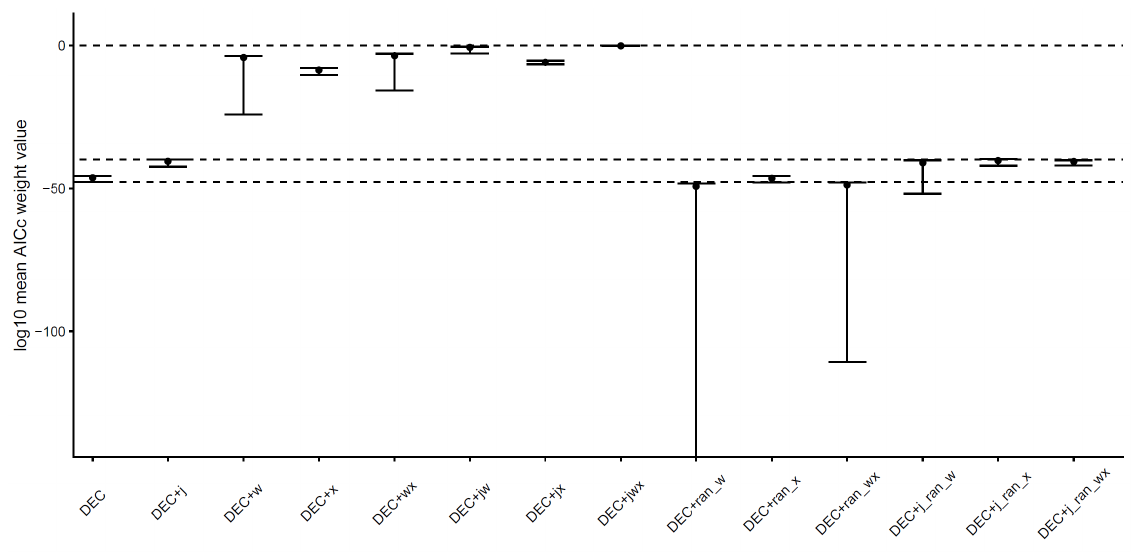

Figure S13. AICc weight value for each model across all runs. A: Analyses with fossils randomly attached to the phylogeny. B: Analyses without fossils. The point is the log10 of the mean AICc weight value, and the error bar represents the maximum and the minimum values. Higher value means higher AICc weight value, which means better supported in model comparison. The names of the model that are included in the model comparison are shown in X axis. The name with “ran” means the model includes the randomly generated distance (+x) or manual dispersal multiplier (+w) matrices, or both. The dashed lines from top to bottom indicates number zero, maximum value of the DEC+j AICc wight value, and minimum value of the DEC AICc weight value respectively.

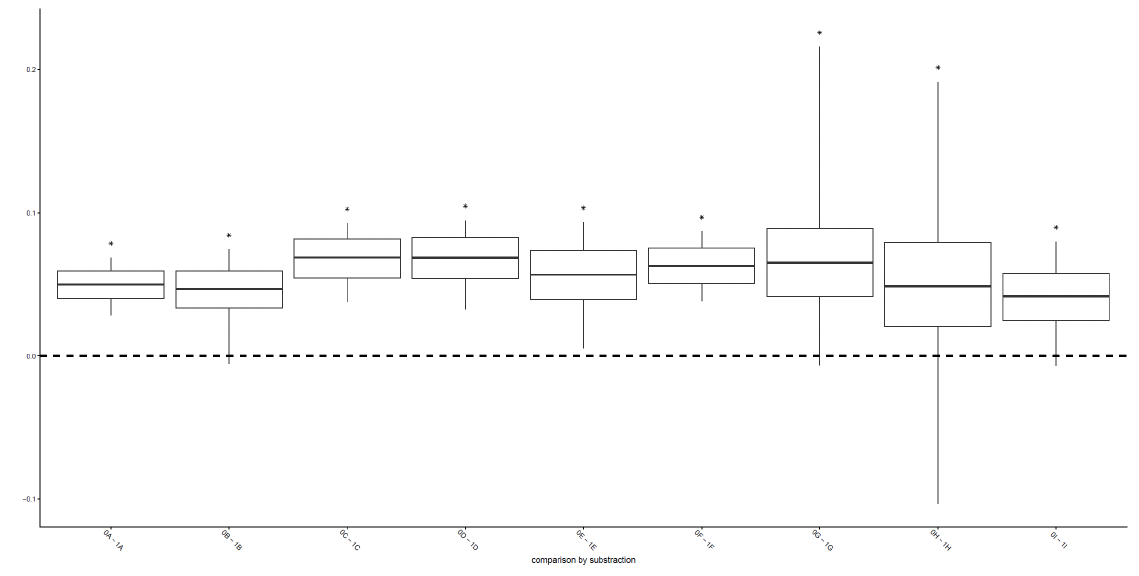

**A**

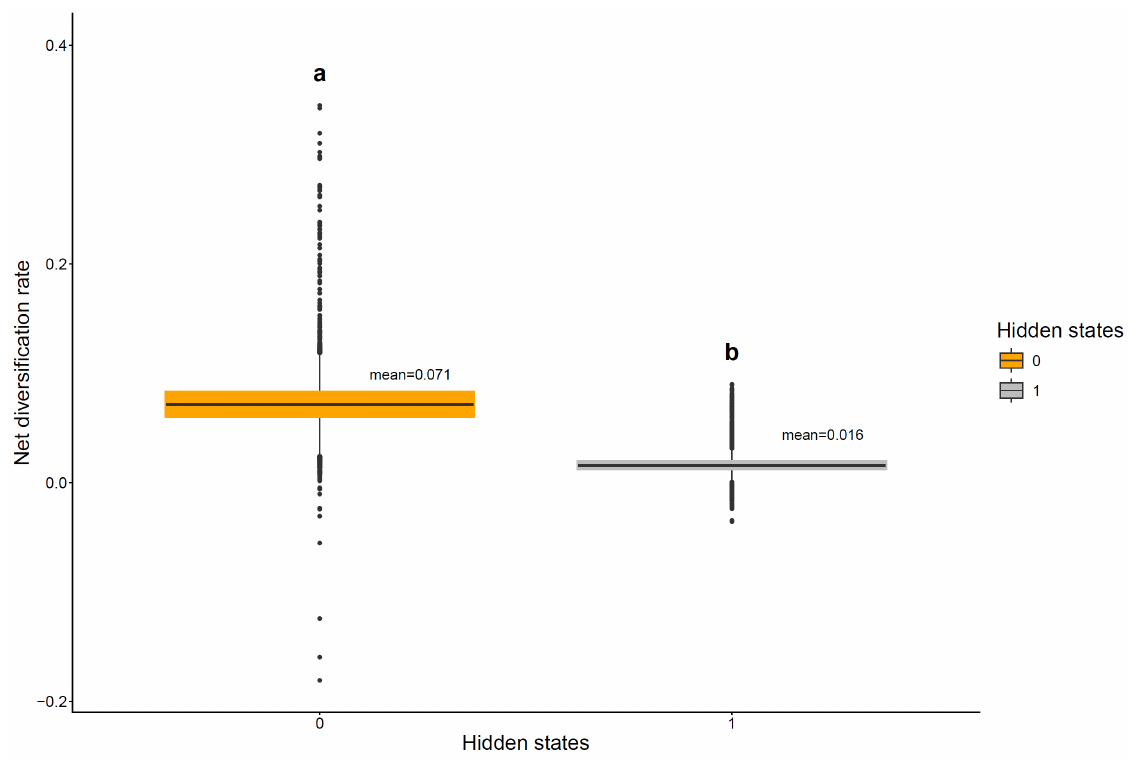

**B**

Figure S14. Boxplot showing posterior estimates of the net diversification rate (speciation rate – extinction rate) for hidden states 0 and 1. A: Boxplot showing stats of subtracting the rates of the same observed states with different hidden states in each iteration, e.g. 0A – 1A, 0B - 1B, etc.. In this case, the lower and upper boundaries of the boxplot box are 2.5% and 97.5% quantile respectively. Number 0 is highlighted with a thick dash line. Asterisks above the box mean number 0 is not within the 95% quantile of the data, meaning significant differences for this comparison. All comparisons in this case are significant. B: Boxplot comparing net diversification rate of the two hidden states. In this case, the upper and lower boundary of the box is the default value, which is the 75th percentile and 25th percentile respectively. Letter above the box indicating group, with different letters indicating significant differences.

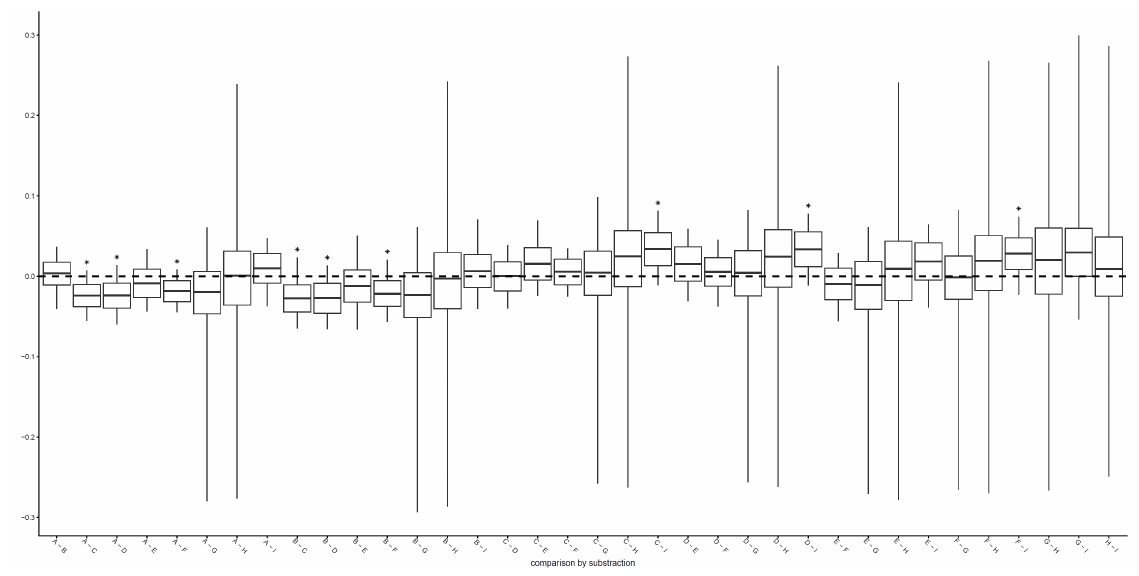

**A**

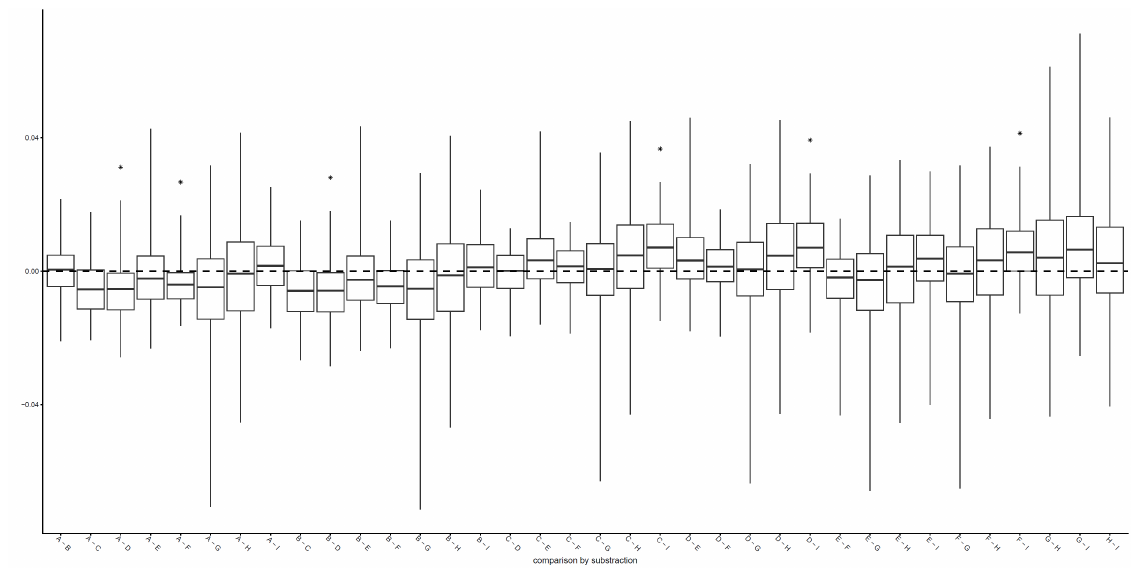

**B**

Figure S15. Boxplot showing comparison of net diversification rate (speciation rate – extinction rate) of all observed states within each hidden state by subtracting the two rates in each iteration. A: comparison among observed states within hidden state 0. B: comparison among observed states within hidden state 1. In this case, the lower and upper boundaries of the boxplot box are 2.5% and 97.5% quantile respectively. Number 0 is highlighted with a thick dash line. Asterisks above the box mean number 0 is not within the 95% quantile of the data, meaning significant differences for this comparison. State coding: A-Africa-Arabia; B-Madagascar; C-Eurasia; D-South Asia; E-Sundaland; F-Australasia; G-Carribean; H-North-Central America; I-South America.

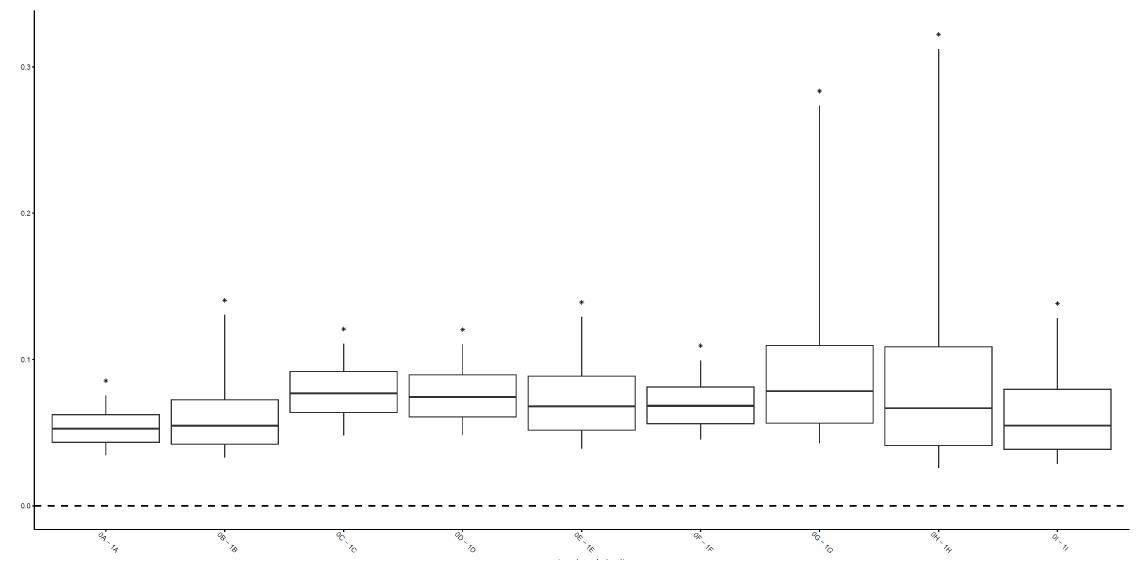

**A**

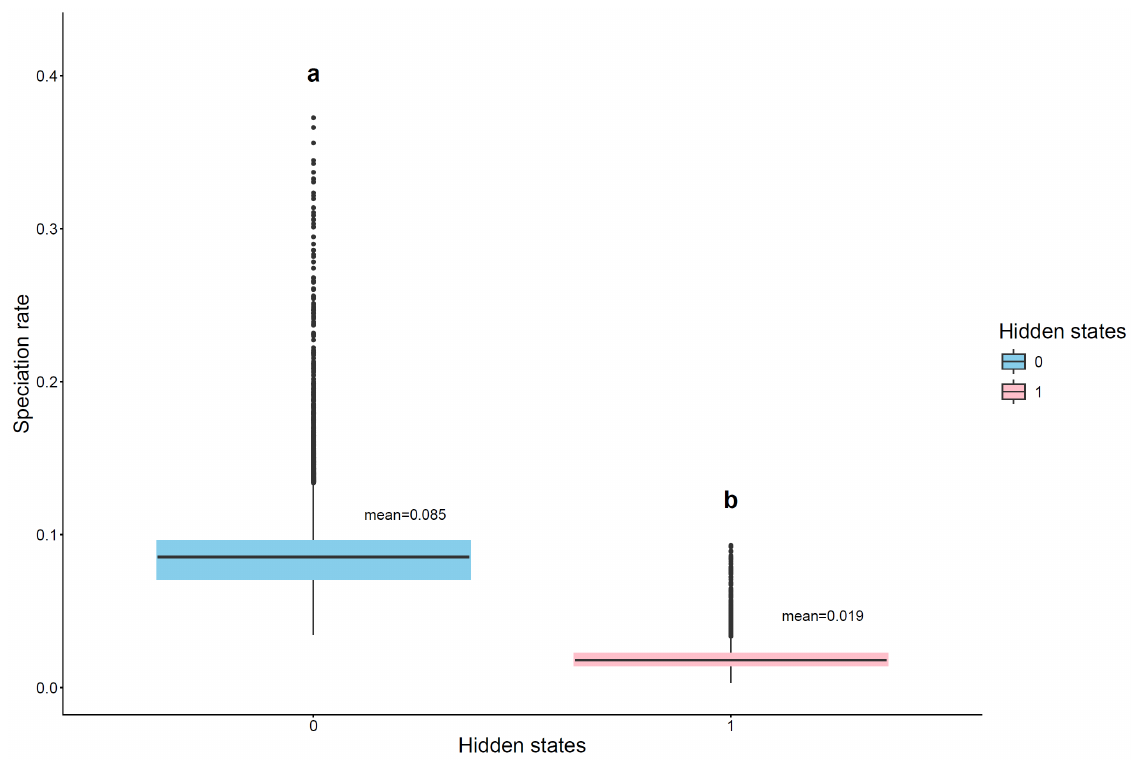

**B**

Figure S16. Boxplot showing posterior estimates of the speciation rate for hidden states 0 and 1. A: Boxplot showing stats of subtracting rate of the same observed states with different hidden states in each iteration, e.g. 0A – 1A, 0B - 1B, etc.. In this case, the lower and upper boundaries of the boxplot box are 2.5% and 97.5% quantile respectively. Number 0 is highlighted with a thick dash line. Asterisks above the box mean number 0 is not within the 95% quantile of the data, meaning significant differences for this comparison. All comparisons in this case are significant. B: Boxplot comparing speciation rate of the two hidden states. In this case, the upper and lower boundary of the box is the default value, which is the 75th percentile and 25th percentile respectively. Letter above the box indicating group, with different letters indicating significant differences.

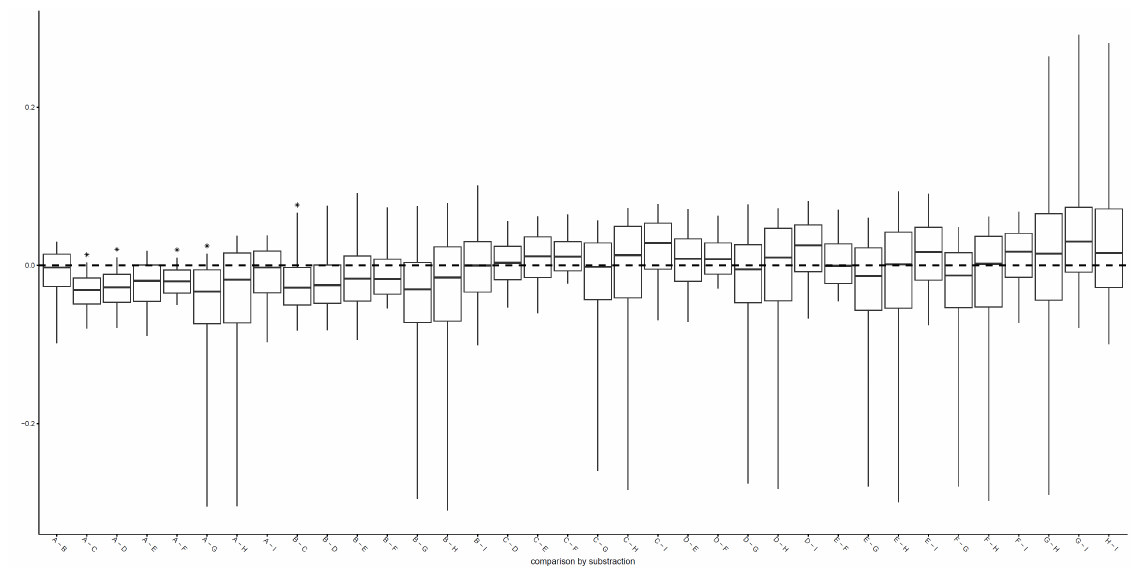

**B**

**A**

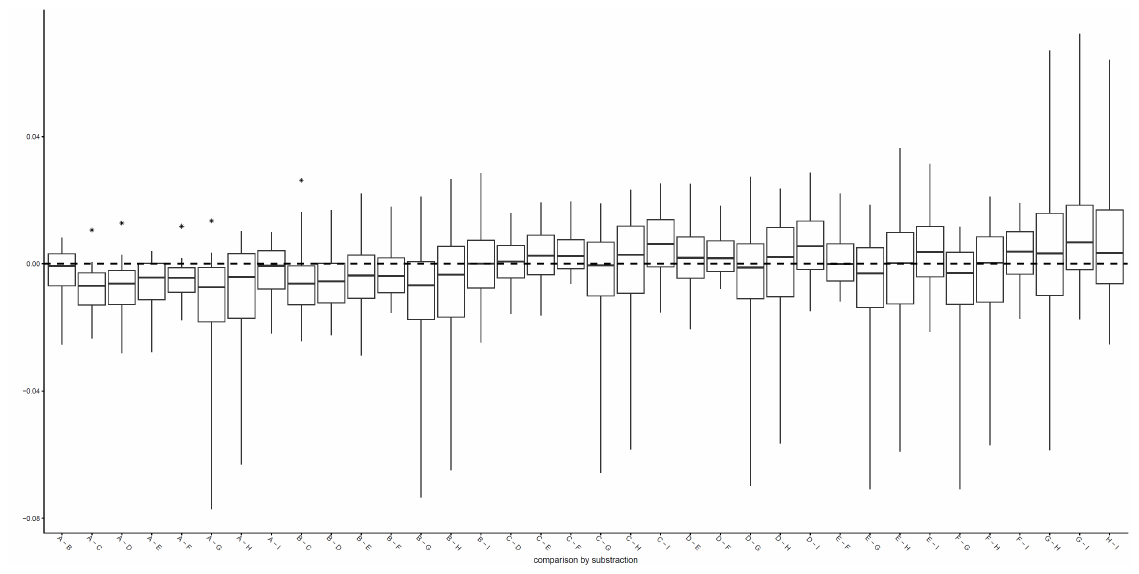

Figure S17. Boxplot showing comparison of speciation rate of all observed states within each hidden state by subtracting the two rates in each iteration. A: comparison among observed states within hidden state 0. B: comparison among observed states within hidden state 1. In this case, the lower and upper boundaries of the boxplot box are 2.5% and 97.5% quantile respectively. Number 0 is highlighted with a thick dash line. Asterisks above the box mean number 0 is not within the 95% quantile of the data, meaning significant differences for this comparison. State coding: A-Africa-Arabia; B-Madagascar; C-Eurasia; D-South Asia; E-Sundaland; F-Australasia; G-Carribean; H-North-Central America; I-South America.

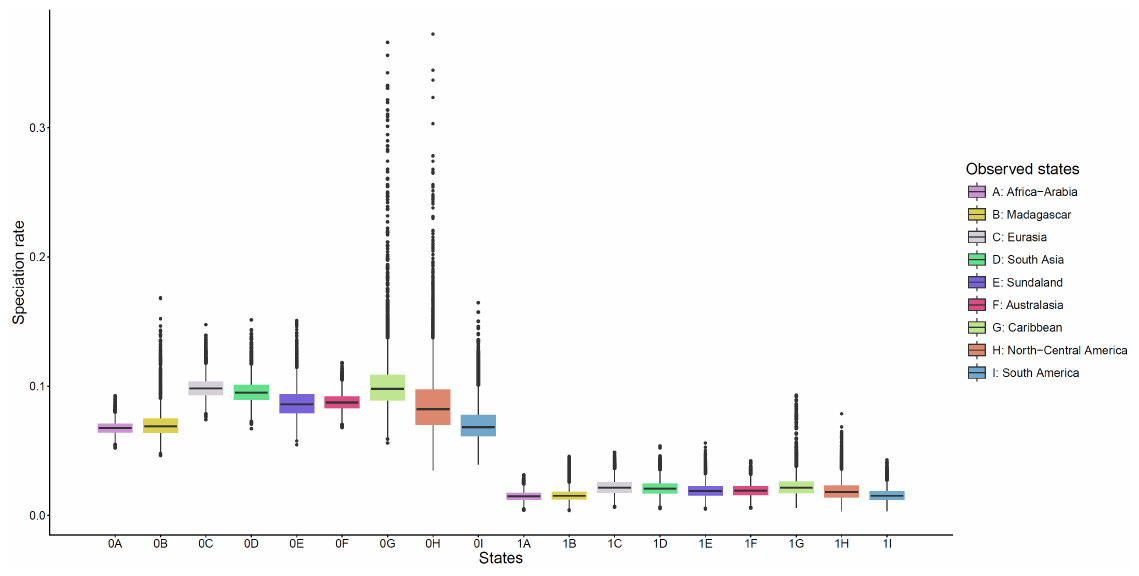

**A**

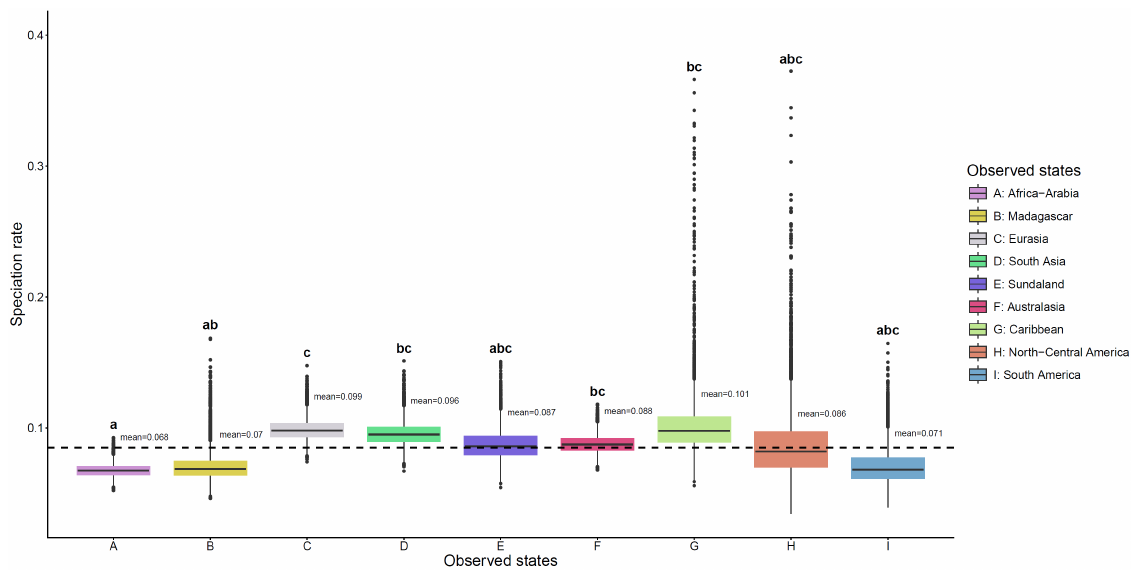

**B**

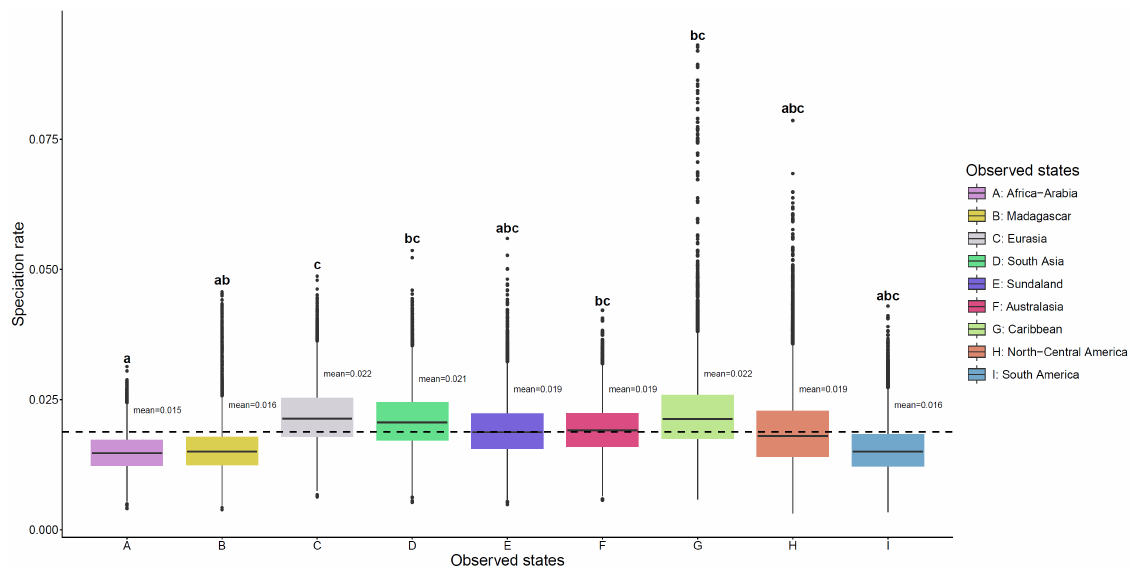

**C**

Figure S18. Boxplot showing posterior estimates of the speciation rate from MuHisse model with two hidden states. A: boxplot of all observed state and hidden state combinations together showing that species in hidden state 0 have higher speciation rate than species in hidden state 1. This difference is significant (see also Figure S13). B & C: boxplot of speciation rate estimates within hidden state 0 and 1. Letter above the box indicating group, with different letters indicating significant differences (see also Figure S14). The thick dash line indicates the mean rate of this hidden state. Observed state coding: A-Africa-Arabia; B-Madagascar; C-Eurasia; D-South Asia; E-Sundaland; F-Australasia; G-Carribean; H-North-Central America; I-South America.

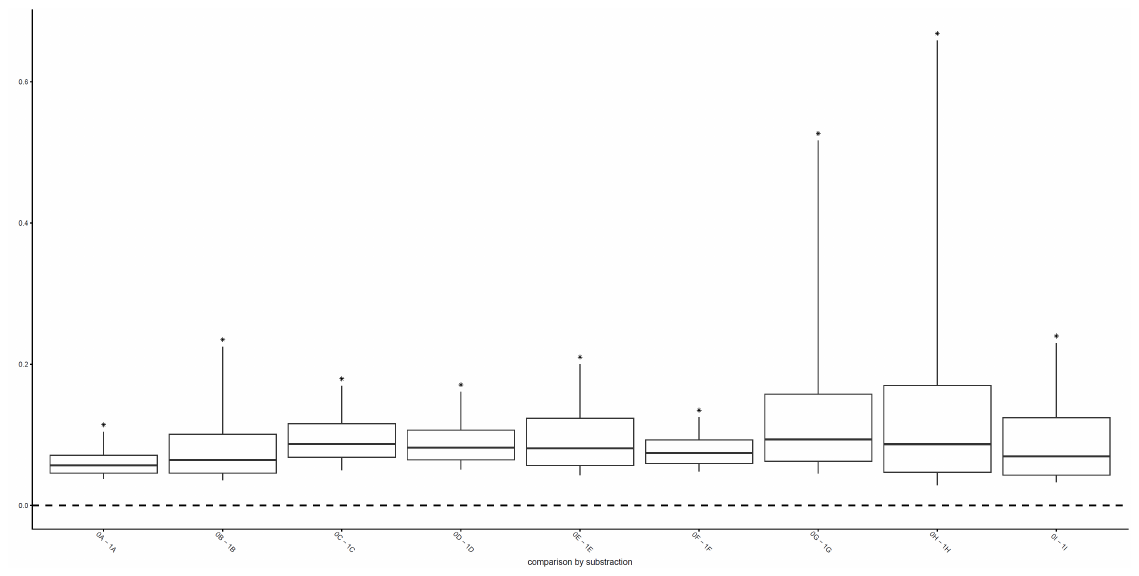

**A**

**B**

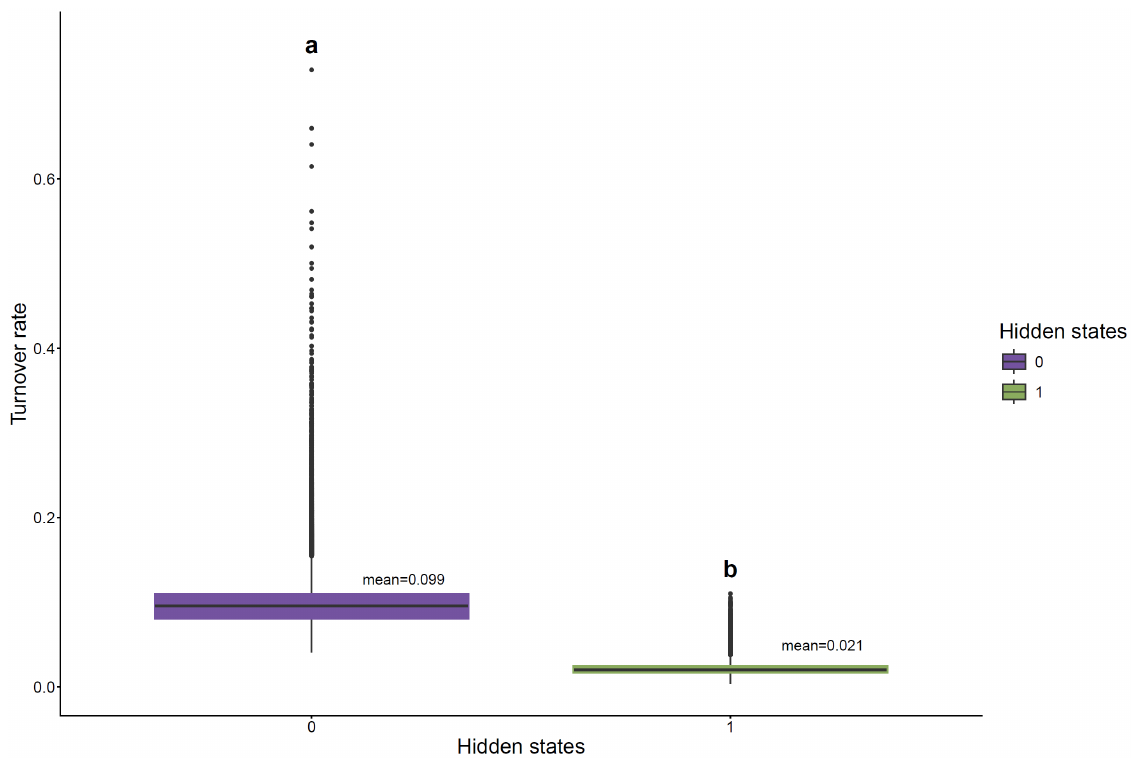

Figure S19. Boxplot showing posterior estimates of the turnover rate (speciation rate + extinction rate) for hidden states 0 and 1. A: Boxplot showing stats of subtracting rate of the same observed states with different hidden states in each iteration, e.g. 0A – 1A, 0B - 1B, etc.. In this case, the lower and upper boundaries of the boxplot box are 2.5% and 97.5% quantile respectively. Number 0 is highlighted with a thick dash line. Asterisks above the box mean number 0 is not within the 95% quantile of the data, meaning significant differences for this comparison. All comparisons in this case are significant. B: Boxplot comparing speciation rate of the two hidden states. In this case, the upper and lower boundary of the box is the default value, which is the 75th percentile and 25th percentile respectively. Letter above the box indicating group, with different letters indicating significant differences.

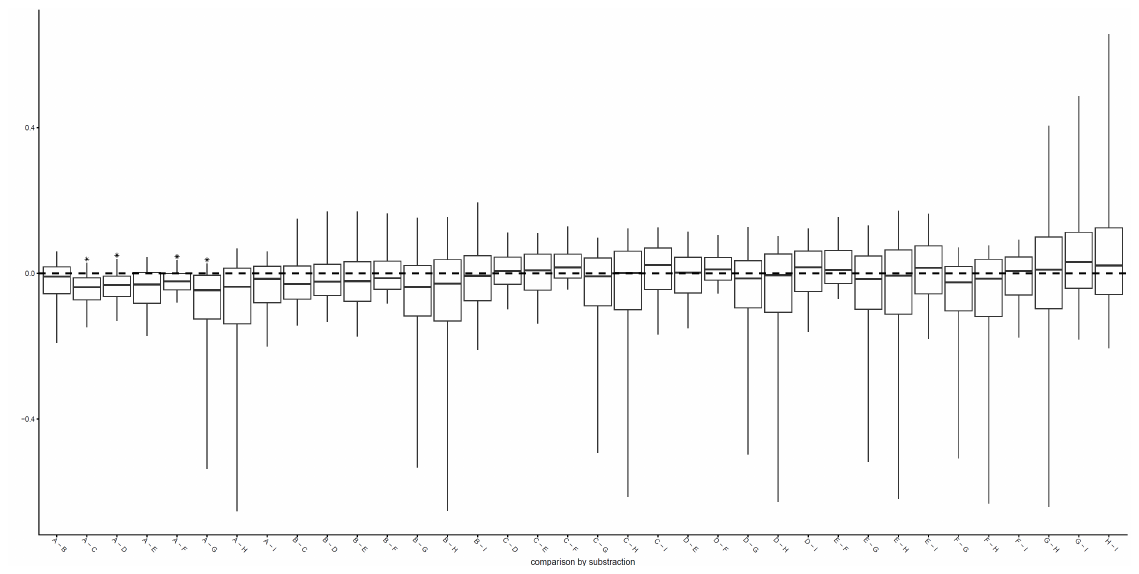

**A**

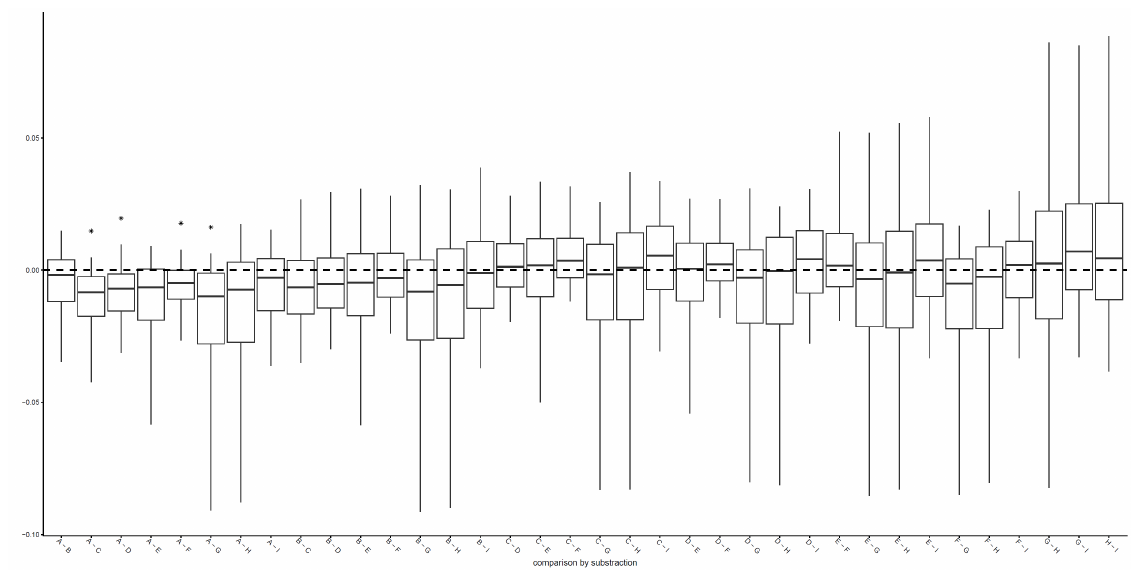

**B**

Figure S20. Boxplot showing comparison of turnover rate (speciation rate + extinction rate) of all observed states within each hidden state by subtracting the two rates in each iteration. A: comparison among observed states within hidden state 0. B: comparison among observed states within hidden state 1. In this case, the lower and upper boundaries of the boxplot box are 2.5% and 97.5% quantile respectively. Number 0 is highlighted with a thick dash line. Asterisks above the box mean number 0 is not within the 95% quantile of the data, meaning significant differences for this comparison. State coding: A-Africa-Arabia; B-Madagascar; C-Eurasia; D-South Asia; E-Sundaland; F-Australasia; G-Carribean; H-North-Central America; I-South America.

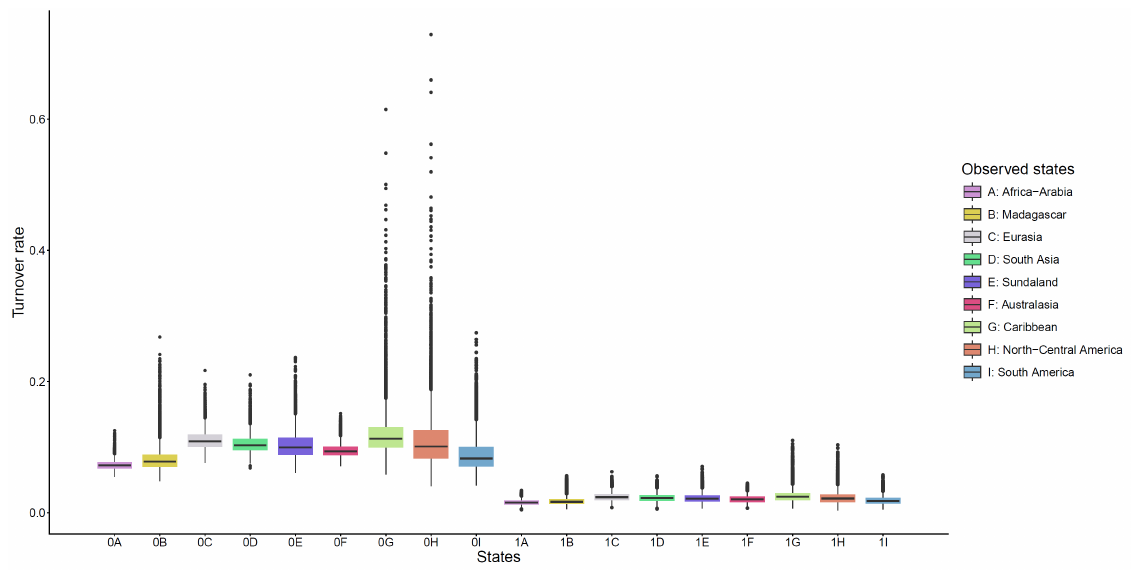

**A**

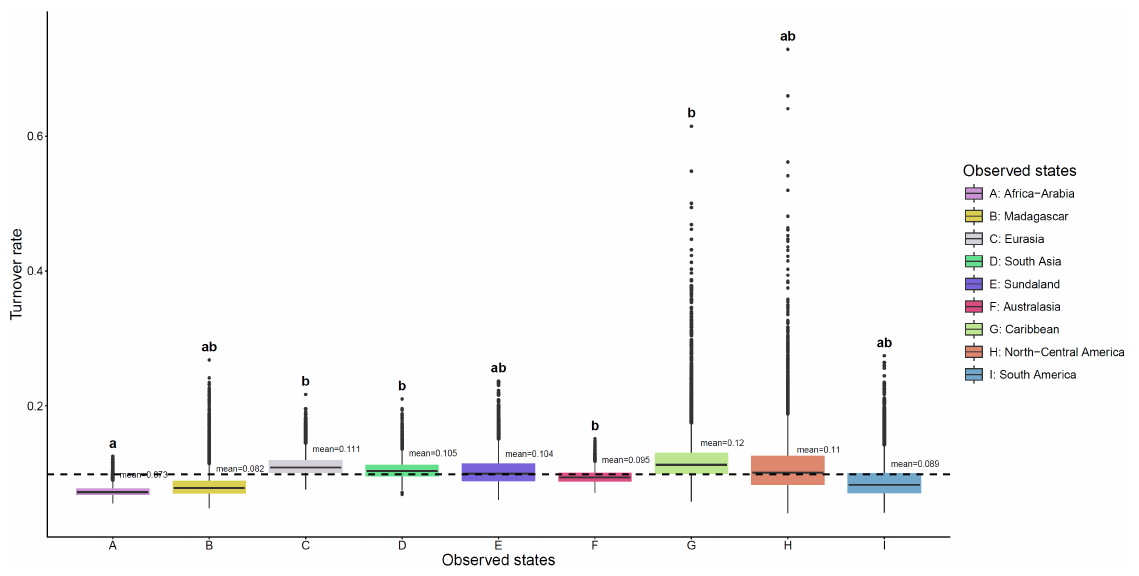

**B**

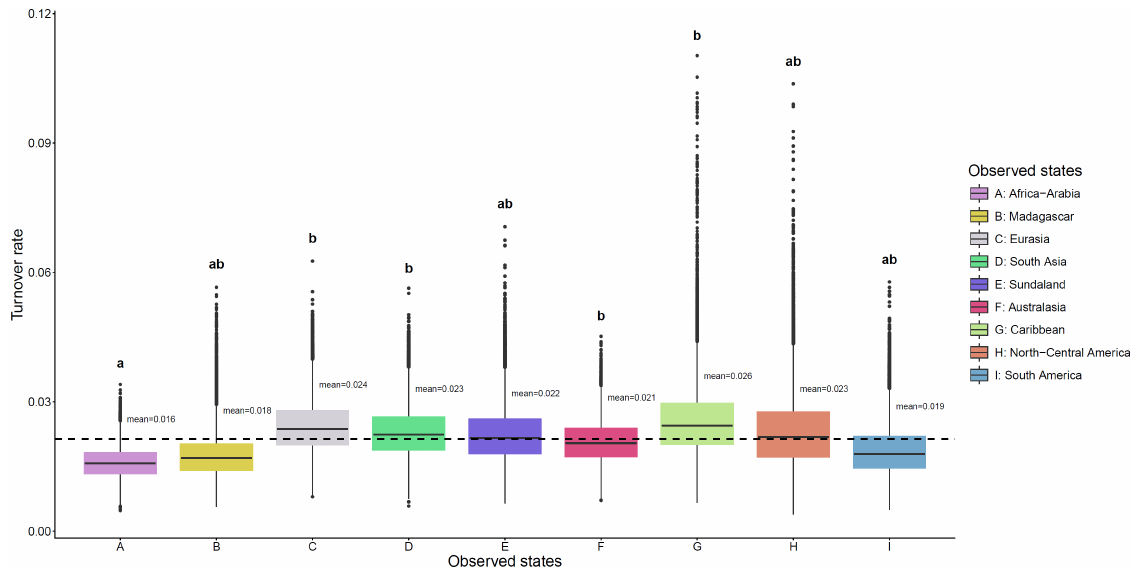

**C**

Figure S21. Boxplot showing posterior estimates of the turnover rate (speciation rate + extinction rate) from MuHisse model with two hidden states. A: boxplot of all observed state and hidden state combinations together showing that species in hidden state 0 have higher speciation rate than species in hidden state 1. This difference is significant (see also Figure S18). B & C: boxplot of turnover rate estimates within hidden state 0 and 1. Letter above the box indicating group, with different letters indicating significant differences (see also Figure S19). The thick dash line indicates the mean rate of this hidden state. Observed state coding: A-Africa-Arabia; B-Madagascar; C-Eurasia; D-South Asia; E-Sundaland; F-Australasia; G-Carribean; H-North-Central America; I-South America.

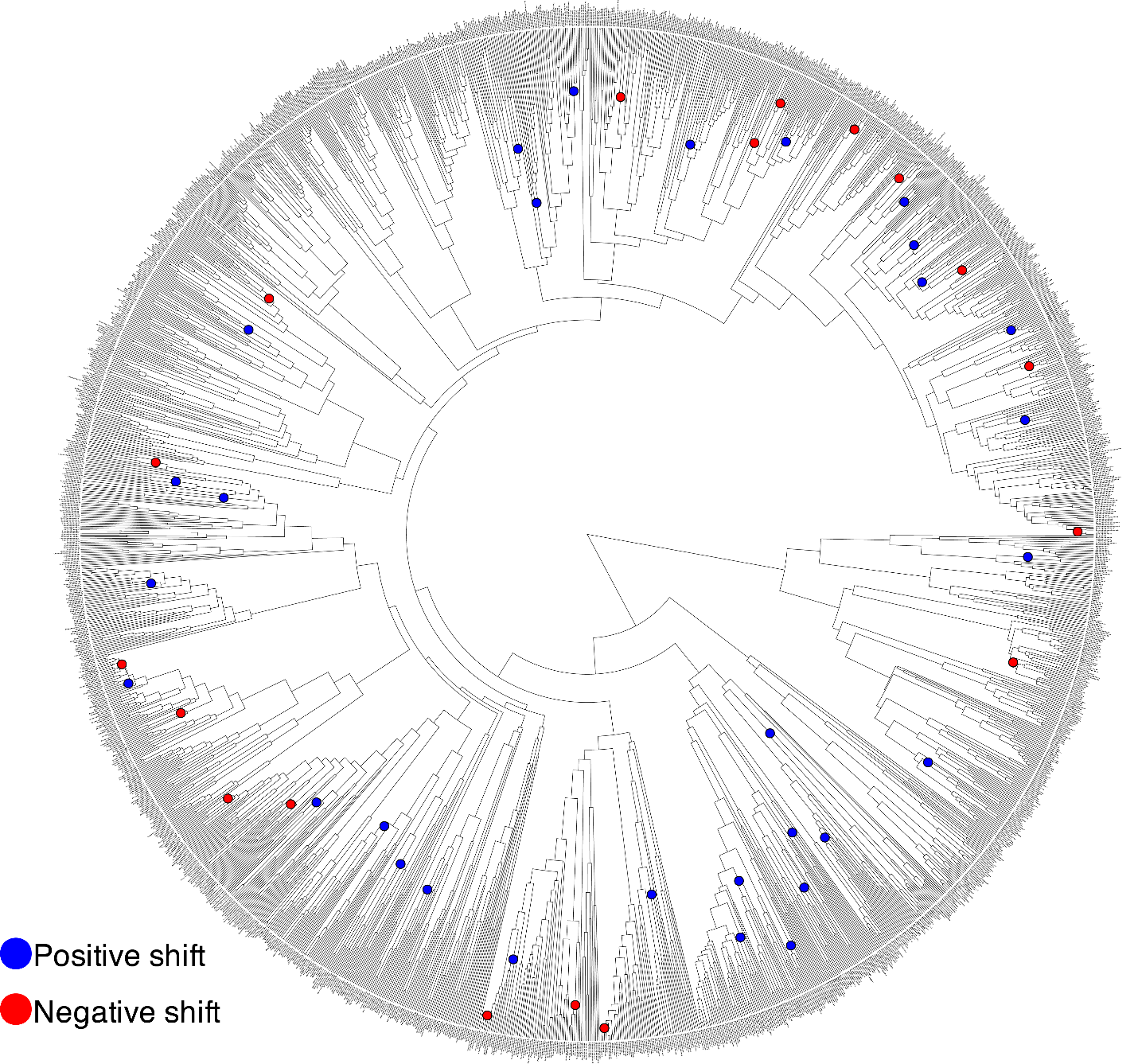

Figure S22. Speciation rates across geckos estimated by ClaDS on time-calibrated tree (UCE-supermatrix MCC) displaying all 1,783 tips. Shifts in rate are indicated by red (above background rate) and blue (below background rate) as estimated by Rrphylo.
